# Genomic evidence for divergent co-infections of SARS-CoV-2 lineages

**DOI:** 10.1101/2021.09.03.458951

**Authors:** Hang-Yu Zhou, Ye-Xiao Cheng, Lin Xu, Jia-Ying Li, Chen-Yue Tao, Cheng-Yang Ji, Na Han, Rong Yang, Yaling Li, Aiping Wu

## Abstract

Recently, patients co-infected by two SARS-CoV-2 lineages have been sporadically reported. Concerns are raised because previous studies have demonstrated co-infection may contribute to the recombination of RNA viruses and cause severe clinic symptoms. In this study, we have estimated the compositional lineage(s), tendentiousness, and frequency of co-infection events in population from a large-scale genomic analysis for SARS-CoV-2 patients. SARS-CoV-2 lineage(s) infected in each sample have been recognized from the assignment of within-host site variations into lineage-defined feature variations by introducing a hypergeometric distribution method. Of all the 29,993 samples, 53 (~0.18%) co-infection events have been identified. Apart from 52 co-infections with two SARS-CoV-2 lineages, one sample with co-infections of three SARS-CoV-2 lineages was firstly identified. As expected, the co-infection events mainly happened in the regions where have co-existed more than two dominant SARS-CoV-2 lineages. However, co-infection of two sub-lineages in Delta lineage were detected as well. Our results provide a useful reference framework for the high throughput detecting of SARS-CoV-2 co-infection events in the Next Generation Sequencing (NGS) data. Although low in average rate, the co-infection events showed an increasing tendency with the increased diversity of SARS-CoV-2. And considering the large base of SARS-CoV-2 infections globally, co-infected patients would be a nonnegligible population. Thus, more clinical research is urgently needed on these patients.

## Introduction

Since its initial report in the late of 2019, severe acute respiratory syndrome coronavirus 2 (SARS-CoV-2) has rapidly developed into a global pandemic^1,2^. The widespread transmission and geographical isolation of SARS-CoV-2 has greatly resulted in its genetic diversity. Until Jul. 18^th^, 2021, thousands of lineages have been clearly defined by the Pangolin nomenclature^3^. Viruses within a defined lineage often share several common mutations and have similar biological properties. Of all the identified lineages, four SARS-CoV-2 variants have been announced as “a variant of concern (VoC)” by the World Health Organization (WHO). Among them, B.1.1.7 (Alpha strain defined by WHO) was estimated to have >50% enhanced transmissibility^4^, B.1.351 (Beta strain defined by WHO) and P.1 (Gamma strain defined by WHO) showed the capacity to evade inhibition by neutralizing antibodies^5^, while B.1.617.2 (Delta strain defined by WHO) caused the greatly increased infections in India and became the dominant epidemic strain in some countries recently^6,7^.

Currently, the re-infection of SARS-CoV-2 have been extensively discussed^8,9^. Meanwhile, more and more evidence has pointed out that the co-infection events caused by different SARS-CoV-2 lineages may have occasionally occured^10–14^. This phenomenon should be paid more attention for at least two reasons. Firstly, previous reports indicated that viral co-infection may cause severe clinic symptoms. For instance, human immunodeficiency virus (HIV) co-infection contributes to rapid disease progression^15^, increased viral load^16^, and requires antiretroviral treatment effective against both HIV variants^17^. Secondly, the co-infection may contribute to the SARS-CoV-2 recombination and accelerate the generation of recombinant viruses, since coronaviruses have relatively high recombination rates^18–20^. Thus, with the increasing diversity of SARS-CoV-2 and the co-existence of multiple regional lineages globally, it is significant to clarify that how often the co-infection occurs in population and what’s the exact compositional lineages of co-infection in individuals.

In theory, genomic evidence should be kept in the deep sequencing data if a patient has been co-infected with two or even more SARS-CoV-2 lineages. Like other RNA viruses, the identified SARS-CoV-2 genomes in patient belongs to quasi-species with many within-host variations^12,21,22^. The identification of co-infection lineages mainly rely on the existence of their lineage-defined feature mutations in viral quasi-species. Recent studies have confirmed the high reliability of illumina sequencing data in detecting within-host variations^23^, and deep sequencing data generated by illumina could be used for formal analysis^24,25^. Benefiting from the worldwide rapid accumulation and open sharing of SARS-CoV-2 genomes, the large-scale genomic dataset provides us substantial support to detect the co-infection events even when they are very rare in population.

In this study, we have collected and analyzed 29,993 paired-end deep sequencing genomes generated by illumina platform from the National Center for Biotechnology Information (NCBI). All these samples had detailed metadata and were isolated from the USA during 1^st^ Jan. 2021 to 31^st^ Jul. 2021. By introducing a hypergeometric-distribution based method, we have successfully decoded the within-host variations from these deep sequencing data and detected the compositional lineages of potential co-infection events. Of all the 29,993 samples, we have identified 53 (~0.18%) co-infection events of SARS-CoV-2. Apart from 52 samples with two co-infected lineages, one sample was co-infected with three lineages. As expected, the co-infection events have mainly happened in the region where co-circulated with two or even more dominant SARS-CoV-2 lineages.

Overall, for the first time, we have captured the genomic evidence for the co-infection events in large-scale sequencing data and provided robust method to identify the compositional lineages of SARS-CoV-2 in co-infected samples. Furthermore, the increased number of co-infected samples might also raise great concerns for the possible viral recombination and decrease of vaccine effectiveness.

## Results

When an individual has been infected by SARS-CoV-2 virus from a single lineage, we can imagine that most of the lineage-defined feature variations could be detected at a similar level in deep sequencing data, and the frequency of each lineage-defined feature variation should be nearly 100% (Fig. 1). However, when an individual has been co-infected by two strains from different SARS-CoV-2 lineages, the situation is a little complicated. Assuming that the co-infected strains from two lineages are propagated independently in a patient, then three evidence could be observed in the genomic sequencing data (Fig. 1). Firstly, feature variations specific to the same lineage A or B possess the similar frequencies. Secondly, the sum of the mean frequencies of feature variations specific to lineages A and B is nearly 100%. Thirdly, the frequencies of the shared feature variations by lineages A and B are nearly 100% (Fig. 1). Based on these three evidence, co-infection event could possibly be identified from the genomic sequencing data.

**Fig. 1.**
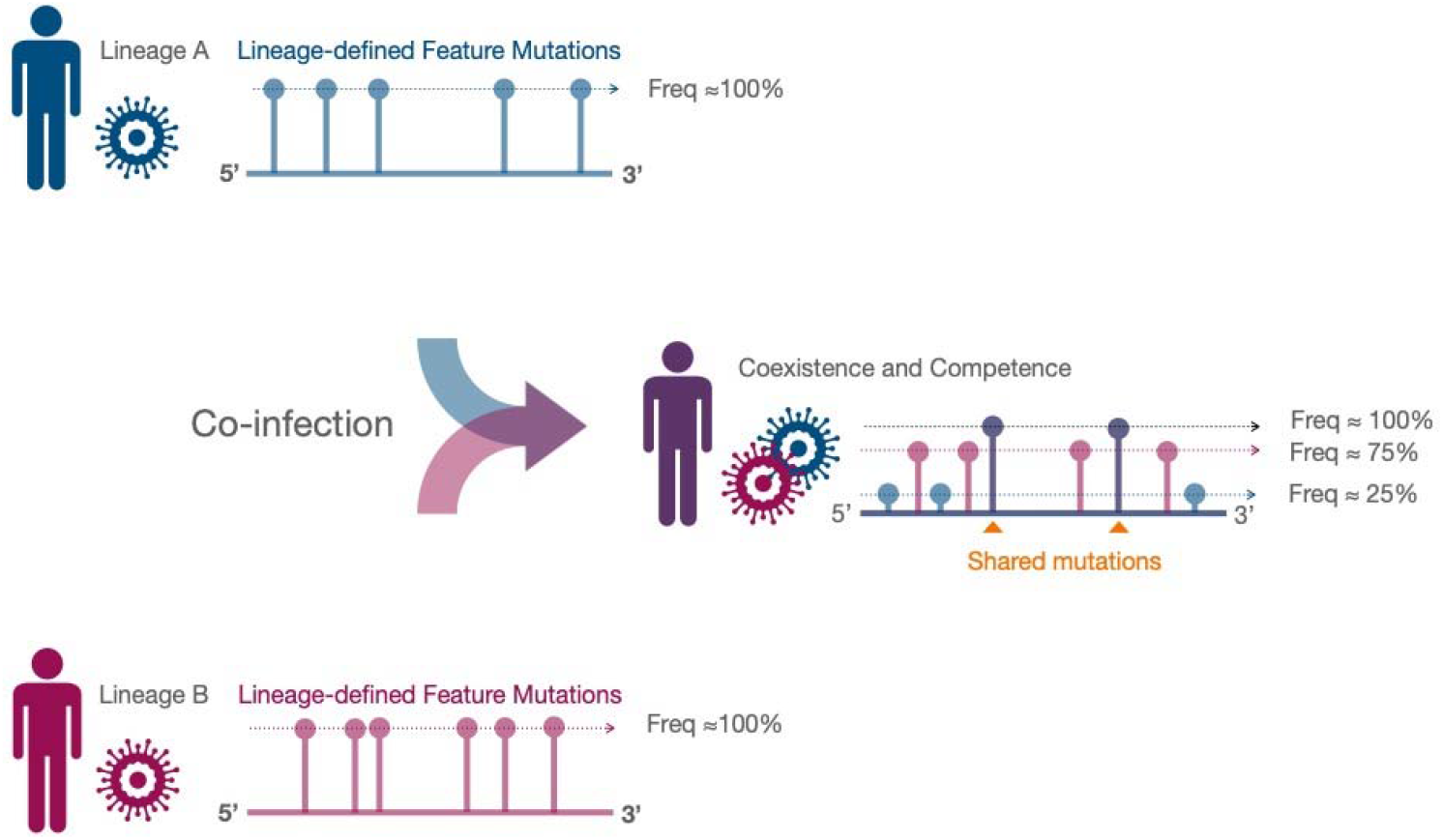
The co-infection pattern of SARS-CoV-2 lineages in patient. In one lineage infection, frequencies of lineage-defined feature variations are ~100%. When a sample was co-infected by two SARS-CoV-2 lineages A and B, the frequencies of each lineage-specific feature variations (blue and pink) have their own levels. The sum of the average frequencies of lineage-specific feature variations is ~100%, and the shared feature variations of lineages A and B (purple) are ~100%.

In this study, we have introduced a hypergeometric-distribution based method to decode the possible lineages in a sequencing sample (Fig. 2 and Methods). Of all the 29,993 samples, 28,627 (95.45%) were concluded to be infected by only one SARS-CoV-2 lineage. As was shown in Fig. 3A, the pattern of feature variations for a typical one-lineage infection is easy to be determined. Namely, most of the feature variations belong to a specific lineage could be detected in the sample. Besides, the feature variations have similar frequency of reads in each site, demonstrating the good genomic homogeneity within a single lineage. In addition, few variations that do not match any lineage-defined feature variations were observed and could be recognized as *de novo* mutations. Furthermore, in this case, the identified Alpha (B.1.1.7) lineage is exactly the dominant lineage in the place where the sample was collected (Fig. 3B), which confirmed the rationality of identifying lineage with its lineage-defined feature variations from deep sequencing data.

**Fig. 2.**
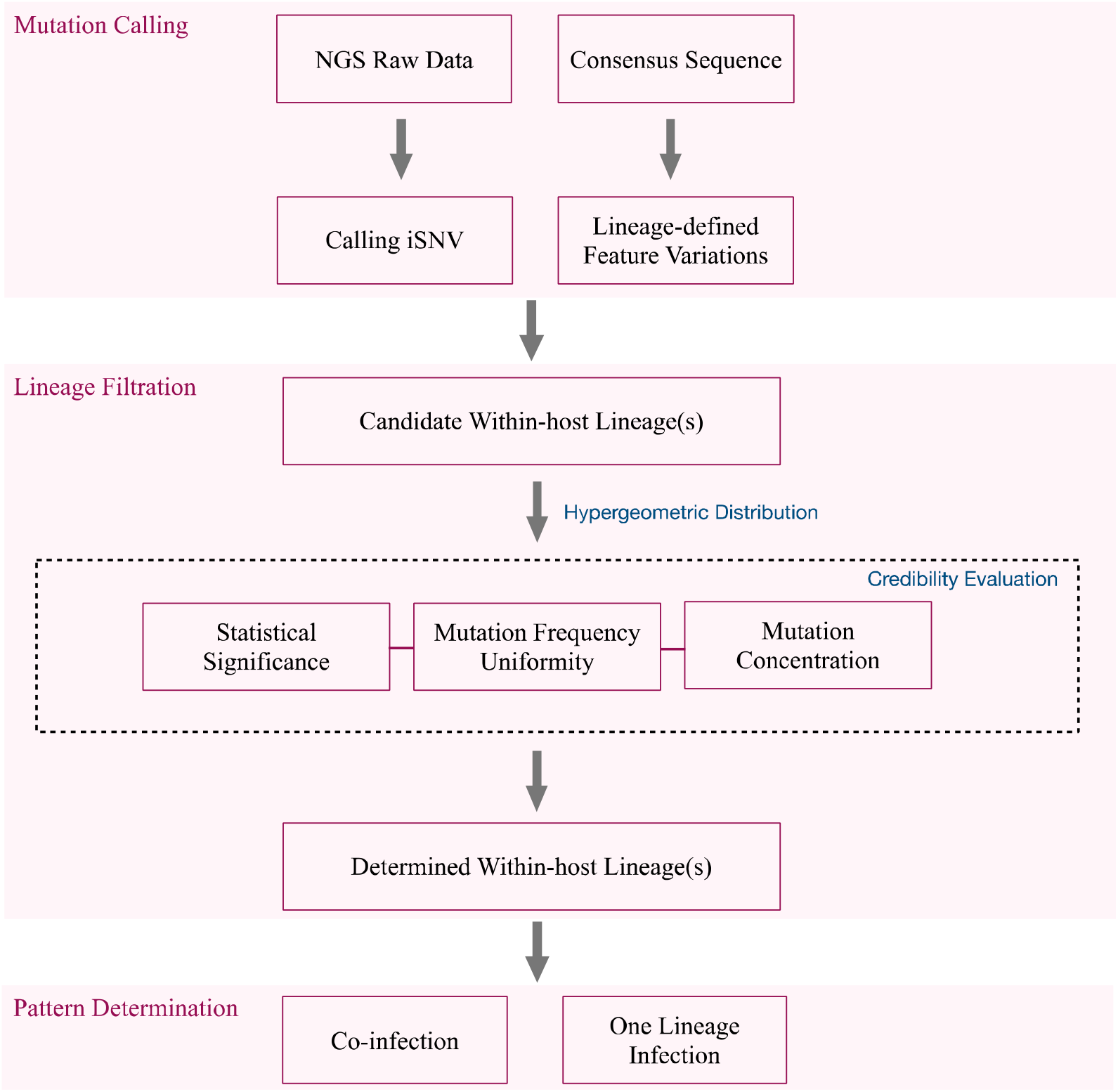
Workflow of detecting within-host SARS-CoV-2 lineage(s). Three modules are developed for identifying the SARS-CoV-2 lineage(s) in each sample, including mutation calling, lineage filtration and pattern determination. In mutation calling module, next-generation sequencing (NGS) raw data (FASTQ file) was imported into CLC genomics workbench for SNV calling. All available consensus sequences in GISAID were collected and identified for feature variations. In lineage filtration module, two files from the mutation calling module were put into a homemade Python script for hypergeometric distribution analysis. The candidate within-host lineages were identified for the next analysis. By ranking candidate lineages according to the statistical significance (p-value), mutation frequency uniformity (Frequency STD) and mutation proportion (P_mutation_/N_mutation_), lineage with the highest confidence was recorded. In the last module for pattern determination, sample with one identified lineage were classified as One lineage infection sample, samples with at least two identified SARS-CoV-2 lineages were classified as Co-infection samples, the remaining samples that cannot be classified into any type were classified as Unclassified.

**Fig. 3.**
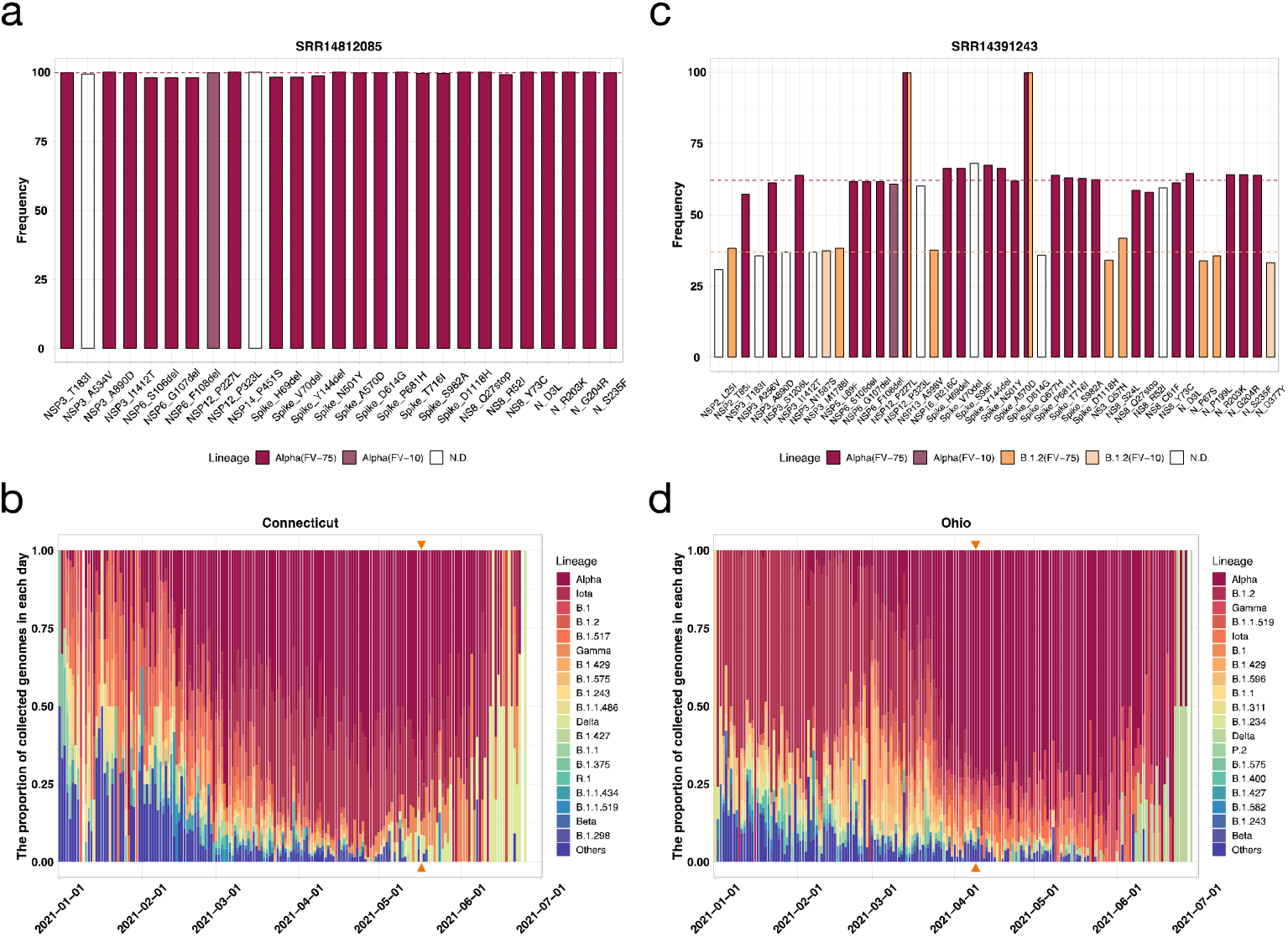
Patterns for one lineage infection and two lineages co-infection. **a** A sample infected by one specific SARS-CoV-2 lineage. Most of the feature variations of the identified Alpha lineage (FV-75 and FV-10) could be detected at the same level. Non-determined variations are shown as white column. **b** The lineage ratio of SARS-CoV-2 lineages isolated in Connecticut State from 1^st^ Jan. 2021 to 1^st^ Jul 2021, covering the location and timepoint of the representative sample used in **a**, that is, Connecticut state and 17^th^ May 2021 (the date was signed with orange arrows). **c** A sample co-infected by two SARS-CoV-2 lineages. Most of the feature variations of two identified lineages (Alpha and B.1.2) were shown in purple and orange. Two shared variations were shown as both purple and orange. **d** The lineage ratio of SARS-CoV-2 lineages isolated in Ohio State from 1^st^ Jan. 2021 to 1^st^ Jul 2021. The sample used in **c** was isolated in Ohio state at 9^th^ Apr. 2021.

In total, 52 samples were clearly classified to be co-infected by SARS-CoV-2 strains from two different lineages (Figs. 3C and S1). As shown in Figure 3C, Alpha (B.1.1.7) and B.1.2 were identified as the two lineages in this co-infected sample. Over 87.5% (21 of 24) FV-75 feature variations of Alpha lineage and 72.7% (8 of 11) FV-75 feature variations of B.1.2 lineage have been detected in this sample, respectively. Meanwhile, the average frequency of Alpha lineage-specific variations is ~65%, while that of B.1.2 lineage is ~35%, and the average frequencies of the two shared variations by Alpha and B.1.2 lineages, including NSP12_P323L and Spike_D614G, were nearly 100%. These observed facts exactly matched with the three hypothesized genomic evidence proposed in Figure 1. The co-infection of Alpha and B.1.2 lineages was also consistent with the epidemiological background of regional SARS-CoV-2. As shown in Fig. 3D, at the collection date (9^th^ Apr. 2021), the Alpha lineage is the dominant lineage in Ohio State, while B.1.2 is the second dominant epidemic lineage.

Apart from co-infection by two lineages, we have unexpectedly identified one co-infected sample with three lineages. As shown in Figure 4, the sample was collected in Connecticut State, USA on 17^th^ May 2021. The three hypothesized genomic evidence (Fig. 1) could be observed in this sample clearly (Fig. 4A). Firstly, most lineage-specific feature variations of Alpha, Iota (B.1.526) and Gamma (P.1) could be identified at their own levels, respectively. Alpha lineage was identified to occupy ~55% of all strains while Iota and Gamma occupied ~25% and ~15%, respectively. Secondly, three feature variations (Spike_N501Y, N_R203K and N_G204R) shared by Gamma and Alpha were nearly 70%, which almost equal to the sum of the mean frequencies of Alpha and Gamma. Thirdly, two feature variations (NSP12_P323L and Spike_D614G) shared by all three lineages were nearly 100%. The detection of these three lineages were also in consistent with the epidemiological patterns of SARS-CoV-2 lineages in the sampling location, Connecticut State (Fig. 4B).

**Fig. 4.**
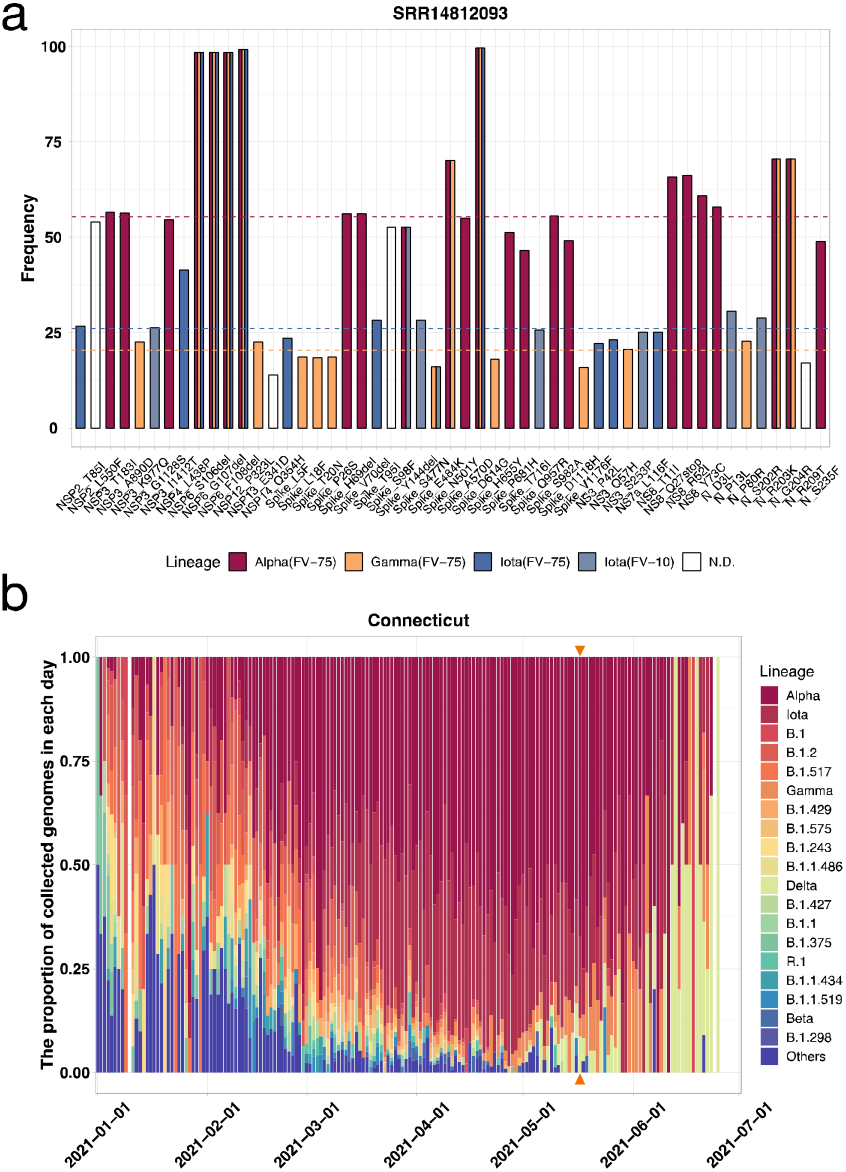
Co-infection of three SARS-CoV-2 lineages. **a** An identified co-infected sample by three SARS-CoV-2 lineages. The feature variations of the three identified lineages (Alpha, Gamma, and Iota) are shown as purple, orange, and blue, respectively. **b** The lineage ratio of SARS-CoV-2 lineages isolated in Connecticut State from 1^st^ Jan. 2021 to 1^st^ Jul. 2021. The sample used in **a** was isolated in Connecticut State on 17^th^ May. 2021 (The day was denoted by orange arrows).

The metadata of all the SARS-CoV-2 co-infected samples (Table 1) were further analyzed. Of all the 53 co-infected samples, 27 are male and 26 are female which indicated no gender bias. The average age is 34 years old (the median is 29 years old) for all 53 patients, in which the youngest patient is nine years old, and the oldest patient is 71 years old. No obvious spatial-temporal bias was found in these samples. For the viral load, we evaluated it with the diagnostic PCR cycle threshold (Ct) value. Compared to the average Ct value of all samples (18.72), the average Ct value of co-infected samples was at similar level as 19.96. Additionally, in these samples, we have found some samples with false identified lineage by NCBI. For instance, a sample has been wrongly classified as B.1 infection (Fig. S2), while we found all the feature variations of it belong to two identified lineages (Alpha and Iota) as ~50%, respectively.

**Table 1:**
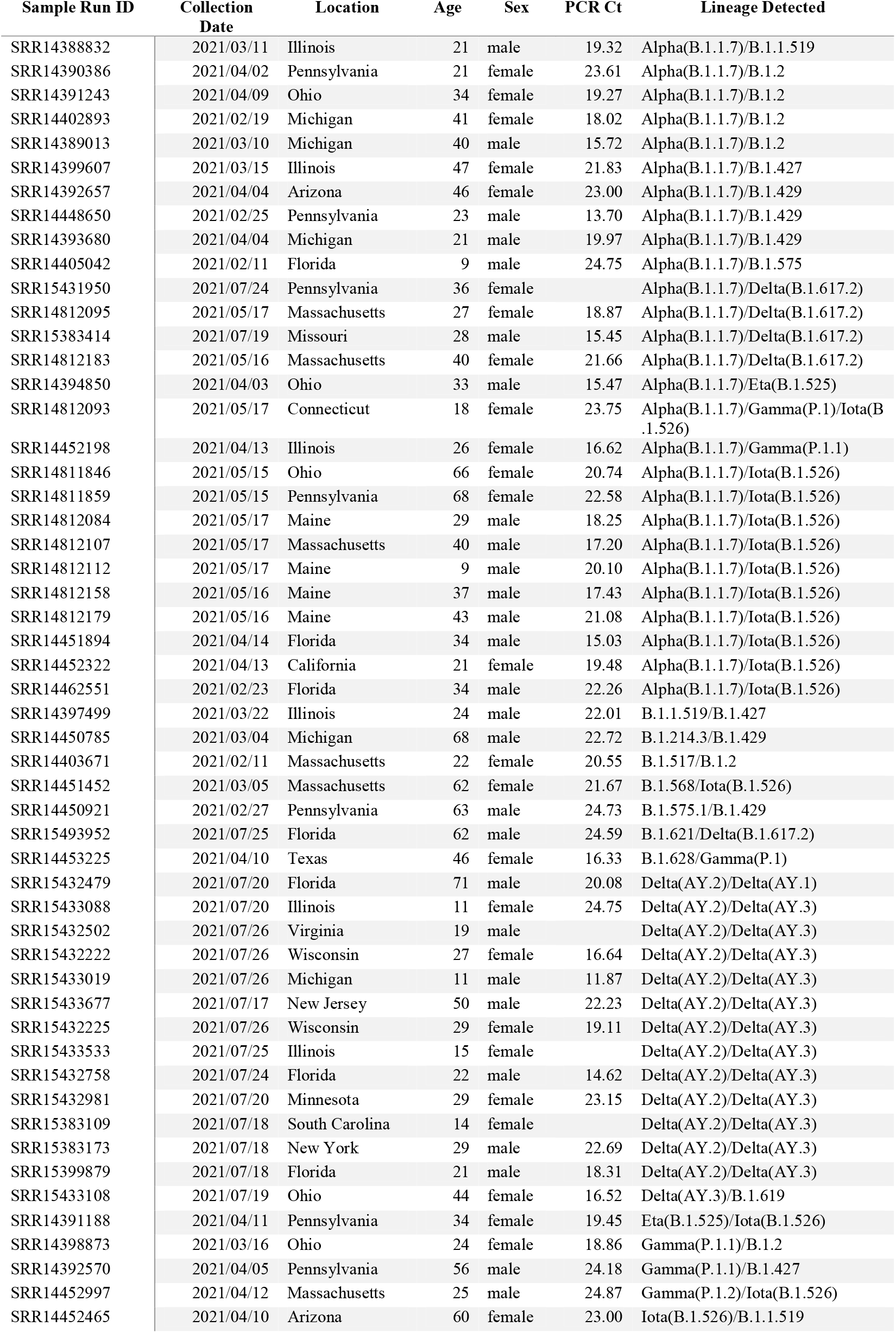
The metadata of all the SARS-CoV-2 co-infected samples.

| Sample Run ID | Collection Date | Location | Age | Sex | PCR Ct | Lineage Detected |
| --- | --- | --- | --- | --- | --- | --- |
| SRR14388832 | 2021/03/11 | Illinois | 21 | male | 19.32 | Alpha(B.1.1.7)/B.1.1.519 |
| SRR14390386 | 2021/04/02 | Pennsylvania | 21 | female | 23.61 | Alpha(B.1.1.7)/B.1.2 |
| SRR14391243 | 2021/04/09 | Ohio | 34 | female | 19.27 | Alpha(B.1.1.7)/B.1.2 |
| SRR14402893 | 2021/02/19 | Michigan | 41 | female | 18.02 | Alpha(B.1.1.7)/B.1.2 |
| SRR14389013 | 2021/03/10 | Michigan | 40 | male | 15.72 | Alpha(B.1.1.7)/B.1.2 |
| SRR14399607 | 2021/03/15 | Illinois | 47 | female | 21.83 | Alpha(B.1.1.7)/B.1.427 |
| SRR14392657 | 2021/04/04 | Arizona | 46 | female | 23.00 | Alpha(B.1.1.7)/B.1.429 |
| SRR14448650 | 2021/02/25 | Pennsylvania | 23 | male | 13.70 | Alpha(B.1.1.7)/B.1.429 |
| SRR14393680 | 2021/04/04 | Michigan | 21 | male | 19.97 | Alpha(B.1.1.7)/B.1.429 |
| SRR14405042 | 2021/02/11 | Florida | 9 | male | 24.75 | Alpha(B.1.1.7)/B.1.575 |
| SRR15431950 | 2021/07/24 | Pennsylvania | 36 | female |  | Alpha(B.1.1.7)/Delta(B.1.617.2) |
| SRR14812095 | 2021/05/17 | Massachusetts | 27 | female | 18.87 | Alpha(B.1.1.7)/Delta(B.1.617.2) |
| SRR15383414 | 2021/07/19 | Missouri | 28 | male | 15.45 | Alpha(B.1.1.7)/Delta(B.1.617.2) |
| SRR14812183 | 2021/05/16 | Massachusetts | 40 | female | 21.66 | Alpha(B.1.1.7)/Delta(B.1.617.2) |
| SRR14394850 | 2021/04/03 | Ohio | 33 | male | 15.47 | Alpha(B.1.1.7)/Eta(B.1.525) |
| SRR14812093 | 2021/05/17 | Connecticut | 18 | female | 23.75 | Alpha(B.1.1.7)/Gamma(P.1)/Iota(B.1.526) |
| SRR14452198 | 2021/04/13 | Illinois | 26 | female | 16.62 | Alpha(B.1.1.7)/Gamma(P.1.1) |
| SRR14811846 | 2021/05/15 | Ohio | 66 | female | 20.74 | Alpha(B.1.1.7)/Iota(B.1.526) |
| SRR14811859 | 2021/05/15 | Pennsylvania | 68 | female | 22.58 | Alpha(B.1.1.7)/Iota(B.1.526) |
| SRR14812084 | 2021/05/17 | Maine | 29 | male | 18.25 | Alpha(B.1.1.7)/Iota(B.1.526) |
| SRR14812107 | 2021/05/17 | Massachusetts | 40 | male | 17.20 | Alpha(B.1.1.7)/Iota(B.1.526) |
| SRR14812112 | 2021/05/17 | Maine | 9 | male | 20.10 | Alpha(B.1.1.7)/Iota(B.1.526) |
| SRR14812158 | 2021/05/16 | Maine | 37 | male | 17.43 | Alpha(B.1.1.7)/Iota(B.1.526) |
| SRR14812179 | 2021/05/16 | Maine | 43 | male | 21.08 | Alpha(B.1.1.7)/Iota(B.1.526) |
| SRR14451894 | 2021/04/14 | Florida | 34 | male | 15.03 | Alpha(B.1.1.7)/Iota(B.1.526) |
| SRR14452322 | 2021/04/13 | California | 21 | female | 19.48 | Alpha(B.1.1.7)/Iota(B.1.526) |
| SRR14462551 | 2021/02/23 | Florida | 34 | male | 22.26 | Alpha(B.1.1.7)/Iota(B.1.526) |
| SRR14397499 | 2021/03/22 | Illinois | 24 | male | 22.01 | B.1.1.519/B.1.427 |
| SRR14450785 | 2021/03/04 | Michigan | 68 | male | 22.72 | B.1.214.3/B.1.429 |
| SRR14403671 | 2021/02/11 | Massachusetts | 22 | female | 20.55 | B.1.517/B.1.2 |
| SRR14451452 | 2021/03/05 | Massachusetts | 62 | female | 21.67 | B.1.568/Iota(B.1.526) |
| SRR14450921 | 2021/02/27 | Pennsylvania | 63 | male | 24.73 | B.1.575.1/B.1.429 |
| SRR15493952 | 2021/07/25 | Florida | 62 | male | 24.59 | B.1.621/Delta(B.1.617.2) |
| SRR14453225 | 2021/04/10 | Texas | 46 | female | 16.33 | B.1.628/Gamma(P.1) |
| SRR15432479 | 2021/07/20 | Florida | 71 | male | 20.08 | Delta(AY.2)/Delta(AY.1) |
| SRR15433088 | 2021/07/20 | Illinois | 11 | female | 24.75 | Delta(AY.2)/Delta(AY.3) |
| SRR15432502 | 2021/07/26 | Virginia | 19 | male |  | Delta(AY.2)/Delta(AY.3) |
| SRR15432222 | 2021/07/26 | Wisconsin | 27 | female | 16.64 | Delta(AY.2)/Delta(AY.3) |
| SRR15433019 | 2021/07/26 | Michigan | 11 | male | 11.87 | Delta(AY.2)/Delta(AY.3) |
| SRR15433677 | 2021/07/17 | New Jersey | 50 | male | 22.23 | Delta(AY.2)/Delta(AY.3) |
| SRR15432225 | 2021/07/26 | Wisconsin | 29 | female | 19.11 | Delta(AY.2)/Delta(AY.3) |
| SRR15433533 | 2021/07/25 | Illinois | 15 | female |  | Delta(AY.2)/Delta(AY.3) |
| SRR15432758 | 2021/07/24 | Florida | 22 | male | 14.62 | Delta(AY.2)/Delta(AY.3) |
| SRR15432981 | 2021/07/20 | Minnesota | 29 | female | 23.15 | Delta(AY.2)/Delta(AY.3) |
| SRR15383109 | 2021/07/18 | South Carolina | 14 | female |  | Delta(AY.2)/Delta(AY.3) |
| SRR15383173 | 2021/07/18 | New York | 29 | male | 22.69 | Delta(AY.2)/Delta(AY.3) |
| SRR15399879 | 2021/07/18 | Florida | 21 | male | 18.31 | Delta(AY.2)/Delta(AY.3) |
| SRR15433108 | 2021/07/19 | Ohio | 44 | female | 16.52 | Delta(AY.3)/B.1.619 |
| SRR14391188 | 2021/04/11 | Pennsylvania | 34 | female | 19.45 | Eta(B.1.525)/Iota(B.1.526) |
| SRR14398873 | 2021/03/16 | Ohio | 24 | female | 18.86 | Gamma(P.1.1)/B.1.2 |
| SRR14392570 | 2021/04/05 | Pennsylvania | 56 | male | 24.18 | Gamma(P.1.1)/B.1.427 |
| SRR14452997 | 2021/04/12 | Massachusetts | 25 | male | 24.87 | Gamma(P.1.2)/Iota(B.1.526) |
| SRR14452465 | 2021/04/10 | Arizona | 60 | female | 23.00 | Iota(B.1.526)/B.1.1.519 |

Since 53 co-infected samples were obtained, we made effort to answer the question that whether the co-infected SARS-CoV-2 lineages have lineage tendentiousness, by assigning each pair of co-infected lineages as a connection to build up a comprehensive network (Fig. 5A). In the co-infected network, the Alpha-Iota, Alpha-Gamma (P.1/P.1.1/P.1.2), Alpha-Delta (B.1.617.2) and Delta (AY.2)-Delta (AY. 3) lineage pairs possess the dominant co-infection events, which is consistent with the dominant co-circulation of these lineages in the USA. It is noteworthy that the number and ratio of co-infection events observed in all collected samples seems to increase with time (Fig. 5B) except during Jun. 2021 when the detected SARS-CoV-2 infections in the USA was at a low level (Fig. S3). The co-infection events mainly occurred during Mar. 2021, Apr. 2021, and Jul. 2021, when multiple lineages were co-epidemic in the USA (Fig. 5C). Remarkably, different from the co-infection events during Jan. 2021 to Jun. 2021, the co-infection events in Jul. 2021 were occurred between two sub-lineages in the Delta lineage, which implied the necessity to keep close eyes on the co-circulation of sub-lineages in the future.

**Fig. 5.**
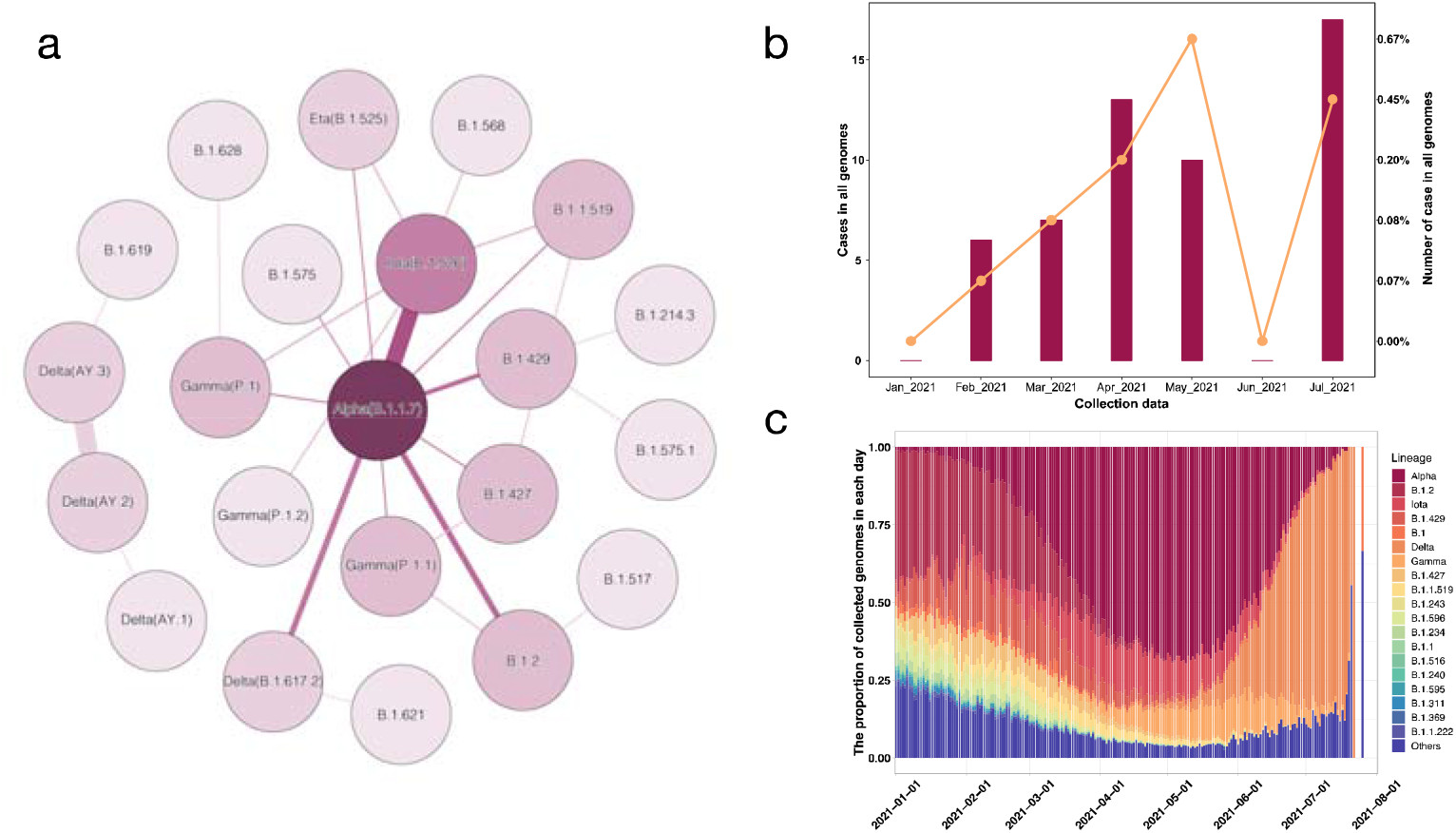
Distribution of co-infection events upon lineage and collection date. **a** Co-infected lineage network for all the identified 53 co-infection events. Every dot represents a lineage, the color depth of each lineage is associated with the occurrence number of this lineage in co-infection events. The thickness of line between dots represents the co-occurrence degree of the linked lineages. The Alpha, Gamma and Delta were the three major lineages that involved in co-infection events. **b** The number and ratio of co-infection events raised with time. **c** The co-circulation pattern of SARS-CoV-2 lineages in the USA. From Feb. 2021 to Apr. 2021, the B.1.2 lineage was outcompeted by Alpha (B.1.1.7), then from Apr. 2021 to Jun. 2021, the Alpha and Gamma lineages were two major co-existed lineages. Later, from Jun. 2021 to now, the Delta lineage has outcompeted all other lineages. *Data have been collected until 31^st^ Jul. 2021.

## Discussion

For most of the SARS-CoV-2 positive samples, no matter infected by one lineage or by multiple lineages, the pattern of mutations in sequencing data fits well with their lineage-defined feature variations. Especially, we observed that the sum of the frequencies of lineage-unique variations was just equal to the average frequencies of their shared variations, demonstrating the co-existence of these lineages within the same sample. Moreover, the epidemiological background of the detected co-infected SARS-CoV-2 samples were highly consisted with the identified lineages for their co-circulations around the sampling locations. The consistence between hypothesis (Fig. 1) and observations (Figs. 3, 4 and S1) provides strong evidence for the detected co-infection events. Also, we believe the co-infection events are truly existed instead of induced by contamination for three reasons. Firstly, the observed co-infection events have obvious lineage tendentiousness rather than random distribution (Fig. 5A). Secondly, we found that the co-infection events were increased with time (Fig. 5B), which was contradictory with the expectation that the contamination samples would decrease with the development of sequencing experience in SARS-CoV-2 detection. Thirdly, the co-infection events occurred with a relatively high rate (~0.18%) in all samples, which was not likely the result of occasionally sequencing contamination.

One question is can we inferred the sources of the co-infection event from their genomic characters? When we assigned variations into lineage(s), we found there were always some undetermined variations. Our further analysis suggested that these undetermined variations could possibly be used to trace the origin of co-infection events. For instance, in a representative co-infected sample (SRR14391243) with two lineages (Fig. 3C), three undetermined variations, NSP13_A598V, Spike_S98F and NS8_C61F, have similar frequencies with the feature variations of the identified Alpha lineage (Fig. 3C). Accordingly, of all the global 2,509,615 viral genomes, only 51 viral strains in Alpha lineage were detected to possess the above three variations as well. Regarding the source of the 51 viral strains, 23 samples were isolated in Ohio State (Fig. S4), demonstrating the Alpha lineage in the co-infected sample might come from a region-specific Alpha lineage circulating in Ohio State. Similarly, five undetermined variations in this co-infected sample were detected to have similar frequencies with feature variations of B.1.2 (Fig. 3C). After scrutinizing all the 2,509,615 viral genomes with the five variations as above, another 51 viral strains could be found. Apparently, different from the situation of Alpha lineage, only 2 strains were isolated in Ohio State while most of the strains with these mutations were in Washington (Fig. S5), demonstrating a complex introduction of the co-infected B.1.2 lineage into Ohio State.

The distribution of co-infection events is region-dependent and time-dependent, indicating that the occurrence of co-infection is result from the interaction between at least two co-circulating SARS-CoV-2 lineages at the particular time and the particular location. For instance, we found that the co-infection events have lineage-bias (Fig. 5A) and increase with time in the USA (Fig. 5B). One possible explanation for this phenomenon is the quick switch of the dominant lineages in the USA during the past seven months. To be specific, with the dominant lineages in the USA changed from B.1.2 to Alpha, the major co-infection lineage has been Alpha lineage in the early of 2021 (Fig. 5A). However, during Jul. 2021, the dominant lineage has been changed from Alpha to Delta strain, followed by the co-infection center switching from Alpha lineage to Delta lineage (Fig. 5A).

To ensure the validity and reliability of our detected co-infection events, we have set a series of stringent criteria (Fig. 2 and Methods). Apart from the determined 53 co-infected samples, 16 additional samples are potentially co-infected (Fig. S6) if we release the parameters slightly. Therefore, we suppose that the co-infected rate of SARS-CoV-2 lineages as ~0.18% in population are under-estimated. If we expand this co-infected rate to all the SARS-CoV-2 infections in the USA or even the whole world, the co-infected patients would be a nonnegligible population. Nevertheless, we cannot make strong conclusion for how co-infected SARS-CoV-2 lineages influence the disease symptoms and the response for disease treatment, which require more research in the future.

## Materials and Methods

### Sample collection

All the 29,993 SRA runs in Project PRJNA716985 were collected from NCBI (https://www.ncbi.nlm.nih.gov). These samples were collected in the USA from Jan. 2021 to Jul. 2021 and sequenced with illumina platform. Samples in this project have been kept with complete meta information, including the collection date, isolated region, sex, and age of patients. To guarantee the accuracy of identified intra-host amino acid variations, only paired-end sequenced samples were selected for further analysis. All the selected samples in the study met the above criteria.

### Calling variations

The collected samples were primarily transformed into FASTQ files with sra-tools. Then the paired-end FASTQ files were imported into CLC Genomics Workbench version 12.0.3 (Qiagen A/S, Vedb æk, Denmark) with Trim Reads Module and mapped to SARS-CoV-2 reference genome (MN908947.3) by using Map Reads to Reference Module. Finally, the Basic Variant Detection Module was used to call within-host nucleotide mutations. These mutations were further converted into amino acid variations by using a homemade Python script.

### Identification of lineage-defined feature variations

The lineage-defined feature variations were defined as the shared lineage-specific signature variations of strains belonging to the same lineage. In general, it was set as the nonsynonymous mutations shared by >= 75% viral strains in a specific lineage (https://outbreak.info/situation-reports/methods#characteristic). However, given the rapid divergence of SARS-CoV-2, many sub-lineages were formed and shared the same feature variations at 75% level, which could not distinguish viral strains belonging to similar lineages. Therefore, in this study, we further introduced the mutations shared by >= 10% viruses to distinguish the neighboring lineages with similar feature variations at 75% level. In total, over 2.5 million SARS-CoV-2 consensus genomes were collected from GISAID^26,27^. All variations that caused nonsynonymous mutations were identified for each viral genome. The lineage of each virus was derived with the Pango nomenclature^3^. A homemade Python script was applied to extract the mutations that shared by >= 75% of all the viruses in one lineage as the 75% feature variations (FV-75). Similarly, mutations shared by >= 10% of all the viruses in one lineage were extracted as 10% feature variations (FV-10). To avoid overfitting, the lineage with few viral genomes globally (<0.01% in all 2.5 million SARS-CoV-2 genomes, or < 250 genomes) were discarded.

### Hypergeometric-distribution based method for detecting SARS-CoV-2 lineage

Under the null hypothesis that each variation has the equal probability of being detected, the number of variations associated with a lineage that overlap with the set of variation follows a hypergeometric distribution. Hence, this process can be conducted using Fisher’s exact test, which uses the hypergeometric distribution. Firstly, all the intra-host variations in NGS raw data were extracted by CLC Genomics Workbench, while feature variations of each lineage were extracted from all the available SARS-CoV-2 consensus sequences in GISAID^26,27^. Secondly, the results were analyzed by a homemade Python script (https://github.com/wuaipinglab/SARS-CoV-2_co-infection). All the potential lineages in a sample could be detected and given a p-value for each test. Thirdly, a judgement process was preformed to evaluate whether the assigned variations in a lineage have similar frequencies, and whether the summed mean frequencies of all lineage-defined feature variations is equal to ~100%. Finally, one lineage infection, multi-lineages co-infection, and infection of other situations were outputted as three individual files, respectively.

### Data and code sharing

The CLC workflow, parameter for CLC modules, ID of screened samples, homemade Python script for identifying potential co-infection events were available online (https://github.com/wuaipinglab/SARS-CoV-2_co-infection).

## Acknowledgments

We thank supporting from The CAMS Initiative for Innovative Medicine (CAMS-I2M, 2016-I2M-1-005) and National Key Plan for Scientific Research and Development of China (2016YFD0500301), China postdoctoral science foundation grant (2019M660548, 2020T130007ZX) and Youthful Teacher Project of Peking Union Medical College (3332019114).

## Author Contributions

A. W. and H.-Y. Z. contributed to the design of this study and the manuscript draft. Y.-X. C. and H.-Y. Z. contributed to the design and encode the homemade Python script for identifying potential co-infection cases. L. X. contributed to the data collection and raw data analysis. J.-Y. L. contributed to data visualization and analysis. C.-Y. T., C.-Y. J. and N. H. contributed to data analysis and workflow optimization. R. Y. contributed to materials preparing. Y. L. provided valuable suggestions on the structure of the article.

## Reference

1. Zhou F, Yu T, Du R, et al. Clinical course and risk factors for mortality of adult inpatients with COVID-19 in Wuhan, China: a retrospective cohort study. The lancet 2020; 395(10229): 1054–62.

2. Dong E, Du H, Gardner L. An interactive web-based dashboard to track COVID-19 in real time. The Lancet infectious diseases 2020; 20(5): 533–4.

3. Rambaut A, Holmes EC, O’Toole Á, et al. A dynamic nomenclature proposal for SARS-CoV-2 lineages to assist genomic epidemiology. Nature microbiology 2020; 5(11): 1403–7.

4. Davies NG, Abbott S, Barnard RC, et al. Estimated transmissibility and impact of SARS-CoV-2 lineage B. 1.1. 7 in England. Science 2021; 372(6538).

5. Hoffmann M, Arora P, Groß R, et al. SARS-CoV-2 variants B. 1.351 and P. 1 escape from neutralizing antibodies. Cell 2021; 184(9): 2384–93. e12.

6. Planas D, Veyer D, Baidaliuk A, et al. Reduced sensitivity of SARS-CoV-2 variant Delta to antibody neutralization. Nature 2021: 1–7.

7. Liu C, Ginn HM, Dejnirattisai W, et al. Reduced neutralization of SARS-CoV-2 B. 1.617 by vaccine and convalescent serum. Cell 2021.

8. Tillett RL, Sevinsky JR, Hartley PD, et al. Genomic evidence for reinfection with SARS-CoV-2: a case study. The Lancet Infectious Diseases 2021; 21(1): 52–8.

9. Sabino EC, Buss LF, Carvalho MP, et al. Resurgence of COVID-19 in Manaus, Brazil, despite high seroprevalence. The Lancet 2021; 397(10273): 452–5.

10. da Silva Francisco Jr R, Benites LF, Lamarca AP, et al. Pervasive transmission of E484K and emergence of VUI-NP13L with evidence of SARS-CoV-2 co-infection events by two different lineages in Rio Grande do Sul, Brazil. Virus research 2021; 296: 198345.

11. Tonkin-Hill G, Martincorena I, Amato R, et al. Patterns of within-host genetic diversity in SARS-CoV-2. BioRxiv 2020.

12. Lythgoe KA, Hall M, Ferretti L, et al. SARS-CoV-2 within-host diversity and transmission. Science 2021; 372(6539).

13. Hashim HO, Mohammed MK, Mousa MJ, et al. Infection with different strains of SARS-CoV-2 in patients with COVID-19. Archives of Biological Sciences 2020; 72(4): 575–85.

14. Samoilov A, Kaptelova V, Bukharina A, et al. Change of dominant strain during dual SARS-CoV-2 infection. medRxiv 2020.

15. Gottlieb GS, Nickle DC, Jensen MA, et al. Dual HIV-1 infection associated with rapid disease progression. The Lancet 2004; 363(9409): 619–22.

16. Van der Kuyl AC, Cornelissen M. Identifying HIV-1 dual infections. Retrovirology 2007; 4(1): 1–12.

17. Ekouevi DK, Eholie SP. Update on HIV-1 and HIV-2 Dual Infection. In: Hope TJ, Stevenson M, Richman D, eds. Encyclopedia of AIDS. New York, NY: Springer New York; 2021: 1–10.

18. Su S, Wong G, Shi W, et al. Epidemiology, genetic recombination, and pathogenesis of coronaviruses. Trends in microbiology 2016; 24(6): 490–502.

19. Zhu Z, Meng K, Meng G. Genomic recombination events may reveal the evolution of coronavirus and the origin of SARS-CoV-2. Scientific reports 2020; 10(1): 1–10.

20. Neches RY, McGee MD, Kyrpides NC. Recombination should not be an afterthought. Nature Reviews Microbiology 2020; 18(11): 606–.

21. Elie B, Lecorche E, Sofonea MT, et al. Inferring SARS-CoV-2 variant within-host kinetics. medRxiv 2021.

22. Ruan Y, Hou M, Li J, et al. One viral sequence for each host?–The neglected within-host diversity as the main stage of SARS-CoV-2 evolution. bioRxiv 2021.

23. Kemp SA, Collier DA, Datir RP, et al. SARS-CoV-2 evolution during treatment of chronic infection. Nature 2021; 592(7853): 277–82.

24. Kim KW, Deveson IW, Pang CNI, et al. Respiratory viral co-infections among SARS-CoV-2 cases confirmed by virome capture sequencing. Scientific reports 2021; 11(1): 1–9.

25. Bull RA, Adikari TN, Ferguson JM, et al. Analytical validity of nanopore sequencing for rapid SARS-CoV-2 genome analysis. Nature communications 2020; 11(1): 1–8.

26. Elbe S, Buckland□Merrett G. Data, disease and diplomacy: GISAID’s innovative contribution to global health. Global Challenges 2017; 1(1): 33–46.

27. Shu Y, McCauley J. GISAID: Global initiative on sharing all influenza data-from vision to reality. Eurosurveillance 2017; 22(13): 30494.

