## Supplemental Information for "Genomic evidence for divergent co-infections of SARS-CoV-2 lineages"

**This PDF file includes:**

Figs. S2 to S4
Captions for Fig. S1 and S6

**Other Supplementary Materials for this manuscript include the following:**

Figs. S1 and S6

**Fig. S1 The 52 samples that were clearly classified to be co-infected by SARS-CoV-2 strains from two different lineages.**

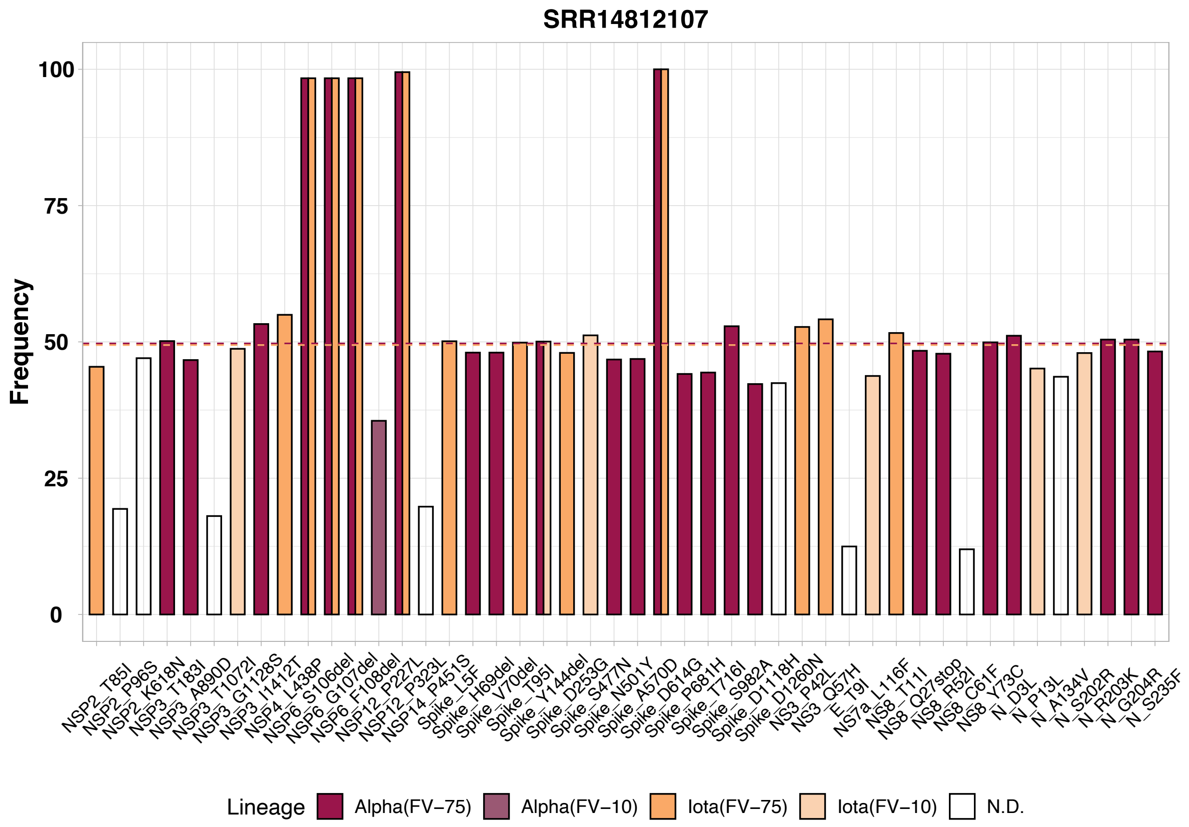


**Fig. S2 The wrongly identified sample.** This sample belongs to two lineage co-infection with all the feature variations of it belong to two identified lineages (Alpha and Iota) as ~50%, respectively.

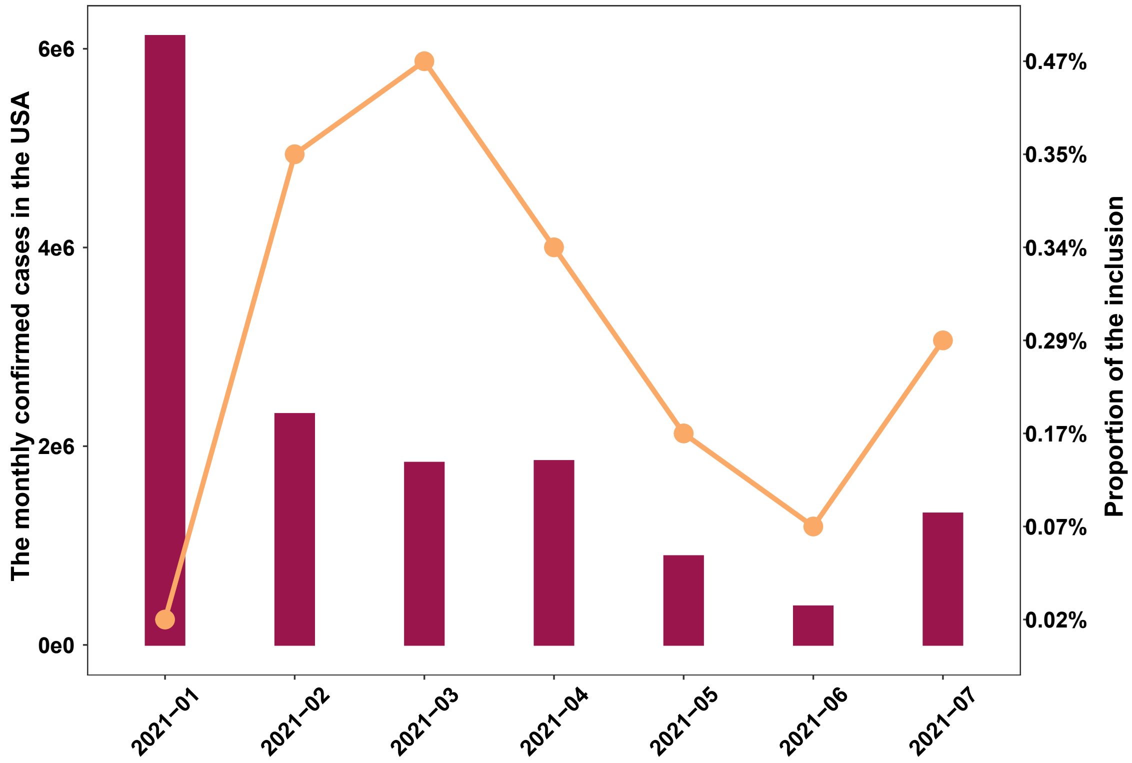


**Fig. S3 The monthly confirmed cases in the USA.** The monthly confirmed cases in the USA was collected from https://github.com/CSSEGISandData/COVID-19/tree/master/csse_19_19_data/csse_covid_19_time_series. Proportion of the inclusion was calculated by dividing the number of monthly confirmed cases by number of the monthly detected samples in this study.

a

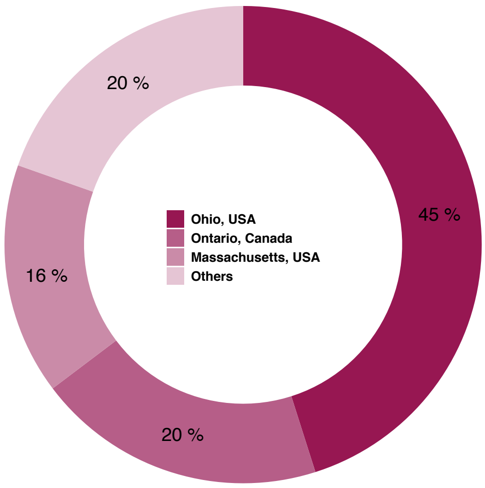


b

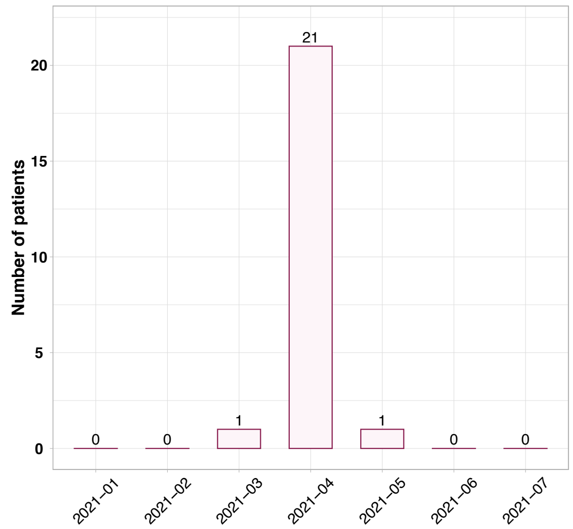


**Fig. S4 The spatio-temporal distribution of SRR14391243 related Alpha lineage.** **a** The spatial distribution of SRR14391243 related Alpha lineage in global. **b** The temporal distribution of SRR14391243 related Alpha lineage.

a

b

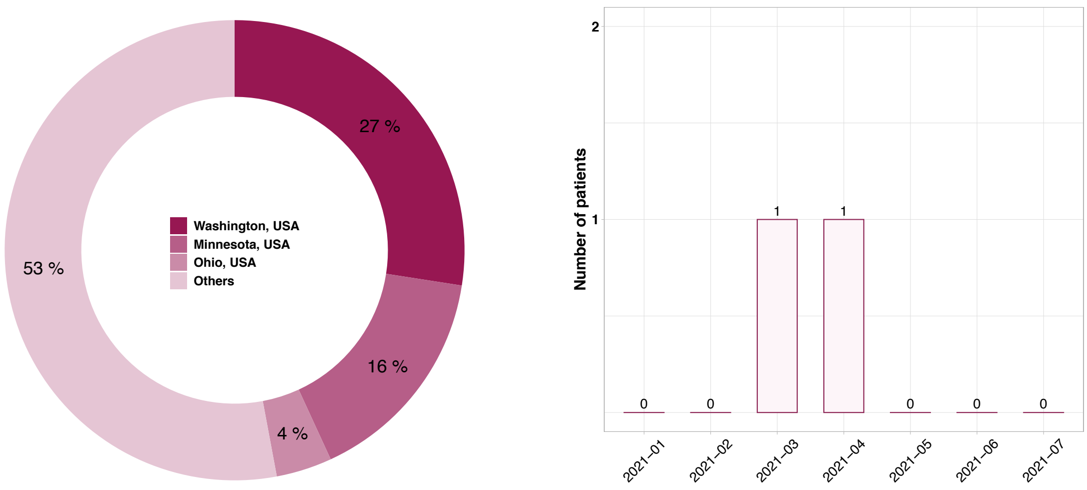


**Fig. S5. The spatio-temporal distribution of SRR14391243 related B.1.2 lineage.** **a** The spatial distribution of SRR14391243 related B.1.2 lineage in global. **b** The temporal distribution of SRR14391243 related B.1.2 lineage.

**Fig. S6. The identified potential co-infection events.**
