## Supplemental Figure 1 for "Genomic evidence for divergent co-infections of SARS-CoV-2 lineages"

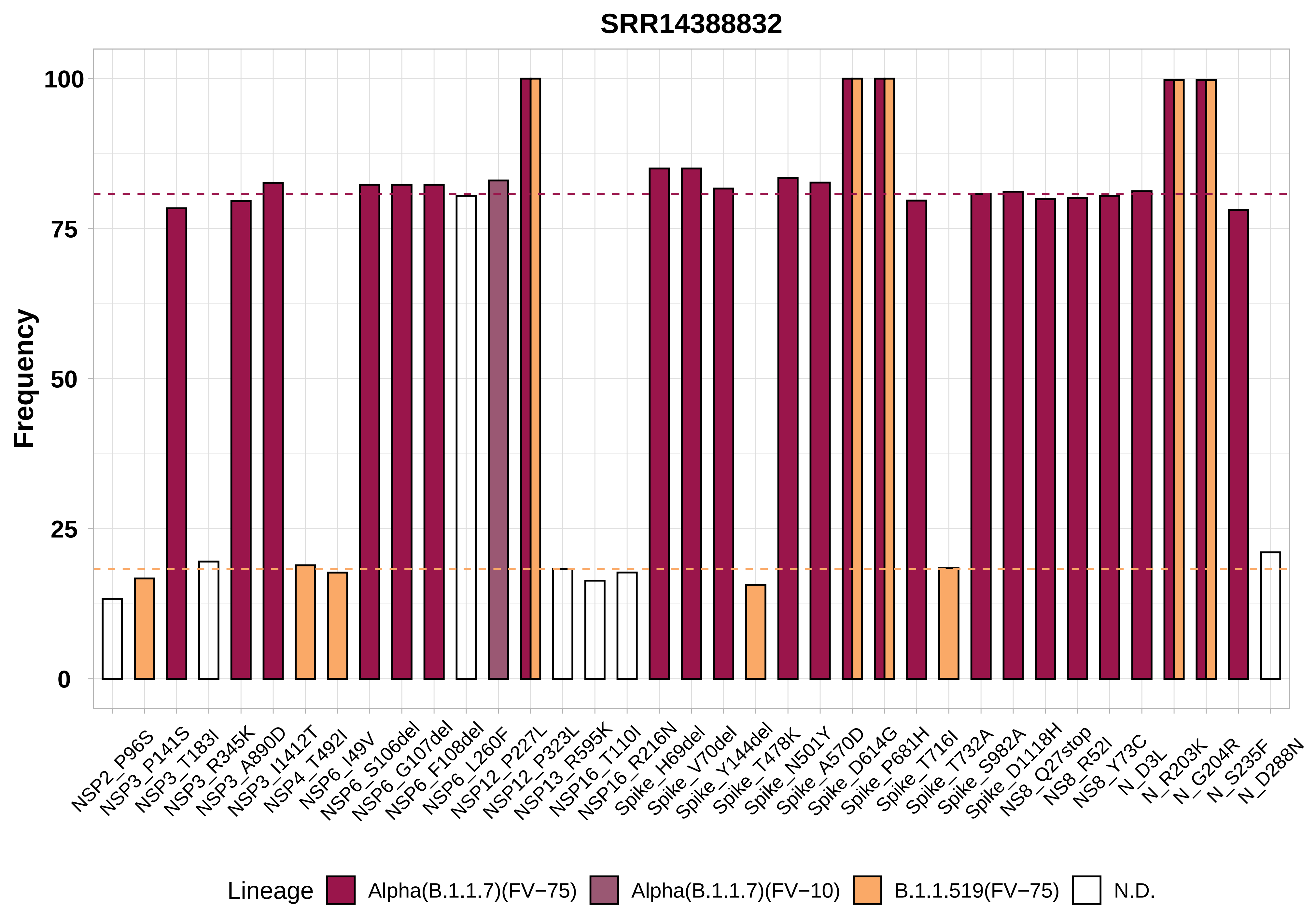
SRR14388832
Frequency
100
75
50
25
0
NSP2_P96S
NSP3_P141S
NSP3_T183I
NSP3_R345K
NSP3_A890D
NSP3_I1412T
NSP4_T492I
NSP6_I49V
NSP6_S106del
NSP6_G107del
NSP6_F108del
NSP6_L260F
NSP12_P227L
NSP12_P323L
NSP13_R595K
NSP16_T110I
NSP16_R216N
Spike_H69del
Spike_V70del
Spike_Y144del
Spike_T478K
Spike_N501Y
Spike_A570D
Spike_D614G
Spike_P681H
Spike_T716I
Spike_T732A
Spike_S982A
Spike_D1118H
NS8_Q27stop
NS8_R52I
NS8_Y73C
N_D3L
N_R203K
N_G204R
N_S235F
N_D288N
Lineage
Alpha(B.1.1.7)(FV−75)
Alpha(B.1.1.7)(FV−10)
B.1.1.519(FV−75)
N.D.

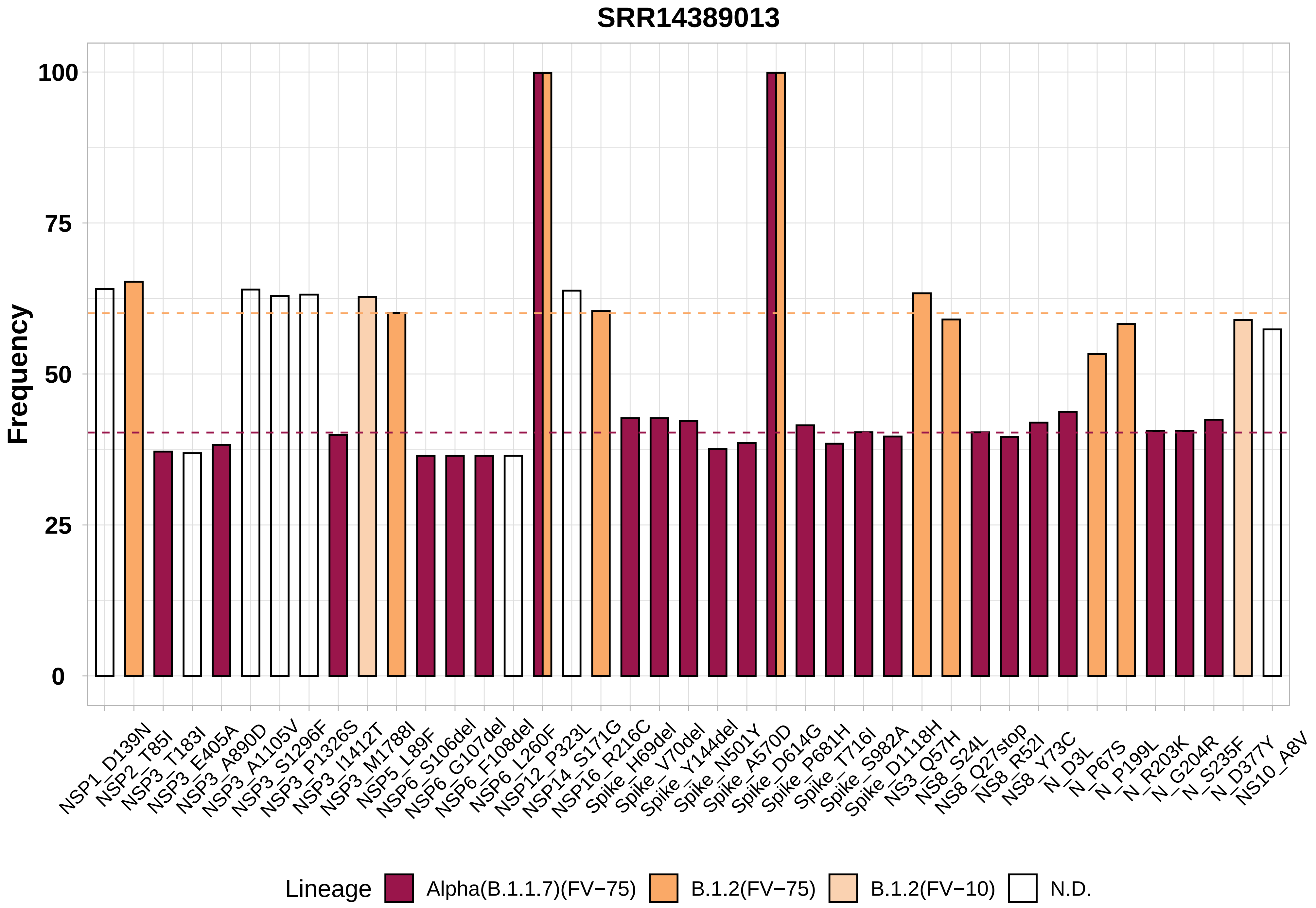

SRR14389013
Frequency
100
75
50
25
0
NSP1_D139N
NSP2_T85I
NSP3_T183I
NSP3_E405A
NSP3_A890D
NSP3_A1105V
NSP3_S1296F
NSP3_P1326S
NSP3_I1412T
NSP3_M1788I
NSP5_L89F
NSP6_S106del
NSP6_G107del
NSP6_F108del
NSP6_L260F
NSP12_P323L
NSP14_S171G
NSP16_R216C
Spike_H69del
Spike_V70del
Spike_Y144del
Spike_N501Y
Spike_A570D
Spike_D614G
Spike_P681H
Spike_T716I
Spike_S982A
Spike_D1118H
NS3_Q57H
NS8_S24L
NS8_Q27stop
NS8_R52I
NS8_Y73C
N_D3L
N_P67S
N_P199L
N_R203K
N_G204R
N_S235F
N_D377Y
NS10_A8V
Lineage
Alpha(B.1.1.7)(FV−75)
B.1.2(FV−75)
B.1.2(FV−10)
N.D.

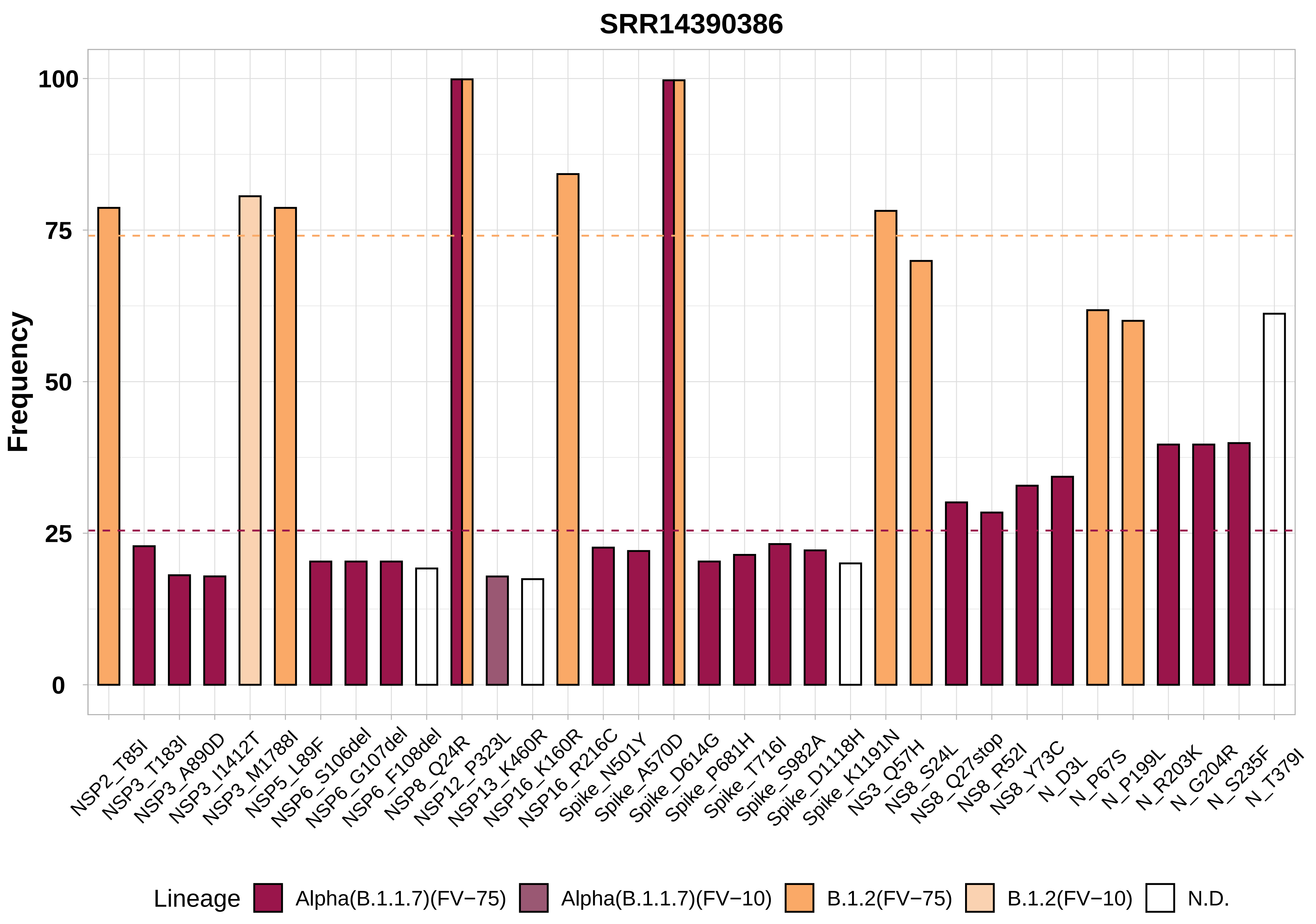
SRR14390386
Frequency
100
75
50
25
0
NSP2_T85I
NSP3_T183I
NSP3_A890D
NSP3_I1412T
NSP3_M1788I
NSP5_L89F
NSP6_S106del
NSP6_G107del
NSP6_F108del
NSP8_Q24R
NSP12_P323L
NSP13_K460R
NSP16_K160R
NSP16_R216C
Spike_N501Y
Spike_A570D
Spike_D614G
Spike_P681H
Spike_T716I
Spike_S982A
Spike_D1118H
Spike_K1191N
NS3_Q57H
NS8_S24L
NS8_Q27stop
NS8_R52I
NS8_Y73C
N_D3L
N_P67S
N_P199L
N_R203K
N_G204R
N_S235F
N_T379I
Lineage
Alpha(B.1.1.7)(FV−75)
Alpha(B.1.1.7)(FV−10)
B.1.2(FV−75)
B.1.2(FV−10)
N.D.

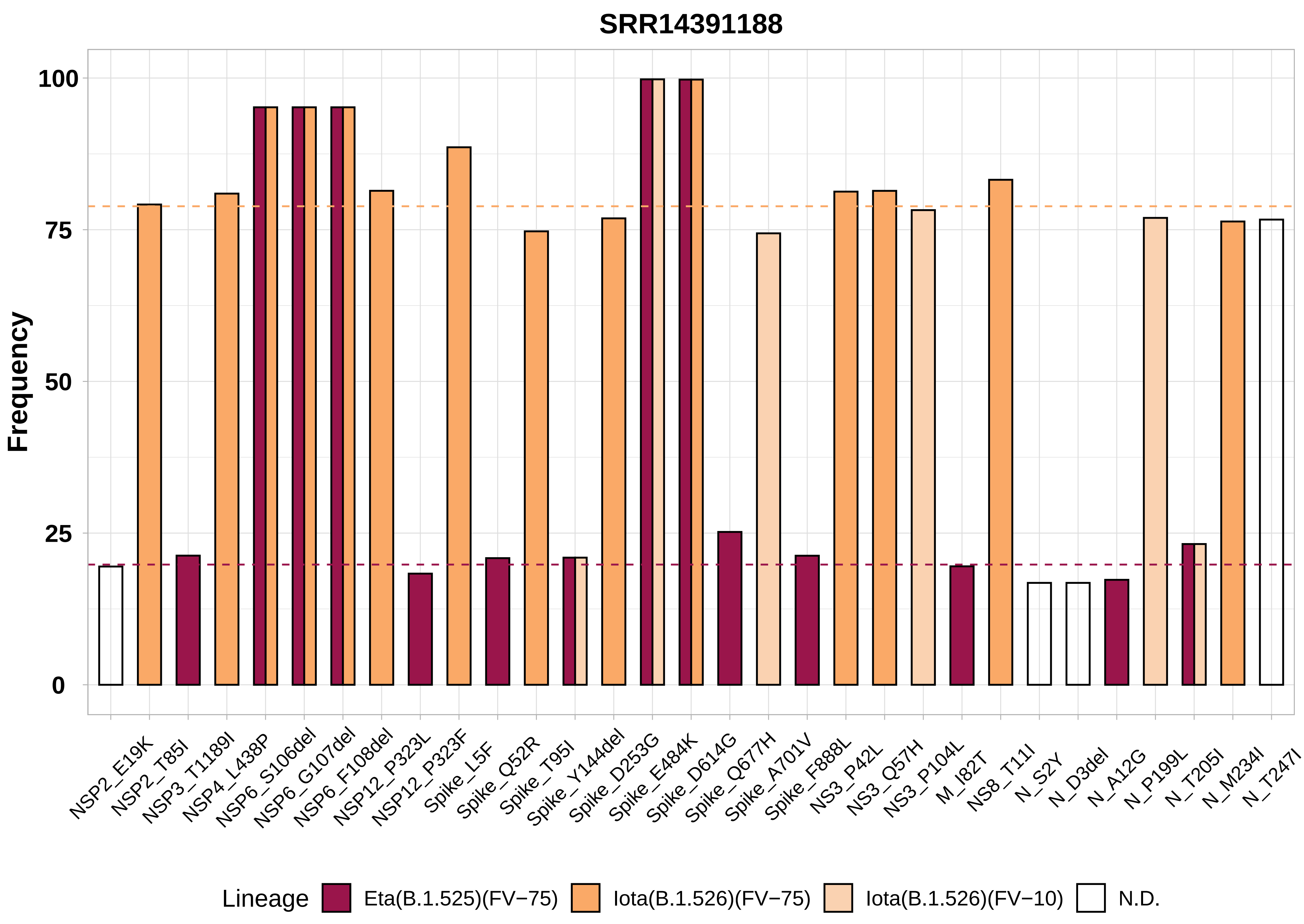
SRR14391188
Frequency
100
75
50
25
0
NSP2_E19K
NSP2_T85I
NSP3_T1189I
NSP4_L438P
NSP6_S106del
NSP6_G107del
NSP6_F108del
NSP12_P323L
NSP12_P323F
Spike_L5F
Spike_Q52R
Spike_T95I
Spike_Y144del
Spike_D253G
Spike_E484K
Spike_D614G
Spike_Q677H
Spike_A701V
Spike_F888L
NS3_P42L
NS3_Q57H
NS3_P104L
M_I82T
NS8_T11I
N_S2Y
N_D3del
N_A12G
N_P199L
N_T205I
N_M234I
N_T247I
Lineage
Eta(B.1.525)(FV−75)
Iota(B.1.526)(FV−75)
Iota(B.1.526)(FV−10)
N.D.

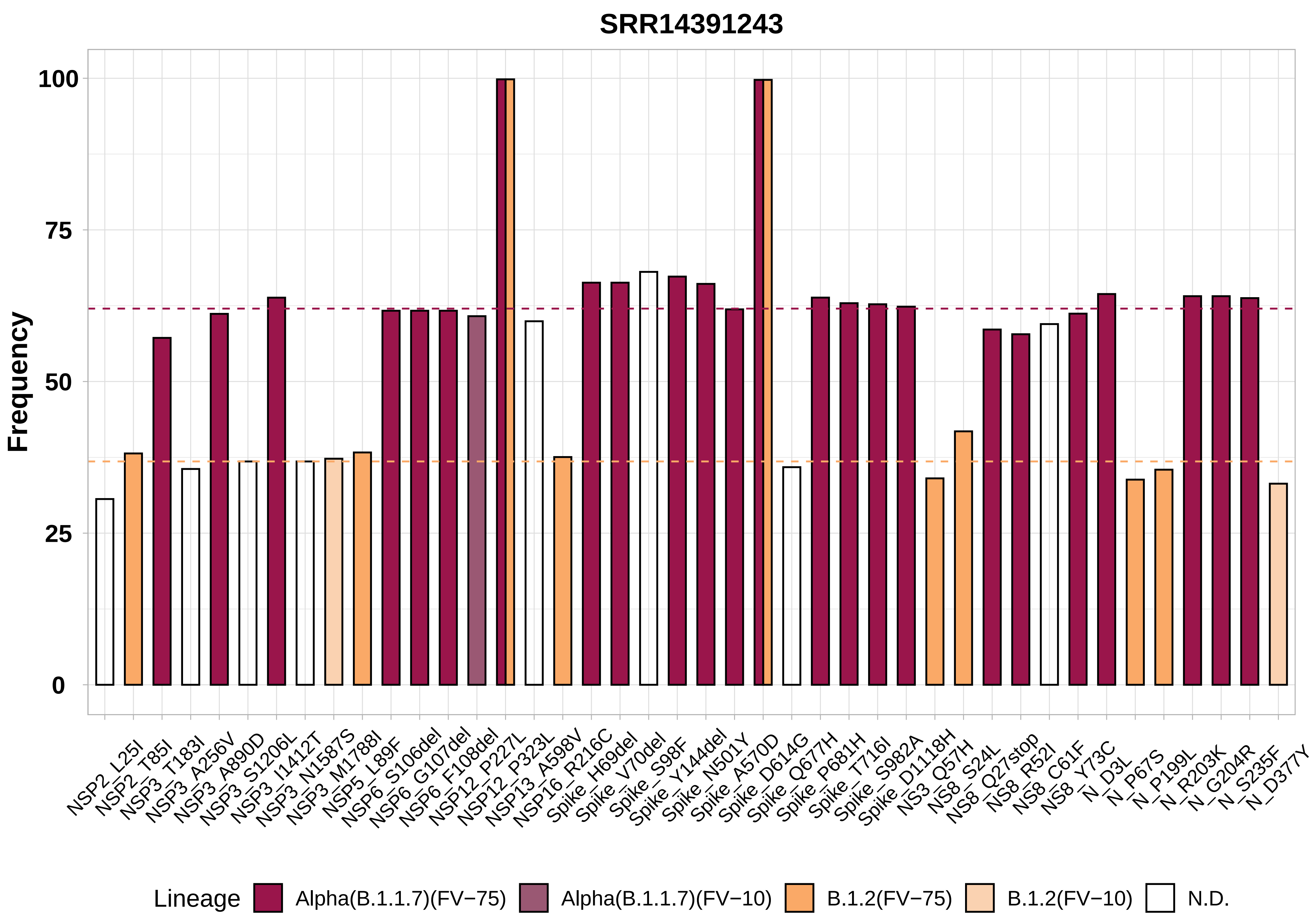
SRR14391243
Frequency
100
75
50
25
0
NSP2_L25I
NSP2_T85I
NSP3_T183I
NSP3_A256V
NSP3_A890D
NSP3_S1206L
NSP3_I1412T
NSP3_N1587S
NSP3_M1788I
NSP5_L89F
NSP6_S106del
NSP6_G107del
NSP6_F108del
NSP12_P227L
NSP12_P323L
NSP13_A598V
NSP16_R216C
Spike_H69del
Spike_V70del
Spike_S98F
Spike_Y144del
Spike_N501Y
Spike_A570D
Spike_D614G
Spike_Q677H
Spike_P681H
Spike_T716I
Spike_S982A
Spike_D1118H
NS3_Q57H
NS8_S24L
NS8_Q27stop
NS8_R52I
NS8_C61F
NS8_Y73C
N_D3L
N_P67S
N_P199L
N_R203K
N_G204R
N_S235F
N_D377Y
Lineage
Alpha(B.1.1.7)(FV−75)
Alpha(B.1.1.7)(FV−10)
B.1.2(FV−75)
B.1.2(FV−10)
N.D.

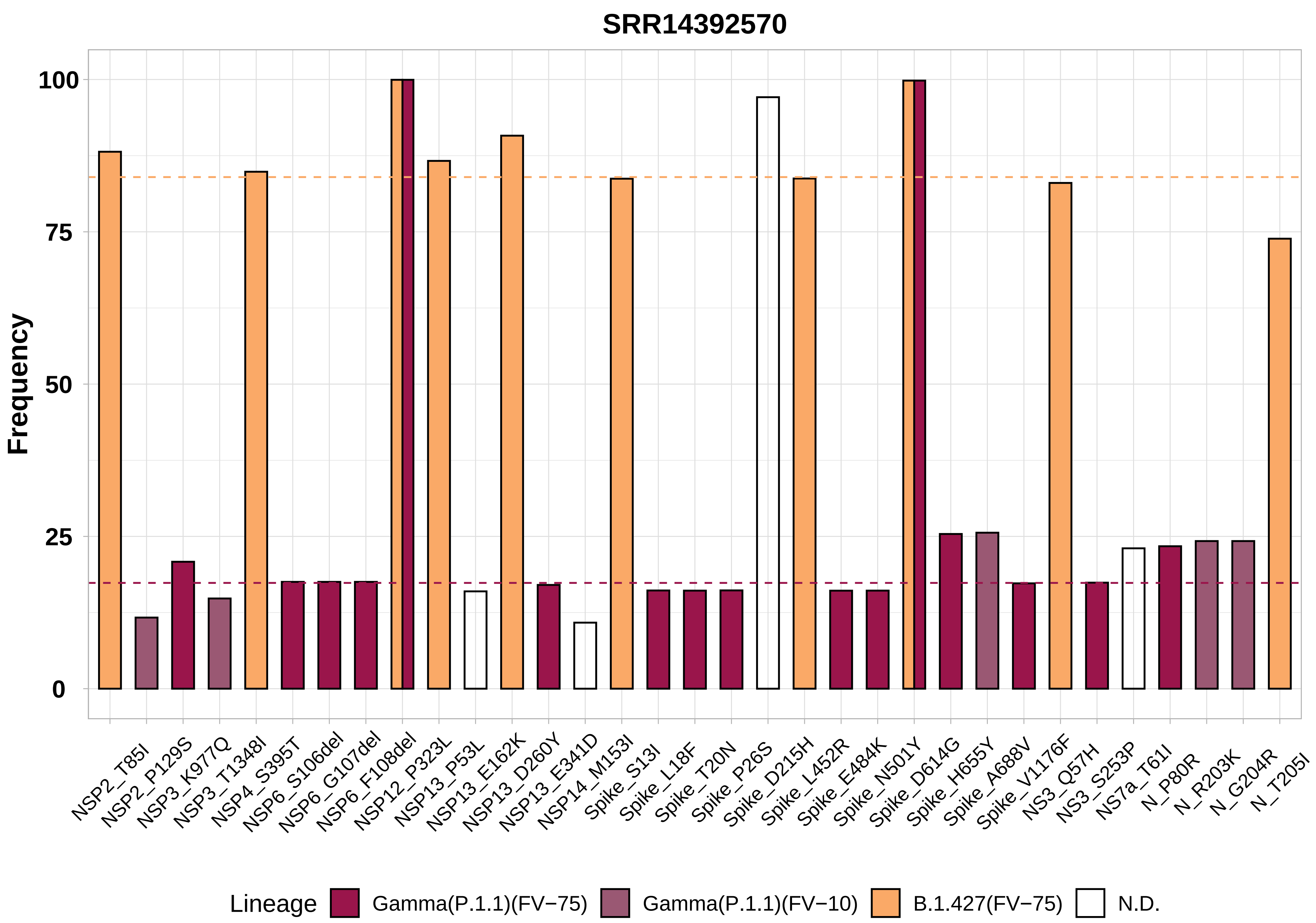

SRR14392570
Frequency
100
75
50
25
0
NSP2_T85I
NSP2_P129S
NSP3_K977Q
NSP3_T1348I
NSP4_S395T
NSP6_S106del
NSP6_G107del
NSP6_F108del
NSP12_P323L
NSP13_P53L
NSP13_E162K
NSP13_D260Y
NSP13_E341D
NSP14_M153I
Spike_S13I
Spike_L18F
Spike_T20N
Spike_P26S
Spike_D215H
Spike_L452R
Spike_E484K
Spike_N501Y
Spike_D614G
Spike_H655Y
Spike_A688V
Spike_V1176F
NS3_Q57H
NS3_S253P
NS7a_T61I
N_P80R
N_R203K
N_G204R
N_T205I
Lineage
Gamma(P.1.1)(FV−75)
Gamma(P.1.1)(FV−10)
B.1.427(FV−75)
N.D.

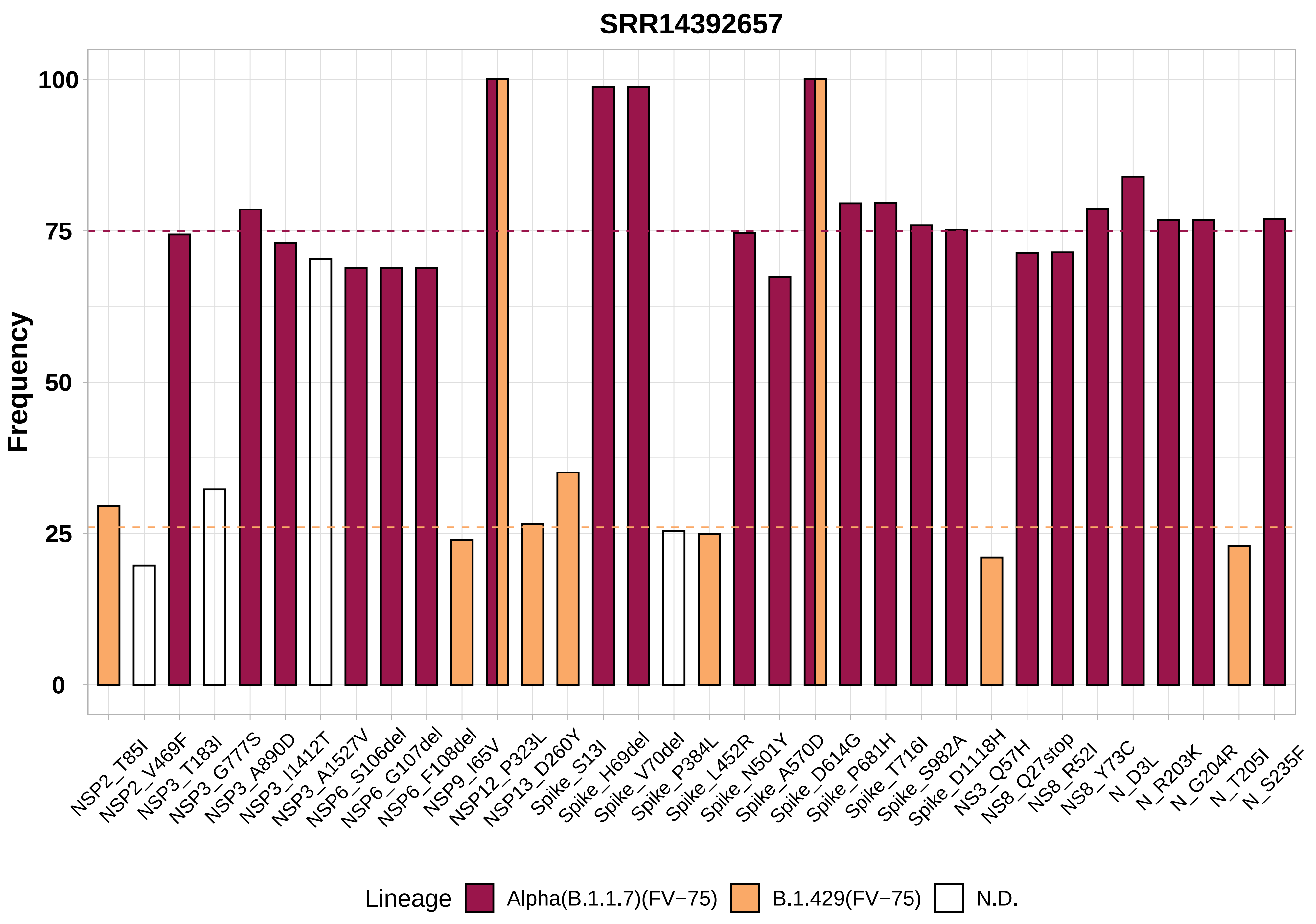

SRR14392657
Frequency
100
75
50
25
0
NSP2_T85I
NSP2_V469F
NSP3_T183I
NSP3_G777S
NSP3_A890D
NSP3_I1412T
NSP3_A1527V
NSP6_S106del
NSP6_G107del
NSP6_F108del
NSP9_I65V
NSP12_P323L
NSP13_D260Y
Spike_S13I
Spike_H69del
Spike_V70del
Spike_P384L
Spike_L452R
Spike_N501Y
Spike_A570D
Spike_D614G
Spike_P681H
Spike_T716I
Spike_S982A
Spike_D1118H
NS3_Q57H
NS8_Q27stop
NS8_R52I
NS8_Y73C
N_D3L
N_R203K
N_G204R
N_T205I
N_S235F
Lineage
Alpha(B.1.1.7)(FV−75)
B.1.429(FV−75)
N.D.

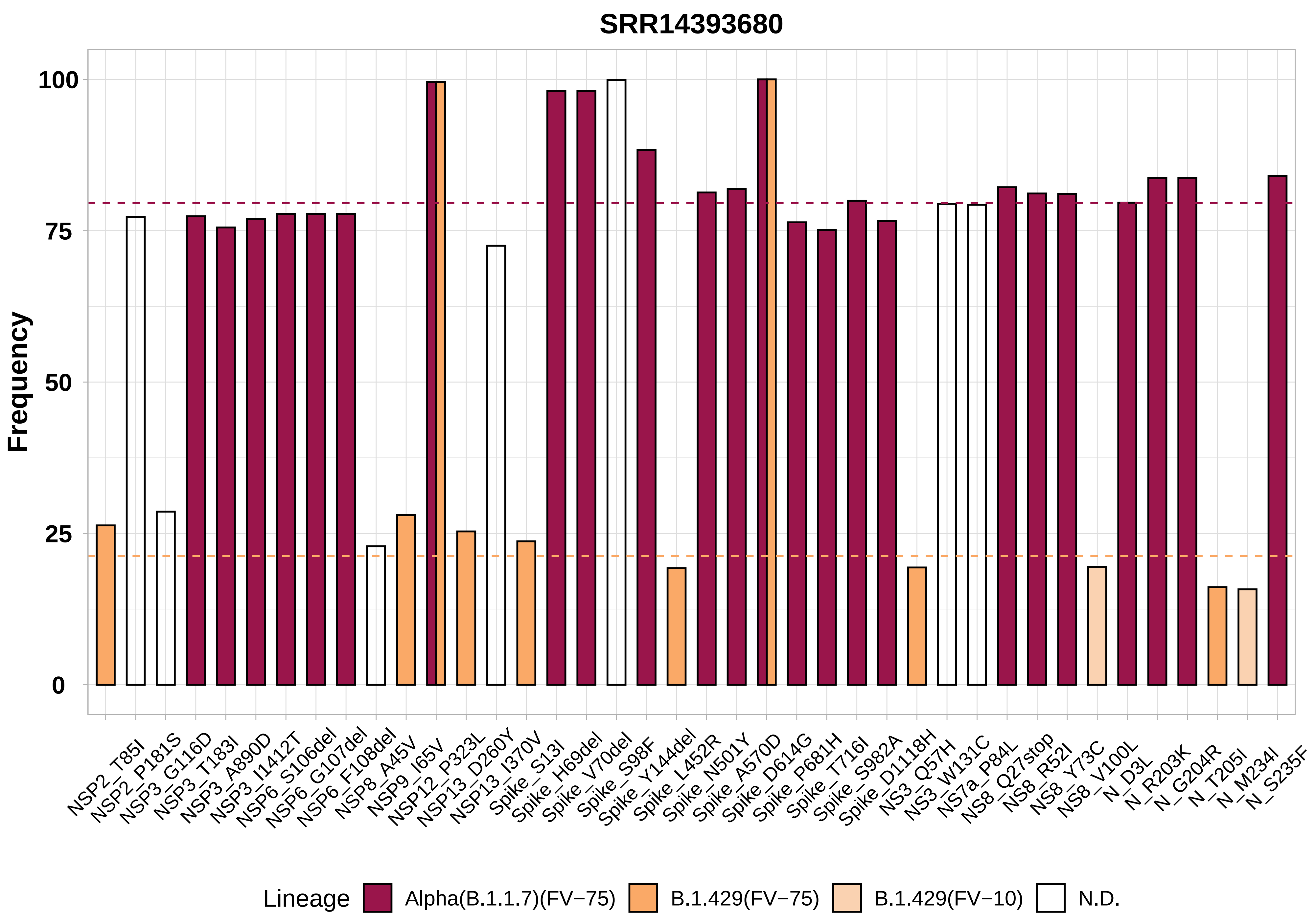

SRR14393680
Frequency
100
75
50
25
0
NSP2_T85I
NSP2_P181S
NSP3_G116D
NSP3_T183I
NSP3_A890D
NSP3_I1412T
NSP6_S106del
NSP6_G107del
NSP6_F108del
NSP8_A45V
NSP9_I65V
NSP12_P323L
NSP13_D260Y
NSP13_I370V
Spike_S13I
Spike_H69del
Spike_V70del
Spike_S98F
Spike_Y144del
Spike_L452R
Spike_N501Y
Spike_A570D
Spike_D614G
Spike_P681H
Spike_T716I
Spike_S982A
Spike_D1118H
NS3_Q57H
NS3_W131C
NS7a_P84L
NS8_Q27stop
NS8_R52I
NS8_Y73C
NS8_V100L
N_D3L
N_R203K
N_G204R
N_T205I
N_M234I
N_S235F
Lineage
Alpha(B.1.1.7)(FV−75)
B.1.429(FV−75)
B.1.429(FV−10)
N.D.

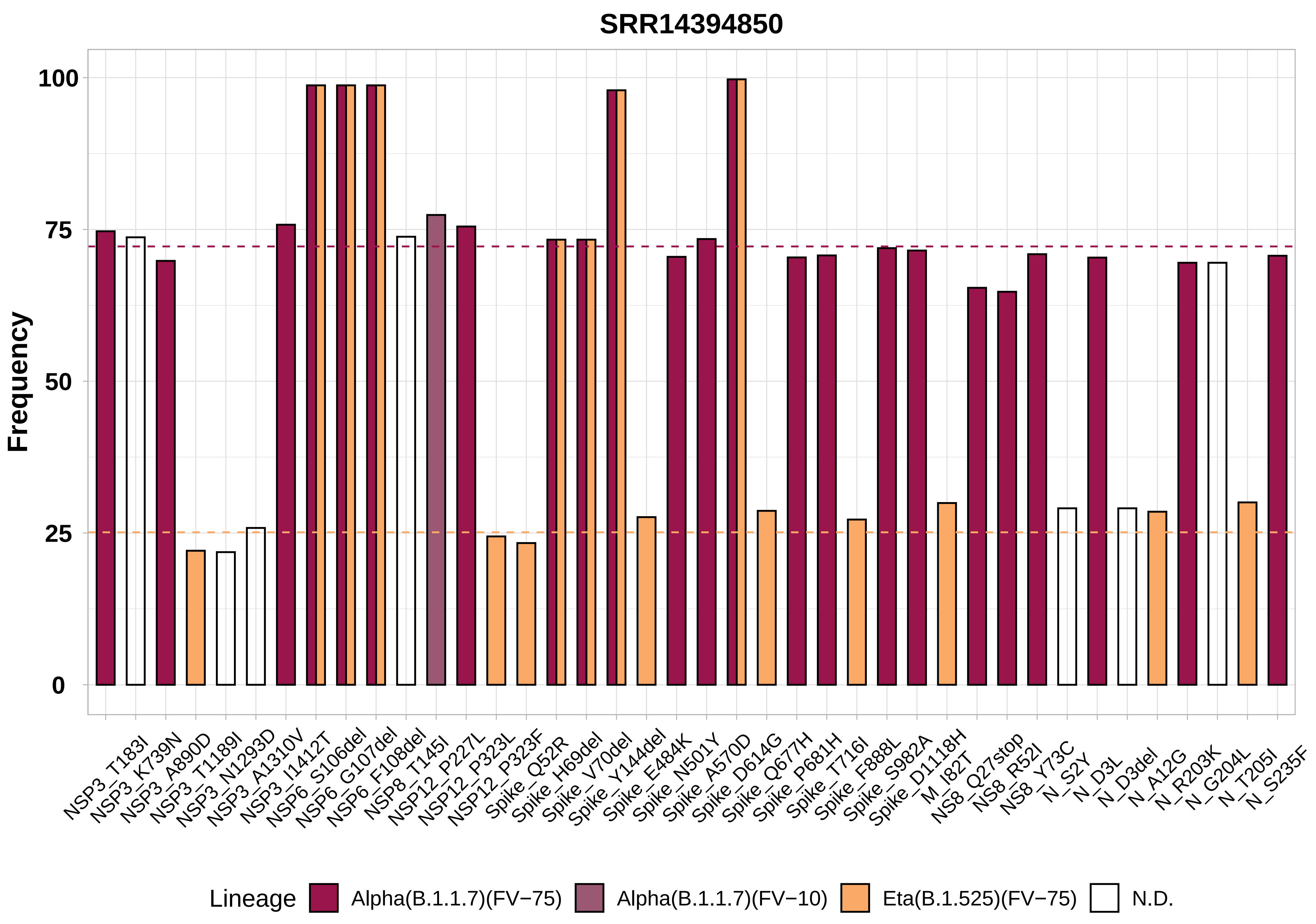

SRR14394850
Frequency
100
75
50
25
0
NSP3_T183I
NSP3_K739N
NSP3_A890D
NSP3_T1189I
NSP3_N1293D
NSP3_A1310V
NSP3_I1412T
NSP6_S106del
NSP6_G107del
NSP6_F108del
NSP8_T145I
NSP12_P227L
NSP12_P323L
NSP12_P323F
Spike_Q52R
Spike_H69del
Spike_V70del
Spike_Y144del
Spike_E484K
Spike_N501Y
Spike_A570D
Spike_D614G
Spike_Q677H
Spike_P681H
Spike_T716I
Spike_F888L
Spike_S982A
Spike_D1118H
M_I82T
NS8_Q27stop
NS8_R52I
NS8_Y73C
N_S2Y
N_D3L
N_D3del
N_A12G
N_R203K
N_G204L
N_T205I
N_S235F
Lineage
Alpha(B.1.1.7)(FV−75)
Alpha(B.1.1.7)(FV−10)
Eta(B.1.525)(FV−75)
N.D.

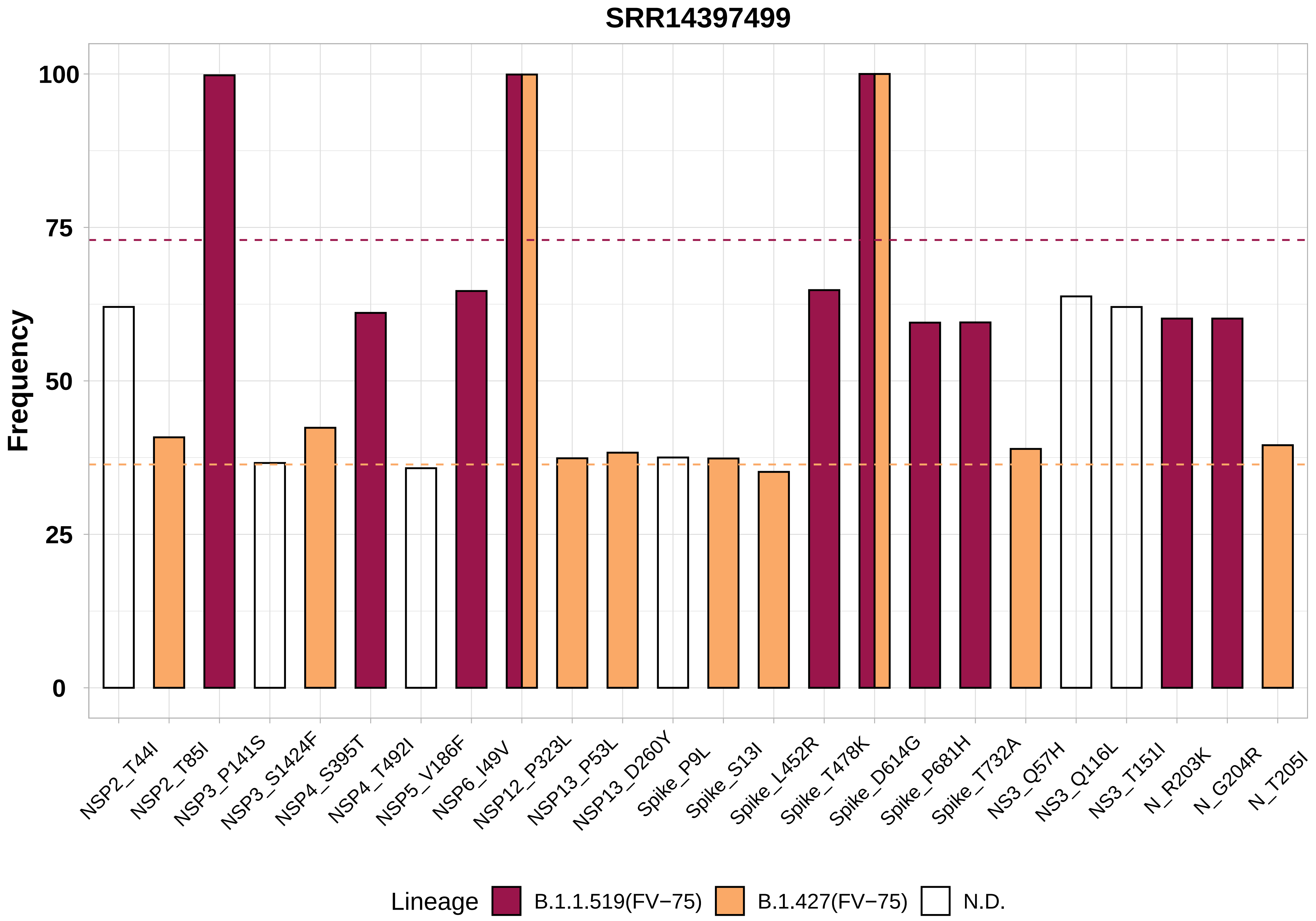

SRR14397499
Frequency
100
75
50
25
0
NSP2_T44I
NSP2_T85I
NSP3_P141S
NSP3_S1424F
NSP4_S395T
NSP4_T492I
NSP5_V186F
NSP6_I49V
NSP12_P323L
NSP13_P53L
NSP13_D260Y
Spike_P9L
Spike_S13I
Spike_L452R
Spike_T478K
Spike_D614G
Spike_P681H
Spike_T732A
NS3_Q57H
NS3_Q116L
NS3_T151I
N_R203K
N_G204R
N_T205I
Lineage
B.1.1.519(FV−75)
B.1.427(FV−75)
N.D.

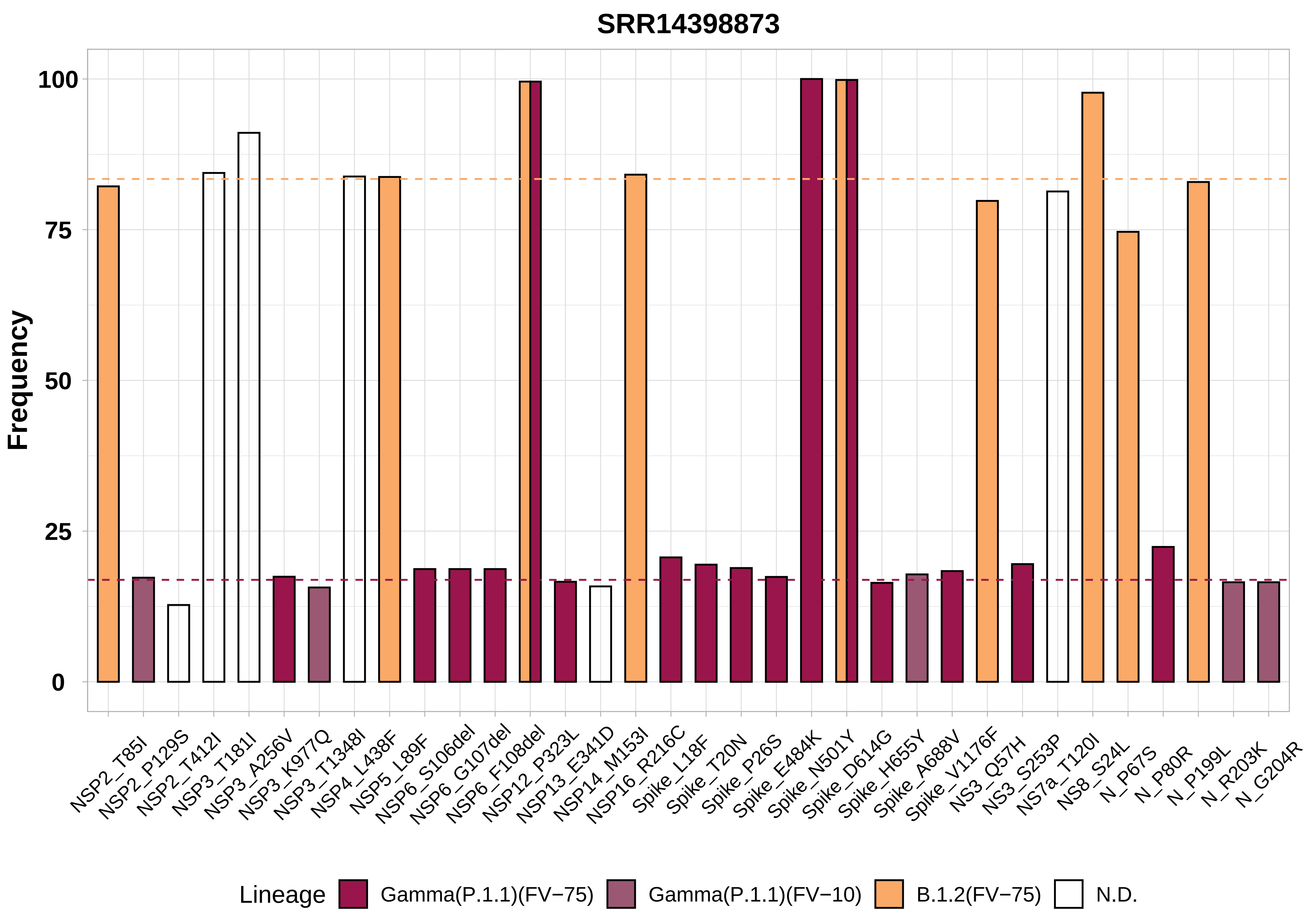

SRR14398873
Frequency
100
75
50
25
0
NSP2_T85I
NSP2_P129S
NSP2_T412I
NSP3_T181I
NSP3_A256V
NSP3_K977Q
NSP3_T1348I
NSP4_L438F
NSP5_L89F
NSP6_S106del
NSP6_G107del
NSP6_F108del
NSP12_P323L
NSP13_E341D
NSP14_M153I
NSP16_R216C
Spike_L18F
Spike_T20N
Spike_P26S
Spike_E484K
Spike_N501Y
Spike_D614G
Spike_H655Y
Spike_A688V
Spike_V1176F
NS3_Q57H
NS3_S253P
NS7a_T120I
NS8_S24L
N_P67S
N_P80R
N_P199L
N_R203K
N_G204R
Lineage
Gamma(P.1.1)(FV−75)
Gamma(P.1.1)(FV−10)
B.1.2(FV−75)
N.D.

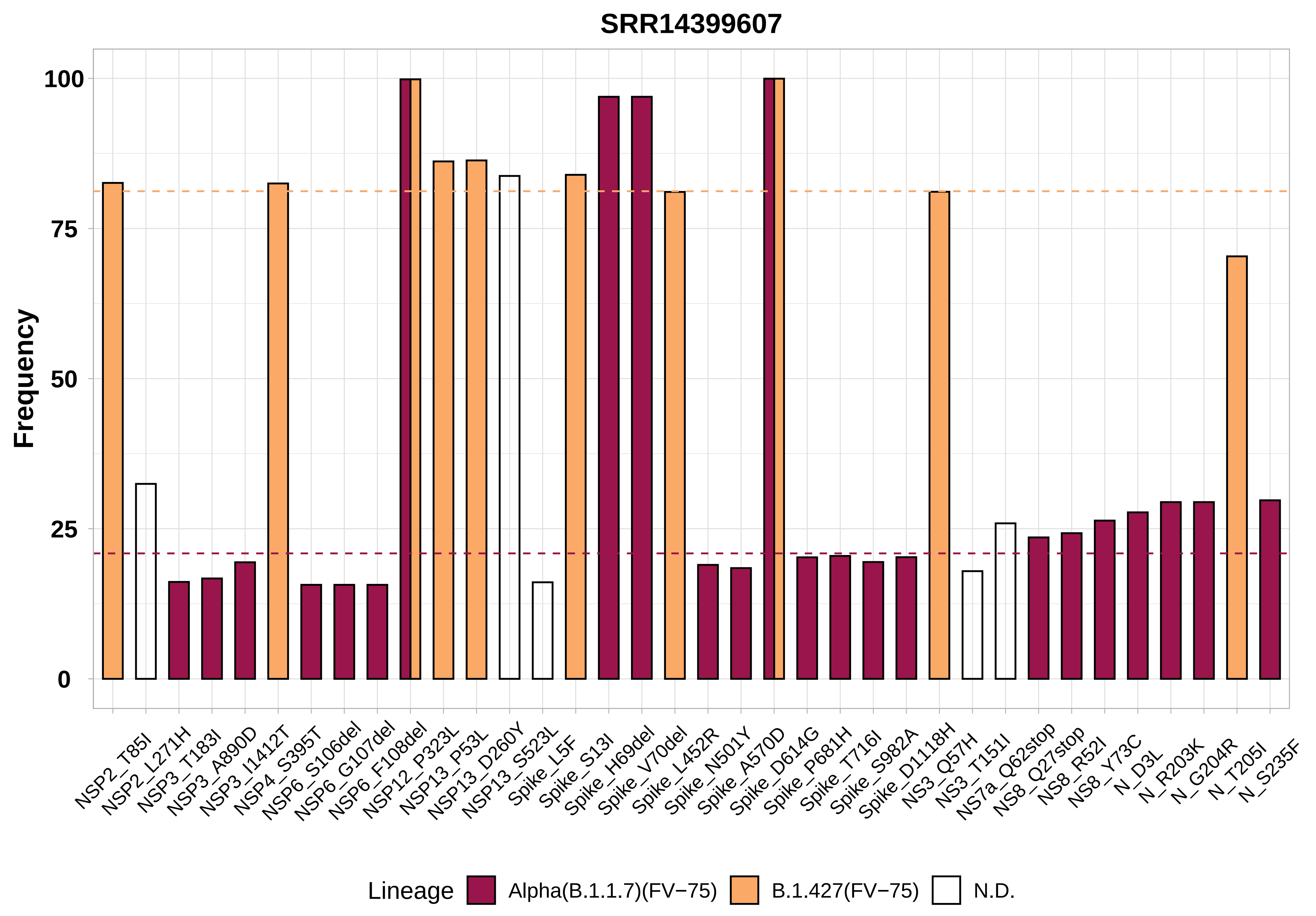

SRR14399607
Frequency
100
75
50
25
0
NSP2_T85I
NSP2_L271H
NSP3_T183I
NSP3_A890D
NSP3_I1412T
NSP4_S395T
NSP6_S106del
NSP6_G107del
NSP6_F108del
NSP12_P323L
NSP13_P53L
NSP13_D260Y
NSP13_S523L
Spike_L5F
Spike_S13I
Spike_H69del
Spike_V70del
Spike_L452R
Spike_N501Y
Spike_A570D
Spike_D614G
Spike_P681H
Spike_T716I
Spike_S982A
Spike_D1118H
NS3_Q57H
NS3_T151I
NS7a_Q62stop
NS8_Q27stop
NS8_R52I
NS8_Y73C
N_D3L
N_R203K
N_G204R
N_T205I
N_S235F
Lineage
Alpha(B.1.1.7)(FV−75)
B.1.427(FV−75)
N.D.

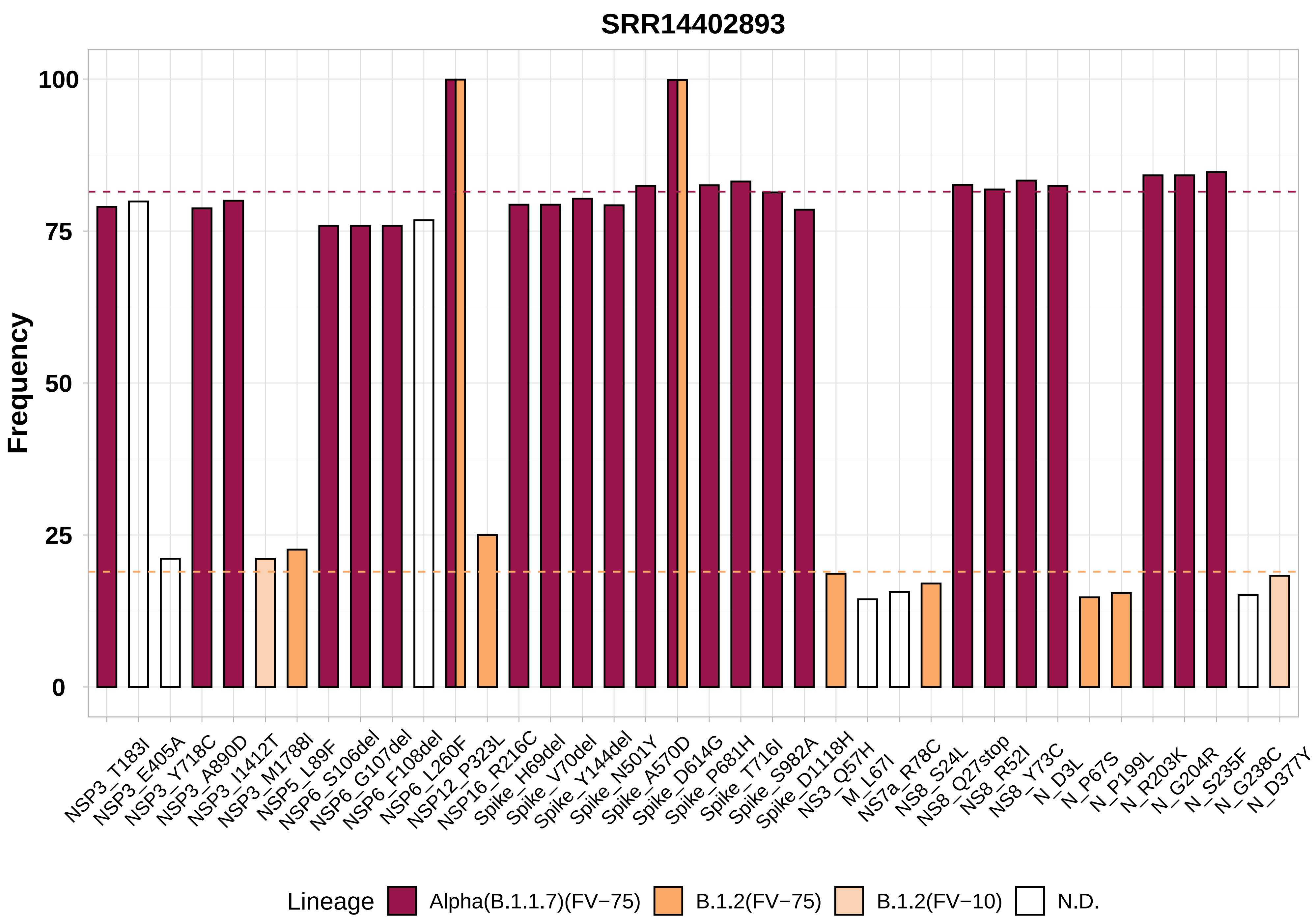
SRR14402893
Frequency
100
75
50
25
0
NSP3_T183I
NSP3_E405A
NSP3_Y718C
NSP3_A890D
NSP3_I1412T
NSP3_M1788I
NSP5_L89F
NSP6_S106del
NSP6_G107del
NSP6_F108del
NSP6_L260F
NSP12_P323L
NSP16_R216C
Spike_H69del
Spike_V70del
Spike_Y144del
Spike_N501Y
Spike_A570D
Spike_D614G
Spike_P681H
Spike_T716I
Spike_S982A
Spike_D1118H
NS3_Q57H
M_L67I
NS7a_R78C
NS8_S24L
NS8_Q27stop
NS8_R52I
NS8_Y73C
N_D3L
N_P67S
N_P199L
N_R203K
N_G204R
N_S235F
N_G238C
N_D377Y
Lineage
Alpha(B.1.1.7)(FV−75)
B.1.2(FV−75)
B.1.2(FV−10)
N.D.

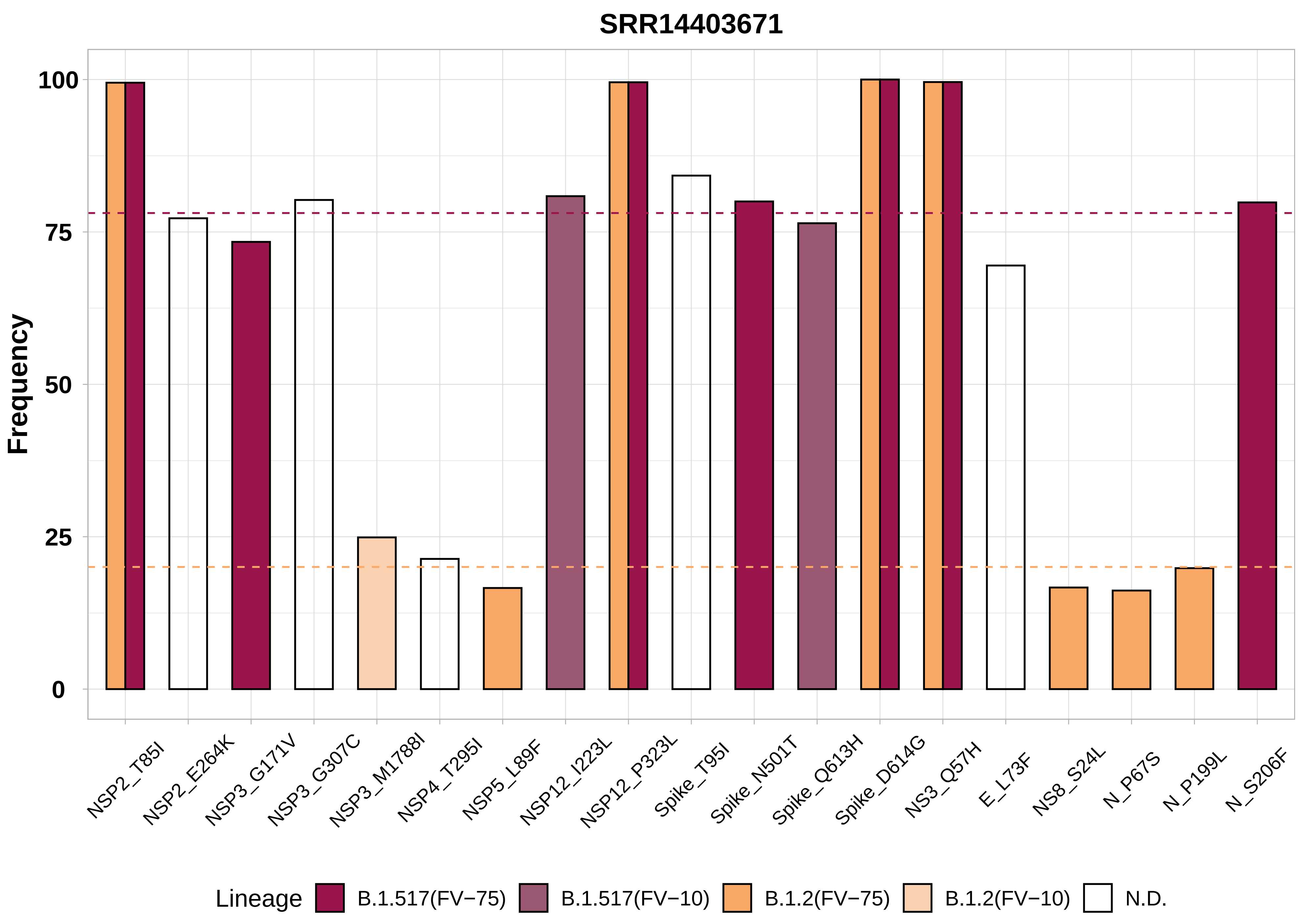
SRR14403671
Frequency
100
75
50
25
0
NSP2_T85I
NSP2_E264K
NSP3_G171V
NSP3_G307C
NSP3_M1788I
NSP4_T295I
NSP5_L89F
NSP12_I223L
NSP12_P323L
Spike_T95I
Spike_N501T
Spike_Q613H
Spike_D614G
NS3_Q57H
E_L73F
NS8_S24L
N_P67S
N_P199L
N_S206F
Lineage
B.1.517(FV−75)
B.1.517(FV−10)
B.1.2(FV−75)
B.1.2(FV−10)
N.D.

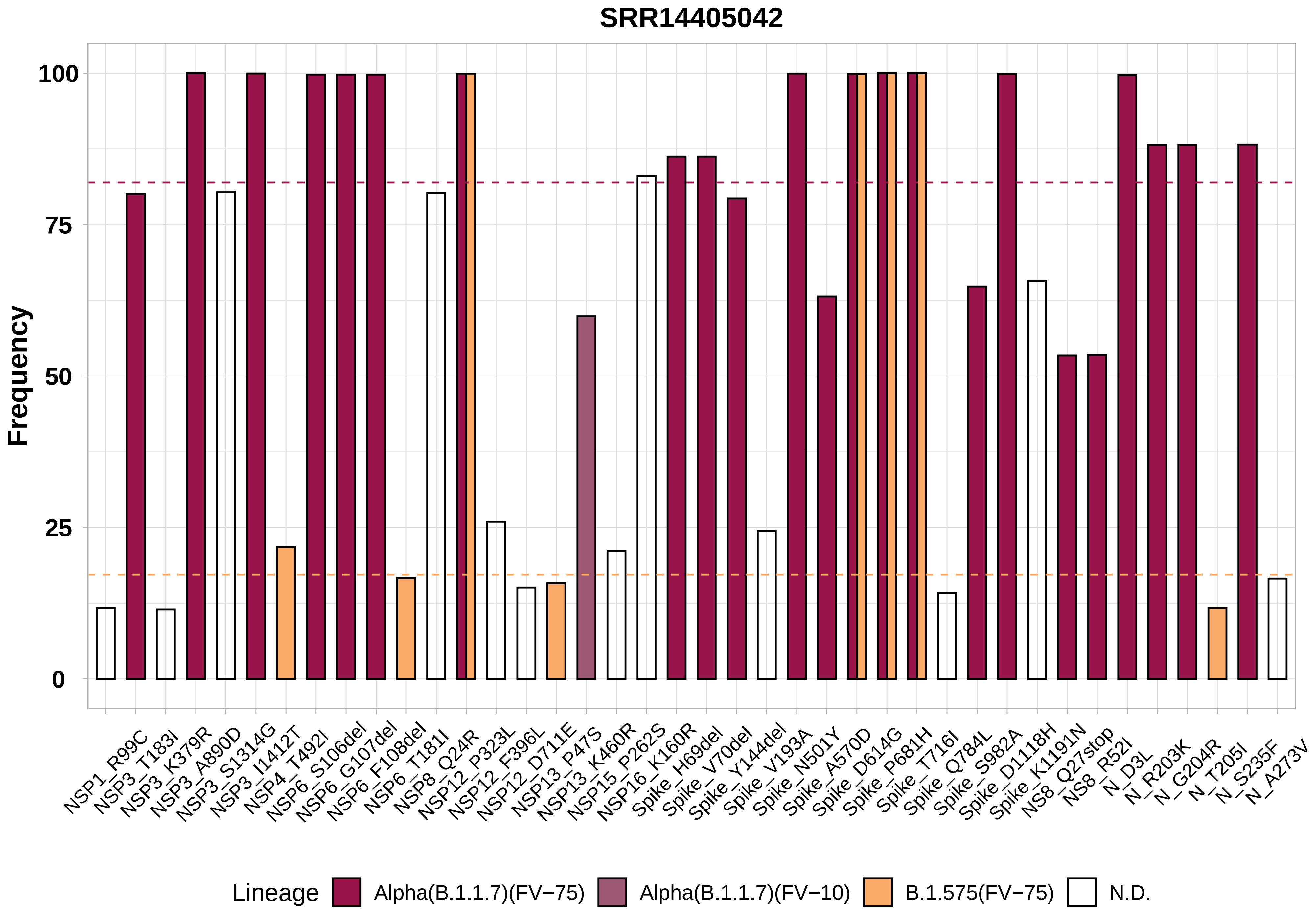

SRR14405042
Frequency
100
75
50
25
0
NSP1_R99C
NSP3_T183I
NSP3_K379R
NSP3_A890D
NSP3_S1314G
NSP3_I1412T
NSP4_T492I
NSP6_S106del
NSP6_G107del
NSP6_F108del
NSP6_T181I
NSP8_Q24R
NSP12_P323L
NSP12_F396L
NSP12_D711E
NSP13_P47S
NSP13_K460R
NSP15_P262S
NSP16_K160R
Spike_H69del
Spike_V70del
Spike_Y144del
Spike_V193A
Spike_N501Y
Spike_A570D
Spike_D614G
Spike_P681H
Spike_T716I
Spike_Q784L
Spike_S982A
Spike_D1118H
Spike_K1191N
NS8_Q27stop
NS8_R52I
N_D3L
N_R203K
N_G204R
N_T205I
N_S235F
N_A273V
Lineage
Alpha(B.1.1.7)(FV−75)
Alpha(B.1.1.7)(FV−10)
B.1.575(FV−75)
N.D.

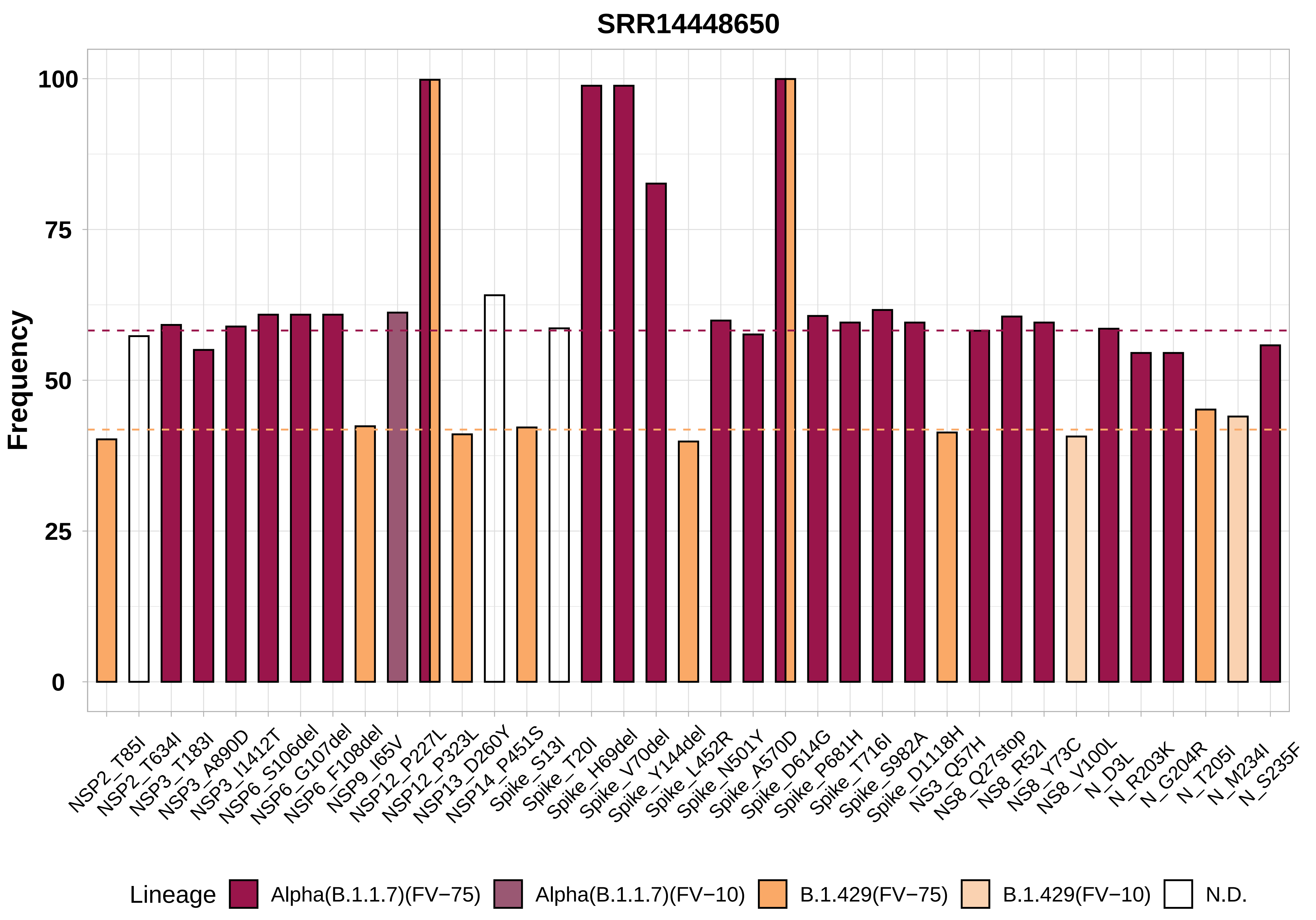

SRR14448650
Frequency
100
75
50
25
0
NSP2_T85I
NSP2_T634I
NSP3_T183I
NSP3_A890D
NSP3_I1412T
NSP6_S106del
NSP6_G107del
NSP6_F108del
NSP9_I65V
NSP12_P227L
NSP12_P323L
NSP13_D260Y
NSP14_P451S
Spike_S13I
Spike_T20I
Spike_H69del
Spike_V70del
Spike_Y144del
Spike_L452R
Spike_N501Y
Spike_A570D
Spike_D614G
Spike_P681H
Spike_T716I
Spike_S982A
Spike_D1118H
NS3_Q57H
NS8_Q27stop
NS8_R52I
NS8_Y73C
NS8_V100L
N_D3L
N_R203K
N_G204R
N_T205I
N_M234I
N_S235F
Lineage
Alpha(B.1.1.7)(FV−75)
Alpha(B.1.1.7)(FV−10)
B.1.429(FV−75)
B.1.429(FV−10)
N.D.

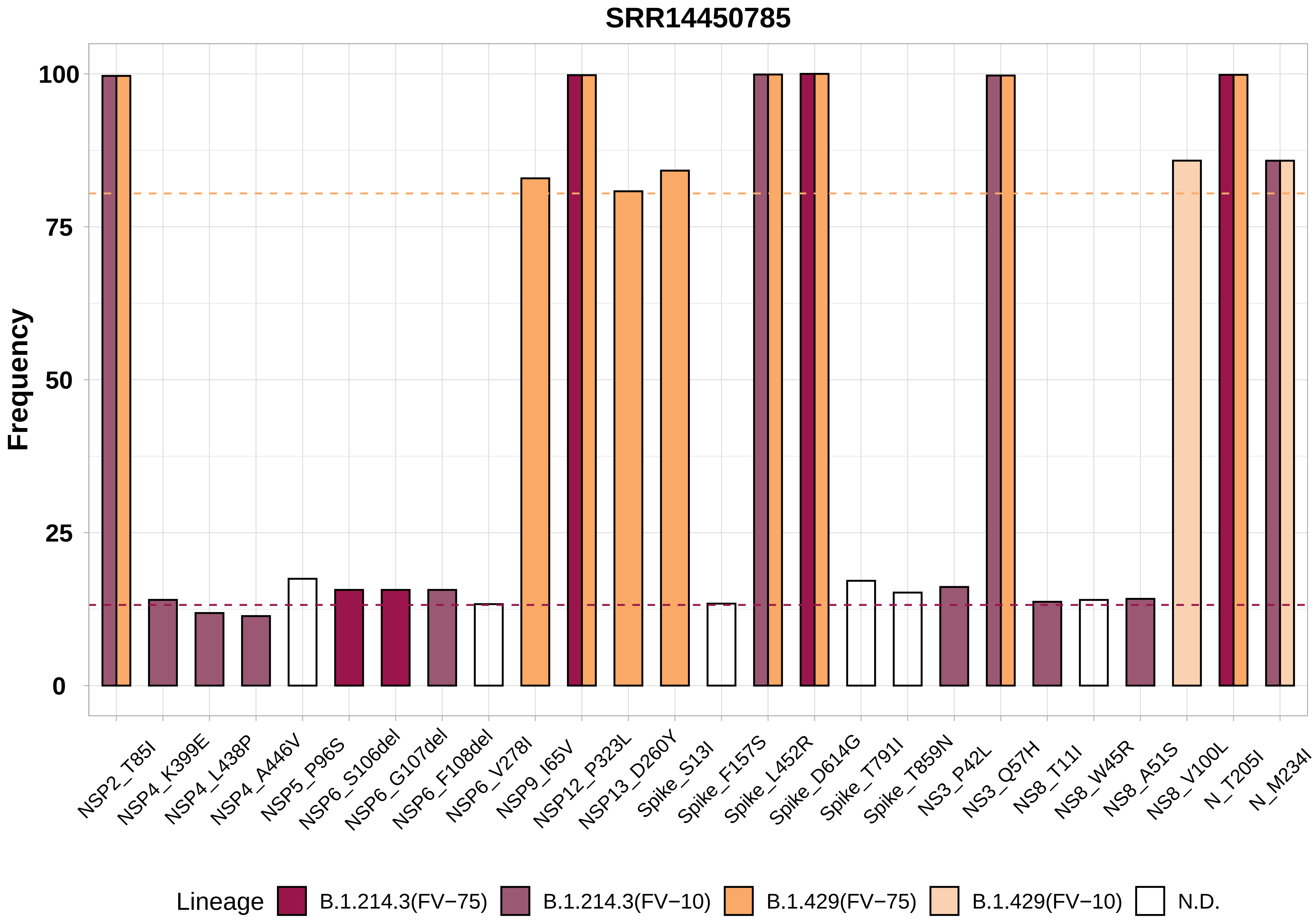

SRR14450785
Frequency
100
75
50
25
0
NSP2_T85I
NSP4_K399E
NSP4_L438P
NSP4_A446V
NSP5_P96S
NSP6_S106del
NSP6_G107del
NSP6_F108del
NSP6_V278I
NSP9_I65V
NSP12_P323L
NSP13_D260Y
Spike_S13I
Spike_F157S
Spike_L452R
Spike_D614G
Spike_T791I
Spike_T859N
NS3_P42L
NS3_Q57H
NS8_T11I
NS8_W45R
NS8_A51S
NS8_V100L
N_T205I
N_M234I
Lineage
B.1.214.3(FV−75)
B.1.214.3(FV−10)
B.1.429(FV−75)
B.1.429(FV−10)
N.D.

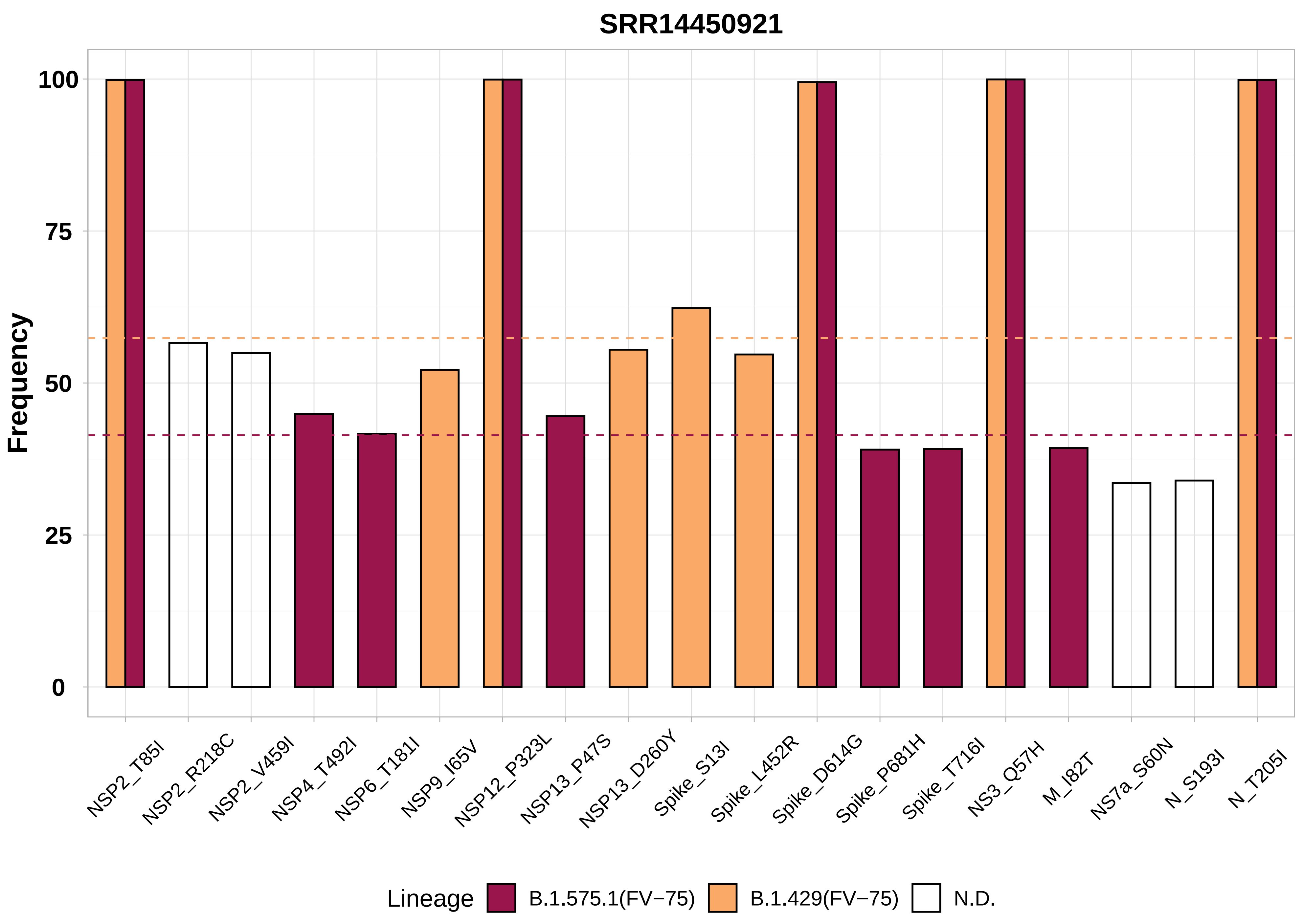

SRR14450921
Frequency
100
75
50
25
0
NSP2_T85I
NSP2_R218C
NSP2_V459I
NSP4_T492I
NSP6_T181I
NSP9_I65V
NSP12_P323L
NSP13_P47S
NSP13_D260Y
Spike_S13I
Spike_L452R
Spike_D614G
Spike_P681H
Spike_T716I
NS3_Q57H
M_I82T
NS7a_S60N
N_S193I
N_T205I
Lineage
B.1.575.1(FV−75)
B.1.429(FV−75)
N.D.

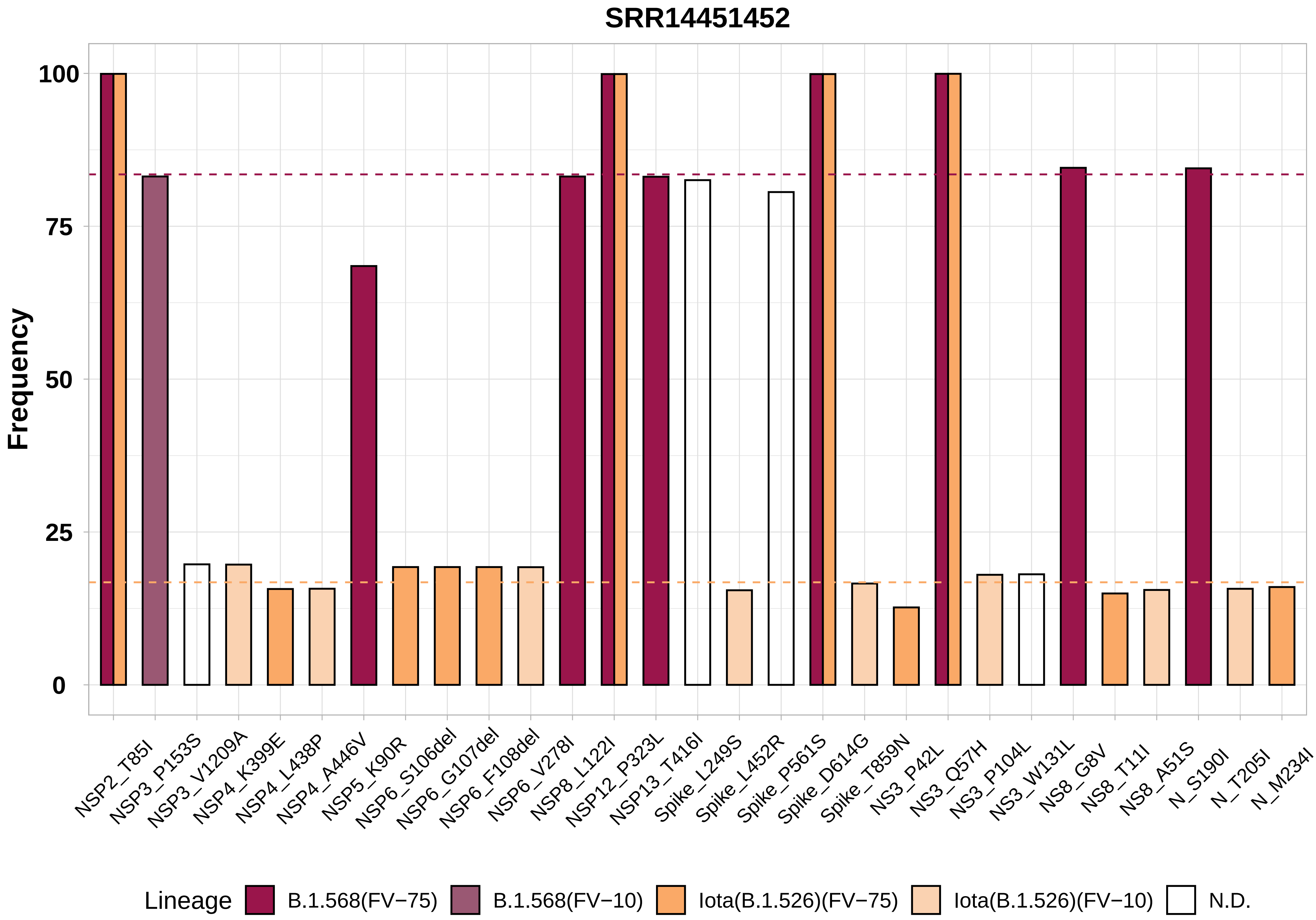

SRR14451452
Frequency
100
75
50
25
0
NSP2_T85I
NSP3_P153S
NSP3_V1209A
NSP4_K399E
NSP4_L438P
NSP4_A446V
NSP5_K90R
NSP6_S106del
NSP6_G107del
NSP6_F108del
NSP6_V278I
NSP8_L122I
NSP12_P323L
NSP13_T416I
Spike_L249S
Spike_L452R
Spike_P561S
Spike_D614G
Spike_T859N
NS3_P42L
NS3_Q57H
NS3_P104L
NS3_W131L
NS8_G8V
NS8_T11I
NS8_A51S
N_S190I
N_T205I
N_M234I
Lineage
B.1.568(FV−75)
B.1.568(FV−10)
Iota(B.1.526)(FV−75)
Iota(B.1.526)(FV−10)
N.D.

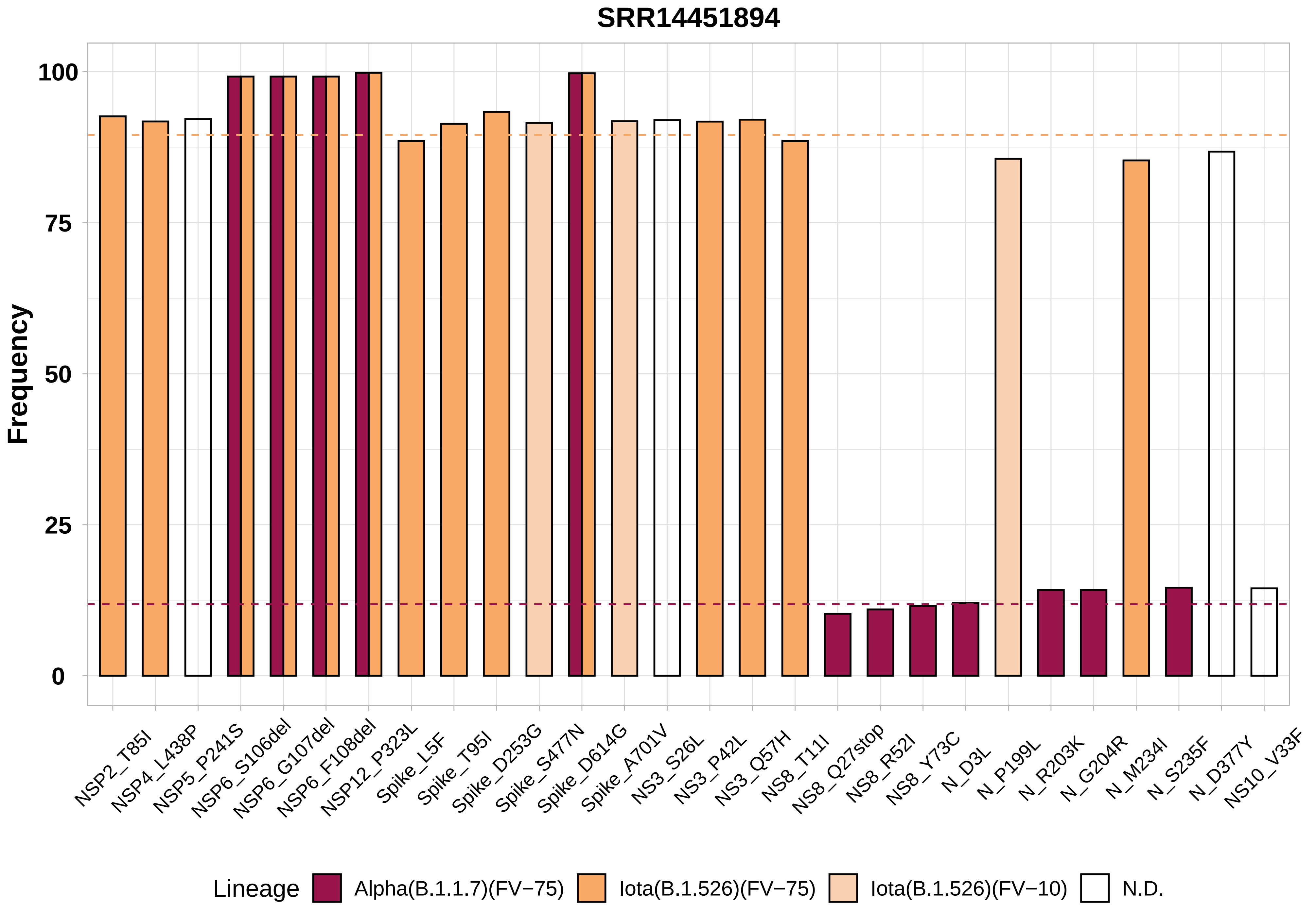

SRR14451894
Frequency
100
75
50
25
0
NSP2_T85I
NSP4_L438P
NSP5_P241S
NSP6_S106del
NSP6_G107del
NSP6_F108del
NSP12_P323L
Spike_L5F
Spike_T95I
Spike_D253G
Spike_S477N
Spike_D614G
Spike_A701V
NS3_S26L
NS3_P42L
NS3_Q57H
NS8_T11I
NS8_Q27stop
NS8_R52I
NS8_Y73C
N_D3L
N_P199L
N_R203K
N_G204R
N_M234I
N_S235F
N_D377Y
NS10_V33F
Lineage
Alpha(B.1.1.7)(FV−75)
Iota(B.1.526)(FV−75)
Iota(B.1.526)(FV−10)
N.D.

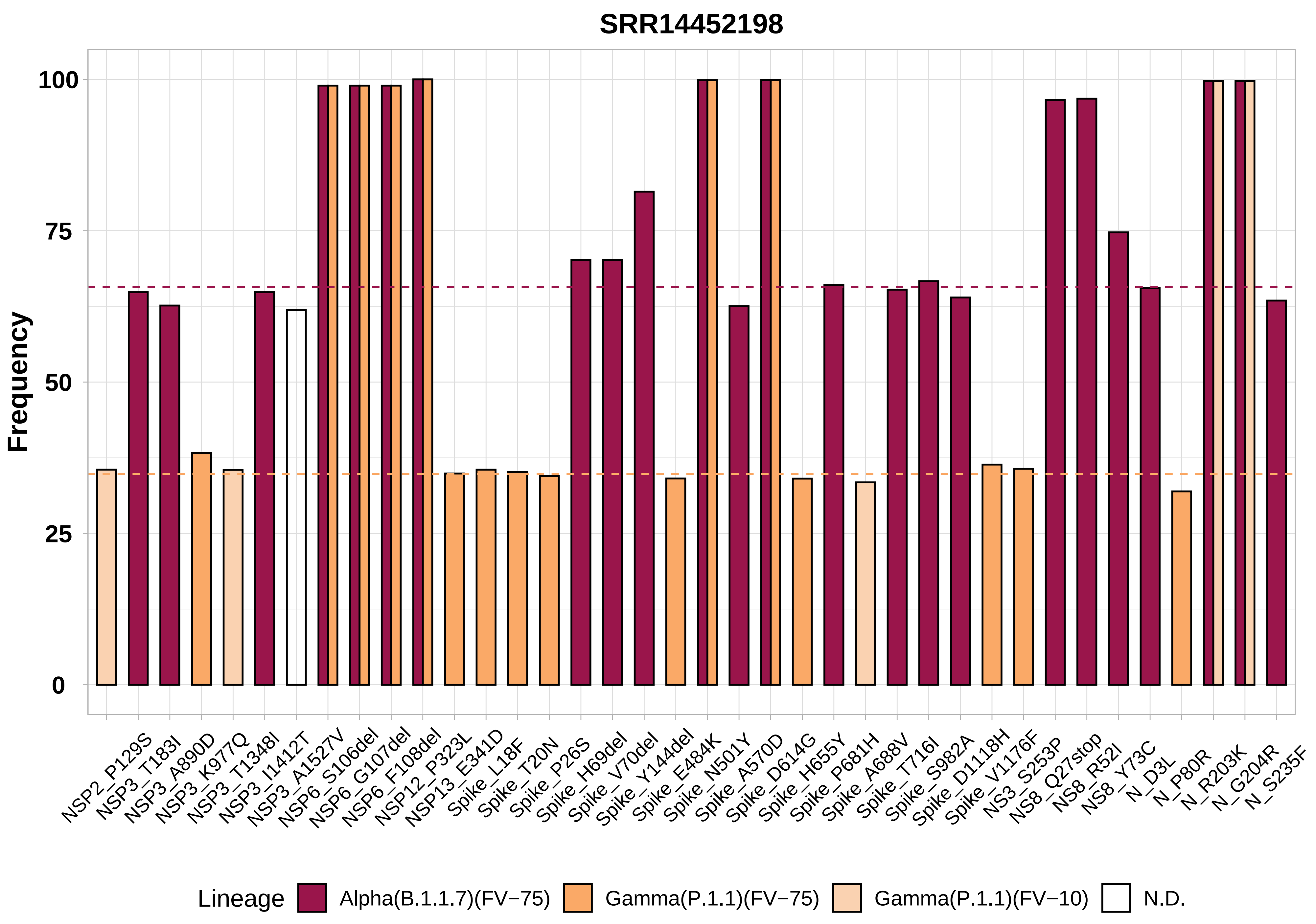

SRR14452198
Frequency
100
75
50
25
0
NSP2_P129S
NSP3_T183I
NSP3_A890D
NSP3_K977Q
NSP3_T1348I
NSP3_I1412T
NSP3_A1527V
NSP6_S106del
NSP6_G107del
NSP6_F108del
NSP12_P323L
NSP13_E341D
Spike_L18F
Spike_T20N
Spike_P26S
Spike_H69del
Spike_V70del
Spike_Y144del
Spike_E484K
Spike_N501Y
Spike_A570D
Spike_D614G
Spike_H655Y
Spike_P681H
Spike_A688V
Spike_T716I
Spike_S982A
Spike_D1118H
Spike_V1176F
NS3_S253P
NS8_Q27stop
NS8_R52I
NS8_Y73C
N_D3L
N_P80R
N_R203K
N_G204R
N_S235F
Lineage
Alpha(B.1.1.7)(FV−75)
Gamma(P.1.1)(FV−75)
Gamma(P.1.1)(FV−10)
N.D.

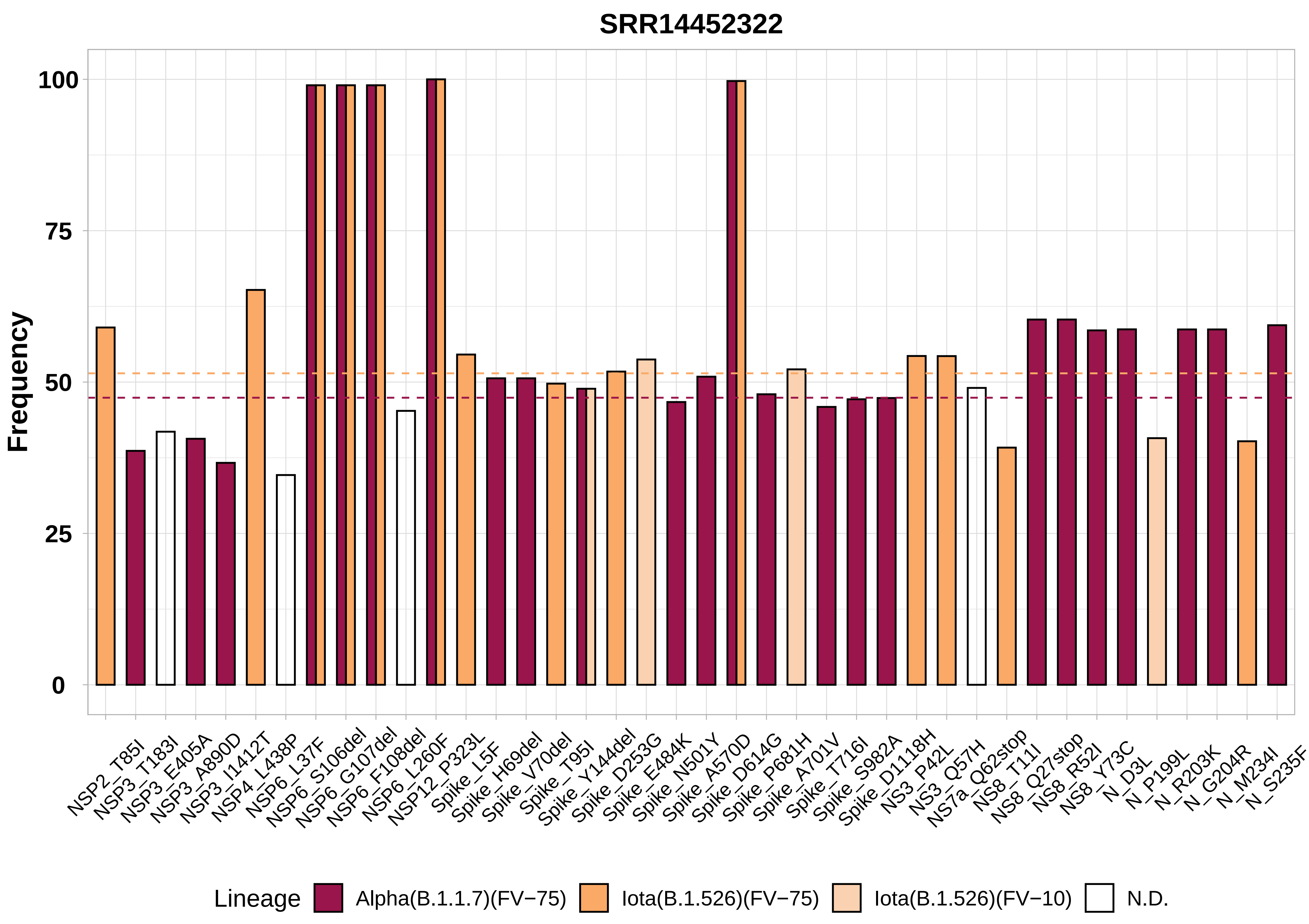

SRR14452322
Frequency
100
75
50
25
0
NSP2_T85I
NSP3_T183I
NSP3_E405A
NSP3_A890D
NSP3_I1412T
NSP4_L438P
NSP6_L37F
NSP6_S106del
NSP6_G107del
NSP6_F108del
NSP6_L260F
NSP12_P323L
Spike_L5F
Spike_H69del
Spike_V70del
Spike_T95I
Spike_Y144del
Spike_D253G
Spike_E484K
Spike_N501Y
Spike_A570D
Spike_D614G
Spike_P681H
Spike_A701V
Spike_T716I
Spike_S982A
Spike_D1118H
NS3_P42L
NS3_Q57H
NS7a_Q62stop
NS8_T11I
NS8_Q27stop
NS8_R52I
NS8_Y73C
N_D3L
N_P199L
N_R203K
N_G204R
N_M234I
N_S235F
Lineage
Alpha(B.1.1.7)(FV−75)
Iota(B.1.526)(FV−75)
Iota(B.1.526)(FV−10)
N.D.

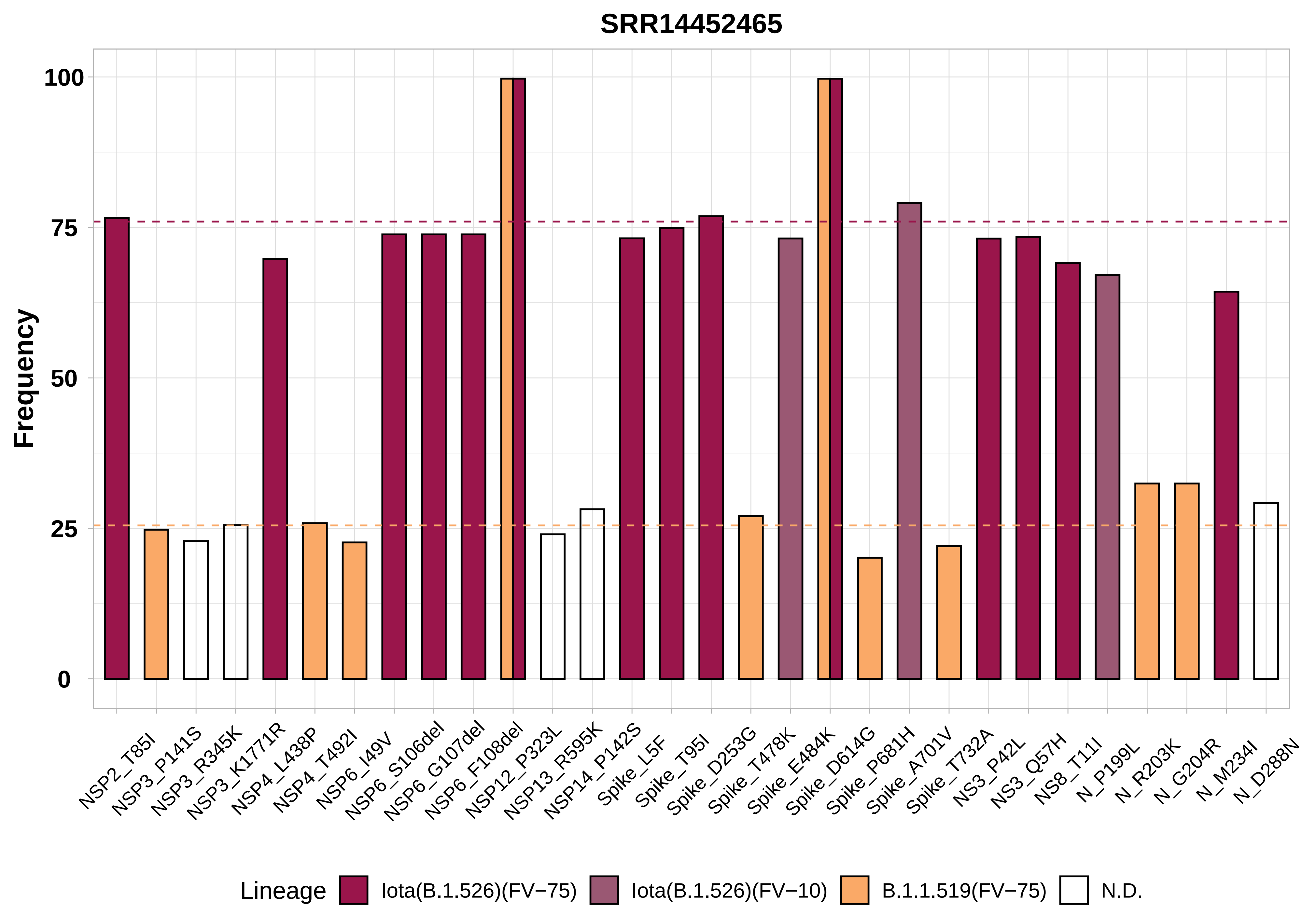
SRR14452465
Frequency
100
75
50
25
0
NSP2_T85I
NSP3_P141S
NSP3_R345K
NSP3_K1771R
NSP4_L438P
NSP4_T492I
NSP6_I49V
NSP6_S106del
NSP6_G107del
NSP6_F108del
NSP12_P323L
NSP13_R595K
NSP14_P142S
Spike_L5F
Spike_T95I
Spike_D253G
Spike_T478K
Spike_E484K
Spike_D614G
Spike_P681H
Spike_A701V
Spike_T732A
NS3_P42L
NS3_Q57H
NS8_T11I
N_P199L
N_R203K
N_G204R
N_M234I
N_D288N
Lineage
Iota(B.1.526)(FV−75)
Iota(B.1.526)(FV−10)
B.1.1.519(FV−75)
N.D.

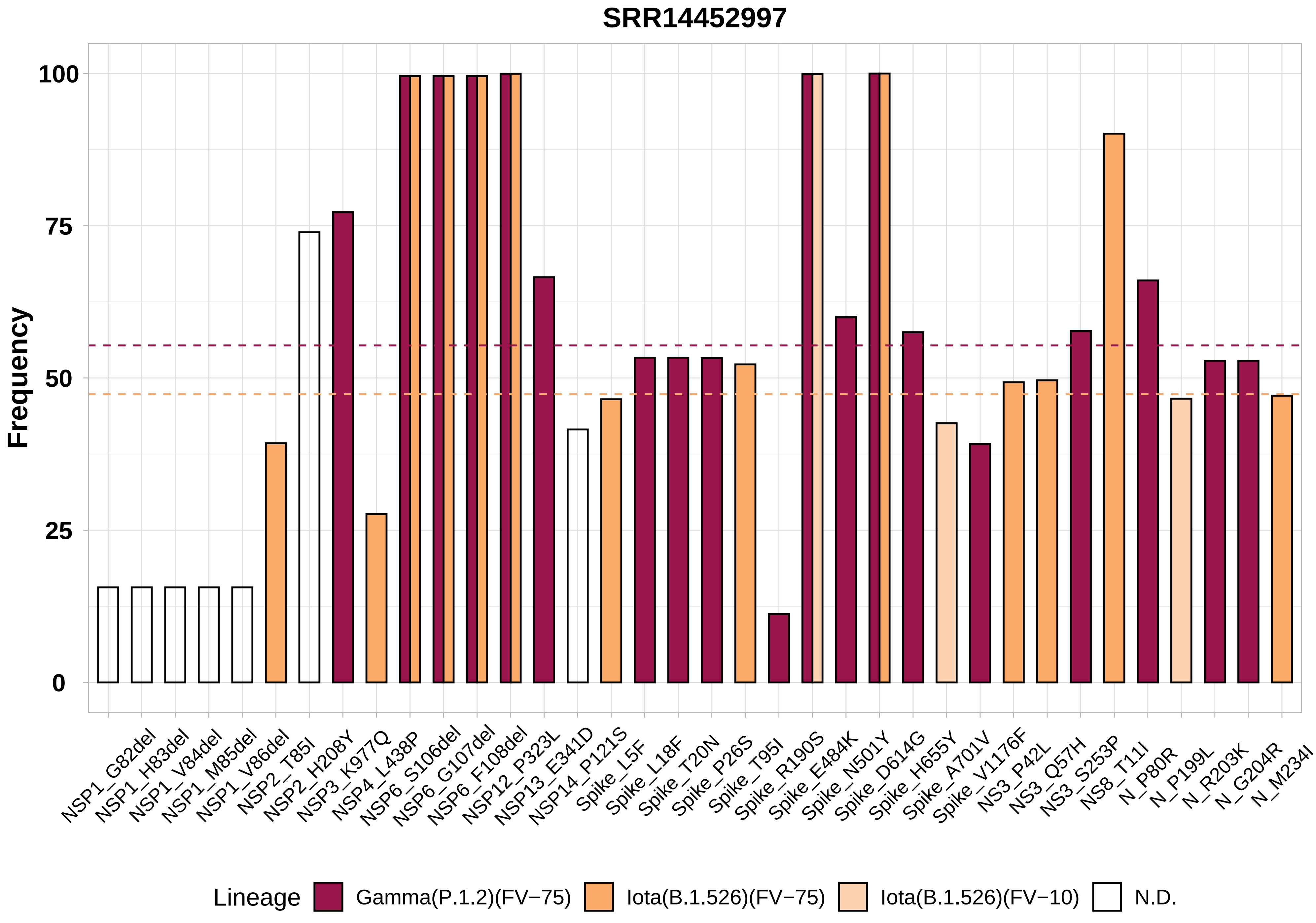
SRR14452997
Frequency
100
75
50
25
0
NSP1_G82del
NSP1_H83del
NSP1_V84del
NSP1_M85del
NSP1_V86del
NSP2_T85I
NSP2_H208Y
NSP3_K977Q
NSP4_L438P
NSP6_S106del
NSP6_G107del
NSP6_F108del
NSP12_P323L
NSP13_E341D
NSP14_P121S
Spike_L5F
Spike_L18F
Spike_T20N
Spike_P26S
Spike_T95I
Spike_R190S
Spike_E484K
Spike_N501Y
Spike_D614G
Spike_H655Y
Spike_A701V
Spike_V1176F
NS3_P42L
NS3_Q57H
NS3_S253P
NS8_T11I
N_P80R
N_P199L
N_R203K
N_G204R
N_M234I
Lineage
Gamma(P.1.2)(FV−75)
Iota(B.1.526)(FV−75)
Iota(B.1.526)(FV−10)
N.D.

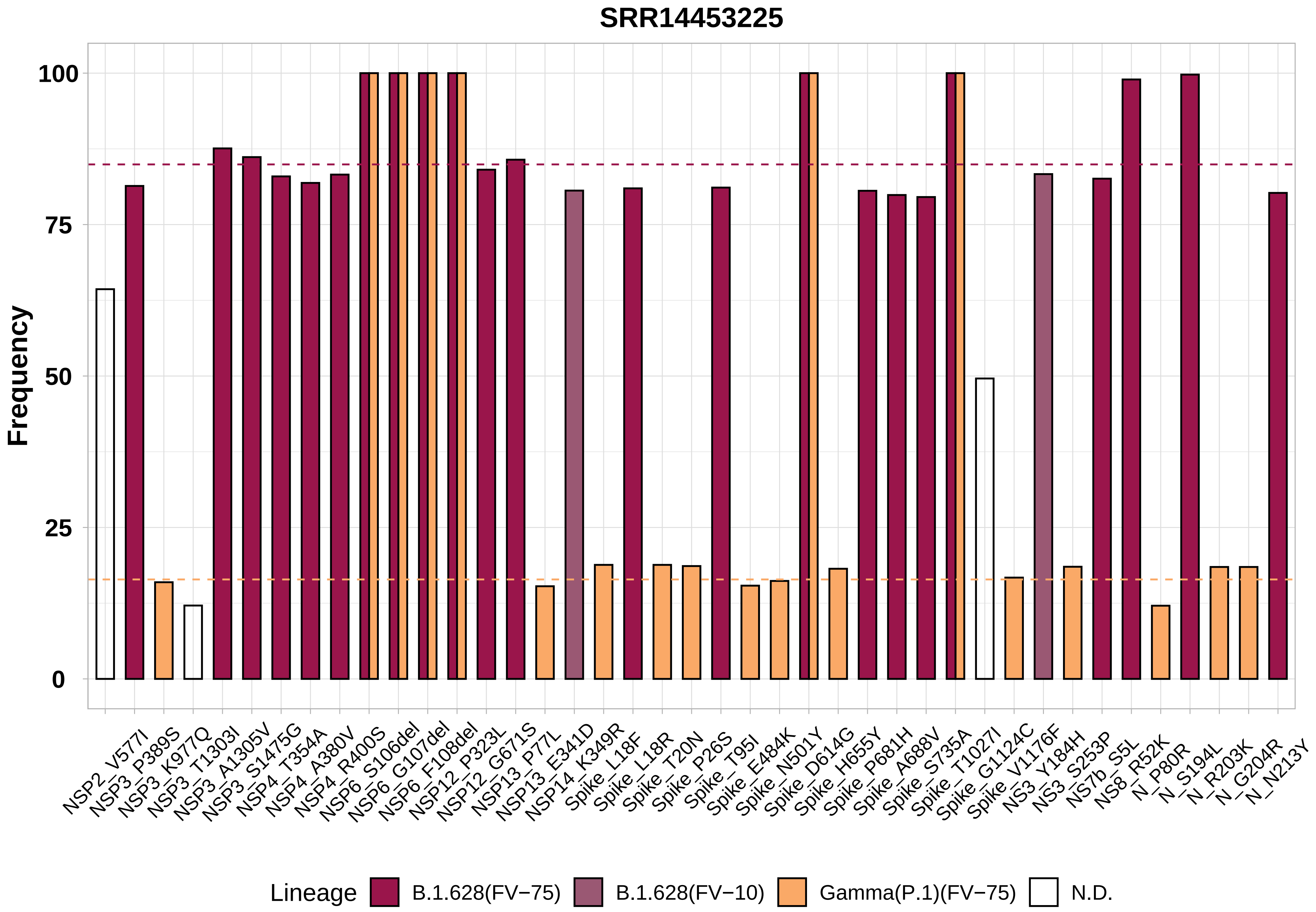
SRR14453225
Frequency
100
75
50
25
0
NSP2_V577I
NSP3_P389S
NSP3_K977Q
NSP3_T1303I
NSP3_A1305V
NSP3_S1475G
NSP4_T354A
NSP4_A380V
NSP4_R400S
NSP6_S106del
NSP6_G107del
NSP6_F108del
NSP12_P323L
NSP12_G671S
NSP13_P77L
NSP13_E341D
NSP14_K349R
Spike_L18F
Spike_L18R
Spike_T20N
Spike_P26S
Spike_T95I
Spike_E484K
Spike_N501Y
Spike_D614G
Spike_H655Y
Spike_P681H
Spike_A688V
Spike_S735A
Spike_T1027I
Spike_G1124C
Spike_V1176F
NS3_Y184H
NS3_S253P
NS7b_S5L
NS8_R52K
N_P80R
N_S194L
N_R203K
N_G204R
N_N213Y
Lineage
B.1.628(FV−75)
B.1.628(FV−10)
Gamma(P.1)(FV−75)
N.D.

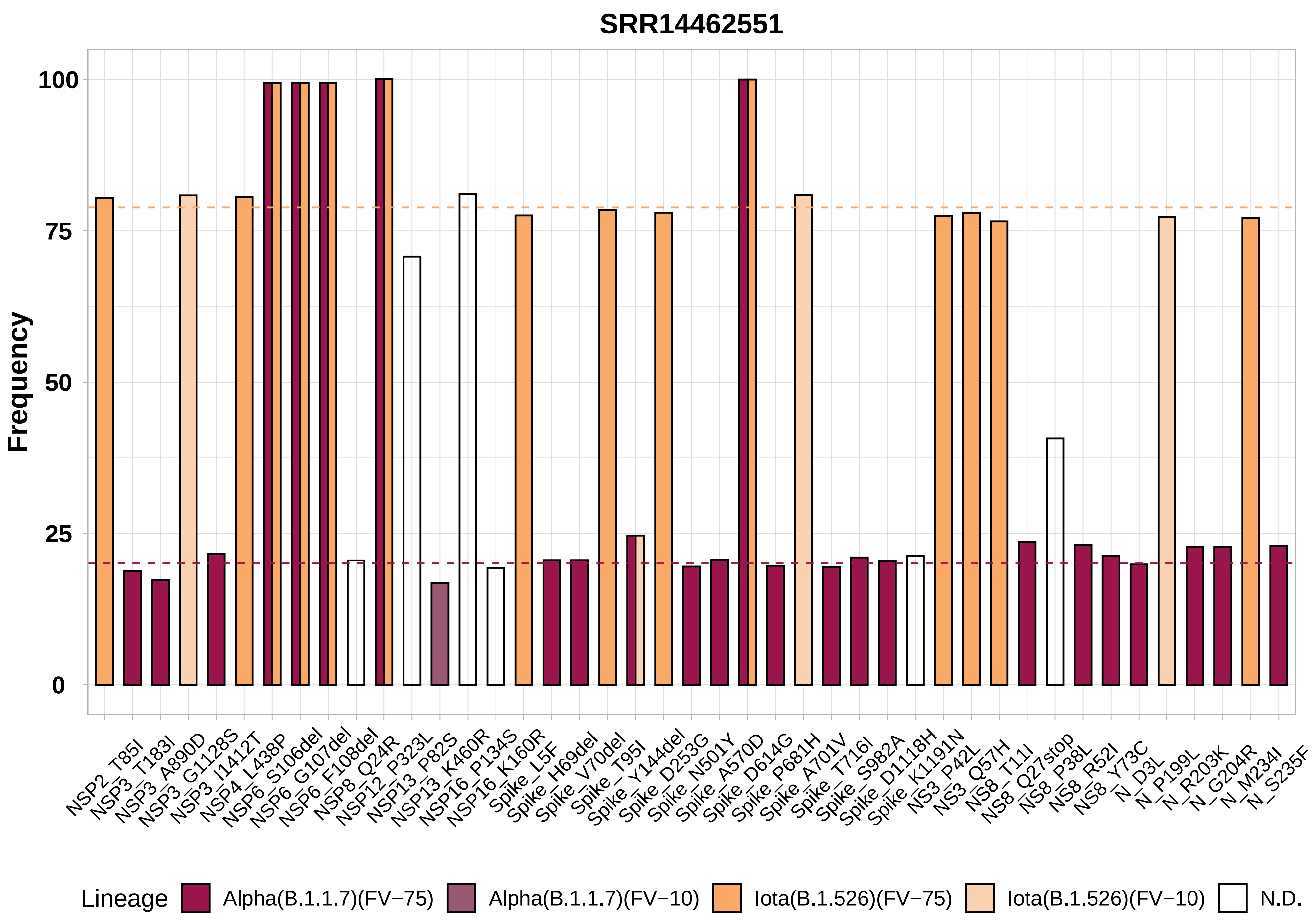

SRR14462551
Frequency
100
75
50
25
0
NSP2_T85I
NSP3_T183I
NSP3_A890D
NSP3_G1128S
NSP3_I1412T
NSP4_L438P
NSP6_S106del
NSP6_G107del
NSP6_F108del
NSP8_Q24R
NSP12_P323L
NSP13_P82S
NSP13_K460R
NSP16_P134S
NSP16_K160R
Spike_L5F
Spike_H69del
Spike_V70del
Spike_T95I
Spike_Y144del
Spike_D253G
Spike_N501Y
Spike_A570D
Spike_D614G
Spike_P681H
Spike_A701V
Spike_T716I
Spike_S982A
Spike_D1118H
Spike_K1191N
NS3_P42L
NS3_Q57H
NS8_T11I
NS8_Q27stop
NS8_P38L
NS8_R52I
NS8_Y73C
N_D3L
N_P199L
N_R203K
N_G204R
N_M234I
N_S235F
Lineage
Alpha(B.1.1.7)(FV−75)
Alpha(B.1.1.7)(FV−10)
Iota(B.1.526)(FV−75)
Iota(B.1.526)(FV−10)
N.D.

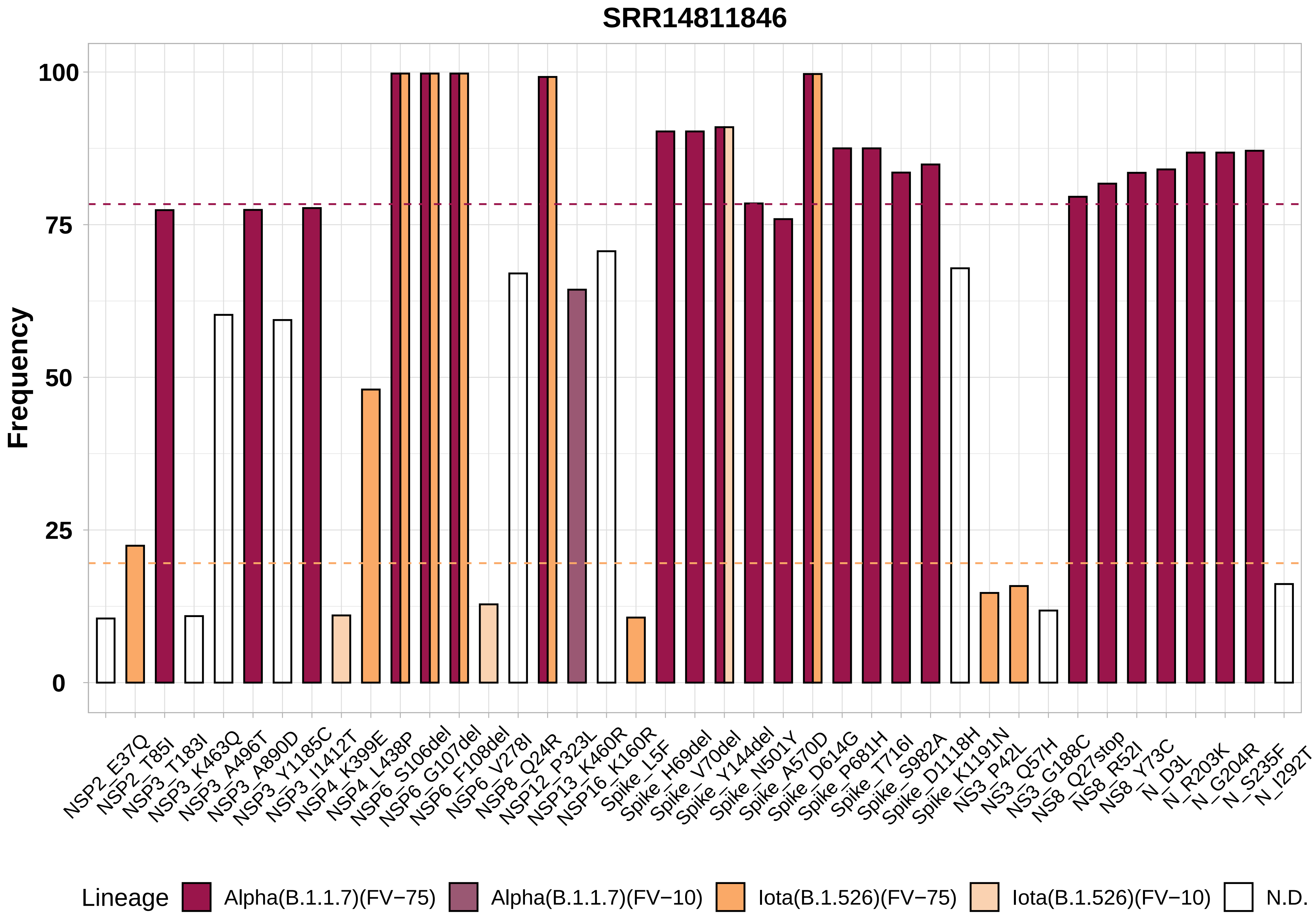
SRR14811846
Frequency
100
75
50
25
0
NSP2_E37Q
NSP2_T85I
NSP3_T183I
NSP3_K463Q
NSP3_A496T
NSP3_A890D
NSP3_Y1185C
NSP3_I1412T
NSP4_K399E
NSP4_L438P
NSP6_S106del
NSP6_G107del
NSP6_F108del
NSP6_V278I
NSP8_Q24R
NSP12_P323L
NSP13_K460R
NSP16_K160R
Spike_L5F
Spike_H69del
Spike_V70del
Spike_Y144del
Spike_N501Y
Spike_A570D
Spike_D614G
Spike_P681H
Spike_T716I
Spike_S982A
Spike_D1118H
Spike_K1191N
NS3_P42L
NS3_Q57H
NS3_G188C
NS8_Q27stop
NS8_R52I
NS8_Y73C
N_D3L
N_R203K
N_G204R
N_S235F
N_I292T
Lineage
Alpha(B.1.1.7)(FV−75)
Alpha(B.1.1.7)(FV−10)
Iota(B.1.526)(FV−75)
Iota(B.1.526)(FV−10)
N.D.

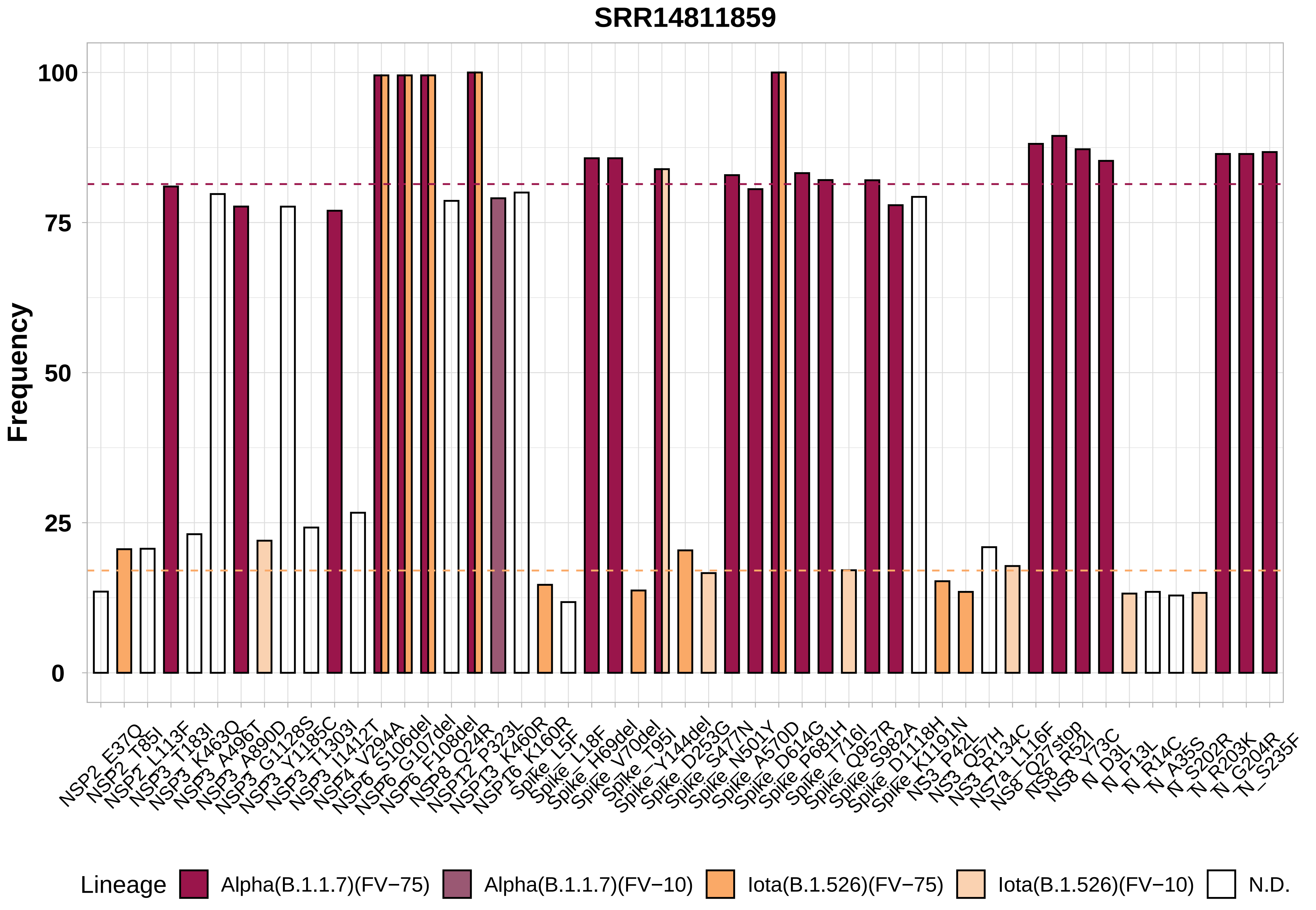
SRR14811859
Frequency
100
75
50
25
0
NSP2_E37Q
NSP2_T85I
NSP2_L113F
NSP3_T183I
NSP3_K463Q
NSP3_A496T
NSP3_A890D
NSP3_G1128S
NSP3_Y1185C
NSP3_T1303I
NSP3_I1412T
NSP4_V294A
NSP6_S106del
NSP6_G107del
NSP6_F108del
NSP8_Q24R
NSP12_P323L
NSP13_K460R
NSP16_K160R
Spike_L5F
Spike_L18F
Spike_H69del
Spike_V70del
Spike_T95I
Spike_Y144del
Spike_D253G
Spike_S477N
Spike_N501Y
Spike_A570D
Spike_D614G
Spike_P681H
Spike_T716I
Spike_Q957R
Spike_S982A
Spike_D1118H
Spike_K1191N
NS3_P42L
NS3_Q57H
NS3_R134C
NS7a_L116F
NS8_Q27stop
NS8_R52I
NS8_Y73C
N_D3L
N_P13L
N_R14C
N_A35S
N_S202R
N_R203K
N_G204R
N_S235F
Lineage
Alpha(B.1.1.7)(FV−75)
Alpha(B.1.1.7)(FV−10)
Iota(B.1.526)(FV−75)
Iota(B.1.526)(FV−10)
N.D.

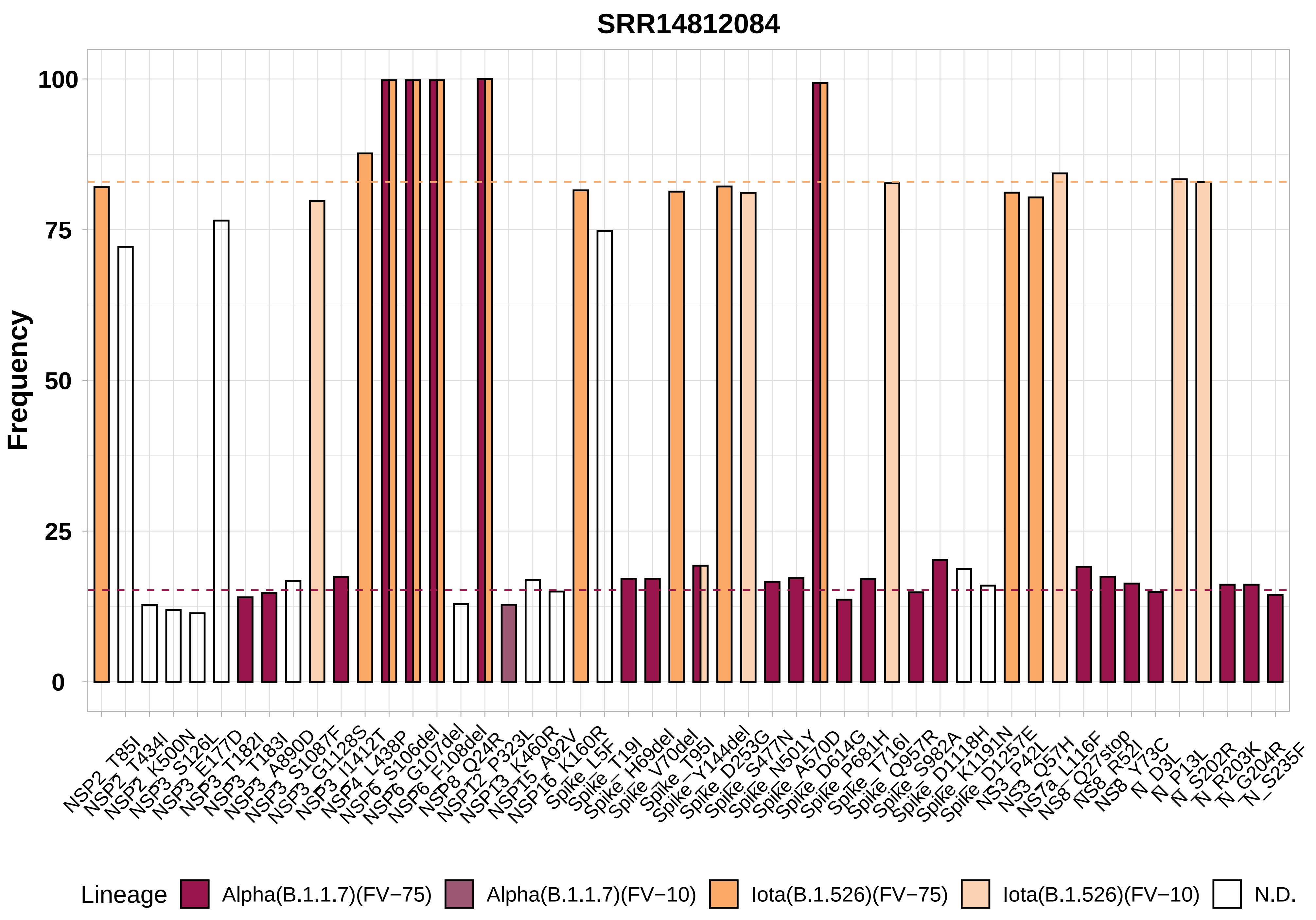

SRR14812084
Frequency
100
75
50
25
0
NSP2_T85I
NSP2_T434I
NSP2_K500N
NSP3_S126L
NSP3_E177D
NSP3_T182I
NSP3_T183I
NSP3_A890D
NSP3_S1087F
NSP3_G1128S
NSP3_I1412T
NSP4_L438P
NSP6_S106del
NSP6_G107del
NSP6_F108del
NSP8_Q24R
NSP12_P323L
NSP13_K460R
NSP15_A92V
NSP16_K160R
Spike_L5F
Spike_T19I
Spike_H69del
Spike_V70del
Spike_T95I
Spike_Y144del
Spike_D253G
Spike_S477N
Spike_N501Y
Spike_A570D
Spike_D614G
Spike_P681H
Spike_T716I
Spike_Q957R
Spike_S982A
Spike_D1118H
Spike_K1191N
Spike_D1257E
NS3_P42L
NS3_Q57H
NS7a_L116F
NS8_Q27stop
NS8_R52I
NS8_Y73C
N_D3L
N_P13L
N_S202R
N_R203K
N_G204R
N_S235F
Lineage
Alpha(B.1.1.7)(FV−75)
Alpha(B.1.1.7)(FV−10)
Iota(B.1.526)(FV−75)
Iota(B.1.526)(FV−10)
N.D.

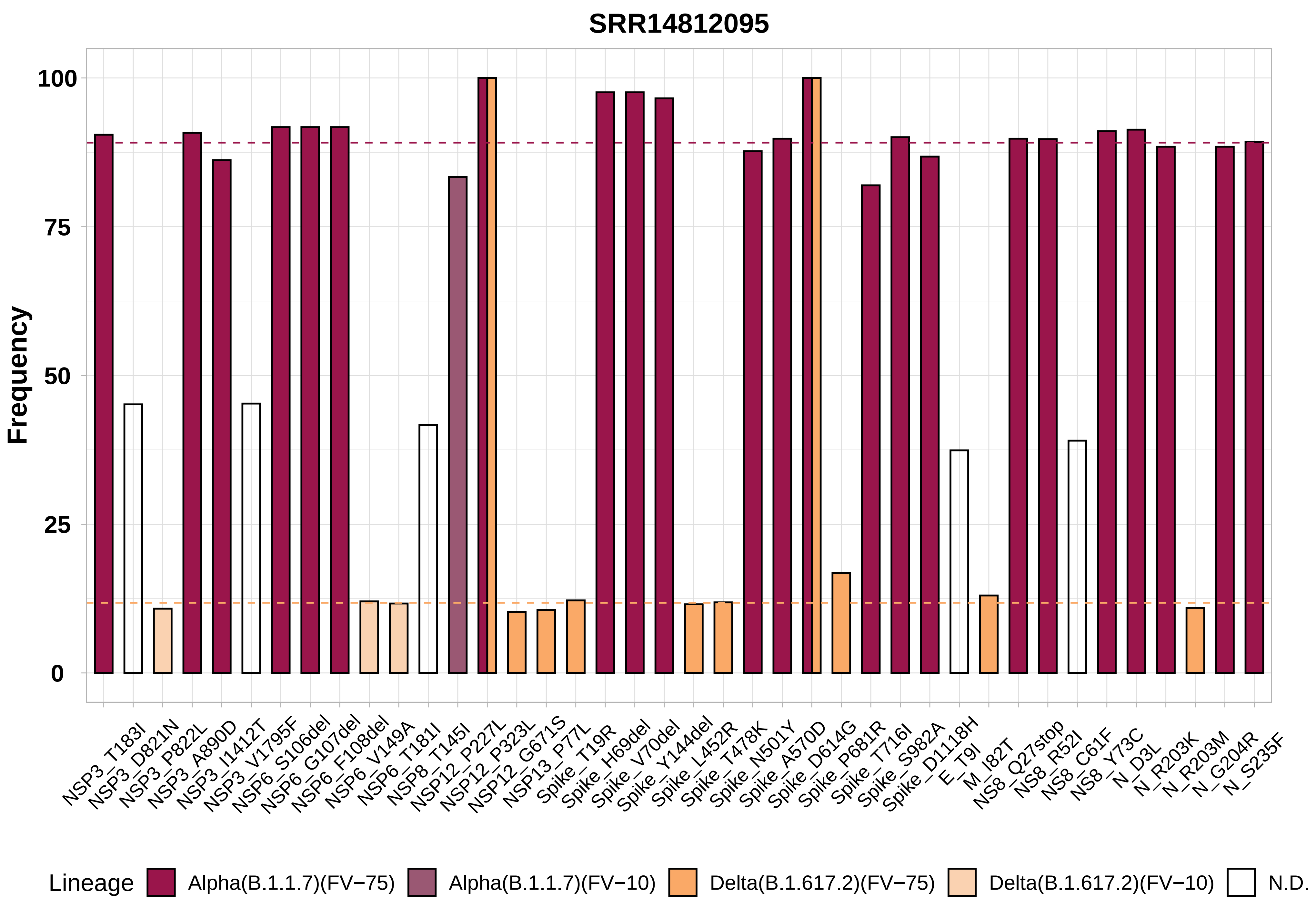

SRR14812095
Frequency
100
75
50
25
0
NSP3_T183I
NSP3_D821N
NSP3_P822L
NSP3_A890D
NSP3_I1412T
NSP3_V1795F
NSP6_S106del
NSP6_G107del
NSP6_F108del
NSP6_V149A
NSP6_T181I
NSP8_T145I
NSP12_P227L
NSP12_P323L
NSP12_G671S
NSP13_P77L
Spike_T19R
Spike_H69del
Spike_V70del
Spike_Y144del
Spike_L452R
Spike_T478K
Spike_N501Y
Spike_A570D
Spike_D614G
Spike_P681R
Spike_T716I
Spike_S982A
Spike_D1118H
E_T9I
M_I82T
NS8_Q27stop
NS8_R52I
NS8_C61F
NS8_Y73C
N_D3L
N_R203K
N_R203M
N_G204R
N_S235F
Lineage
Alpha(B.1.1.7)(FV−75)
Alpha(B.1.1.7)(FV−10)
Delta(B.1.617.2)(FV−75)
Delta(B.1.617.2)(FV−10)
N.D.

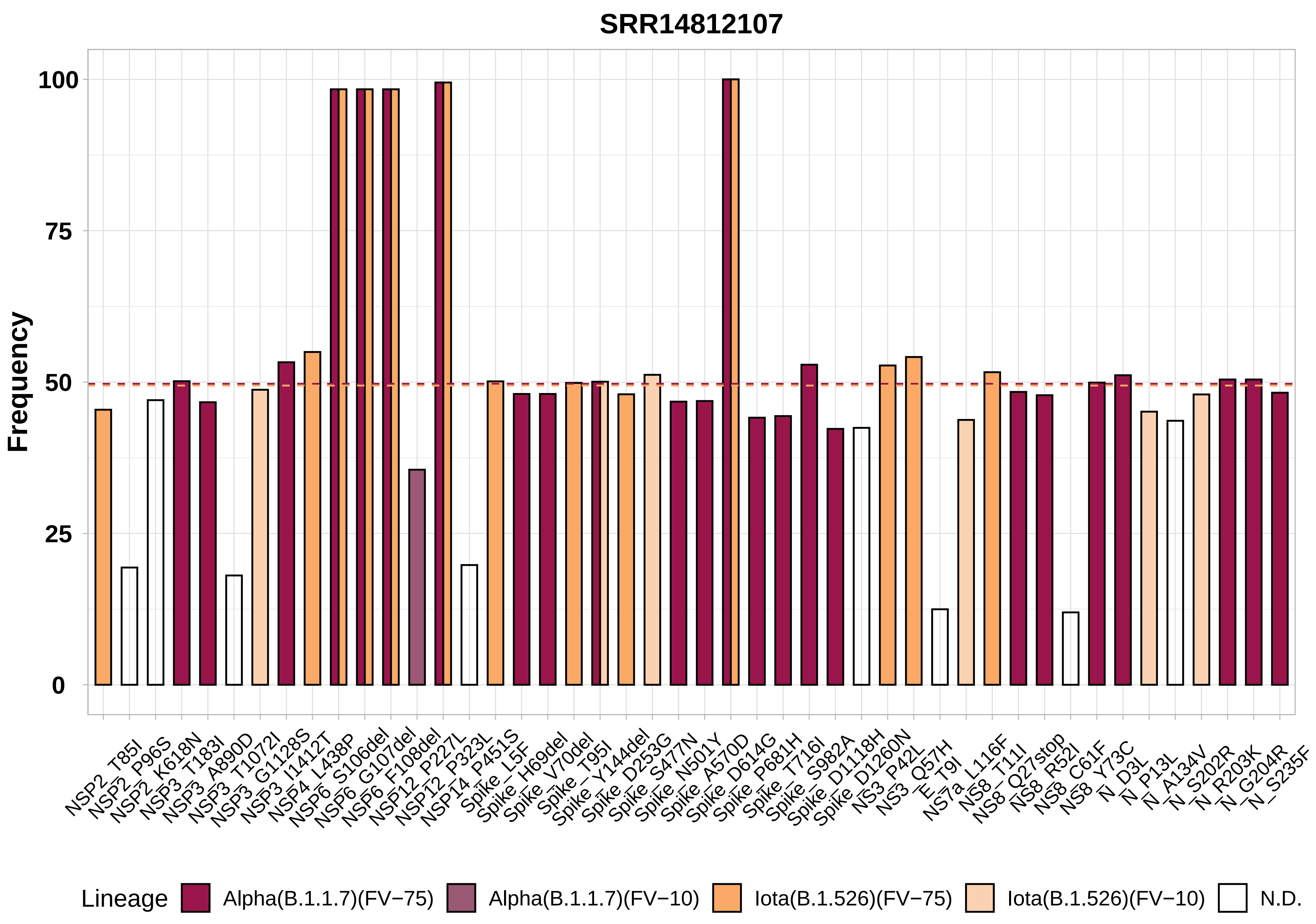
SRR14812107
Frequency
100
75
50
25
0
NSP2_T85I
NSP2_P96S
NSP2_K618N
NSP3_T183I
NSP3_A890D
NSP3_T1072I
NSP3_G1128S
NSP3_I1412T
NSP4_L438P
NSP6_S106del
NSP6_G107del
NSP6_F108del
NSP12_P227L
NSP12_P323L
NSP14_P451S
Spike_L5F
Spike_H69del
Spike_V70del
Spike_T95I
Spike_Y144del
Spike_D253G
Spike_S477N
Spike_N501Y
Spike_A570D
Spike_D614G
Spike_P681H
Spike_T716I
Spike_S982A
Spike_D1118H
Spike_D1260N
NS3_P42L
NS3_Q57H
E_T9I
NS7a_L116F
NS8_T11I
NS8_Q27stop
NS8_R52I
NS8_C61F
NS8_Y73C
N_D3L
N_P13L
N_A134V
N_S202R
N_R203K
N_G204R
N_S235F
Lineage
Alpha(B.1.1.7)(FV−75)
Alpha(B.1.1.7)(FV−10)
Iota(B.1.526)(FV−75)
Iota(B.1.526)(FV−10)
N.D.

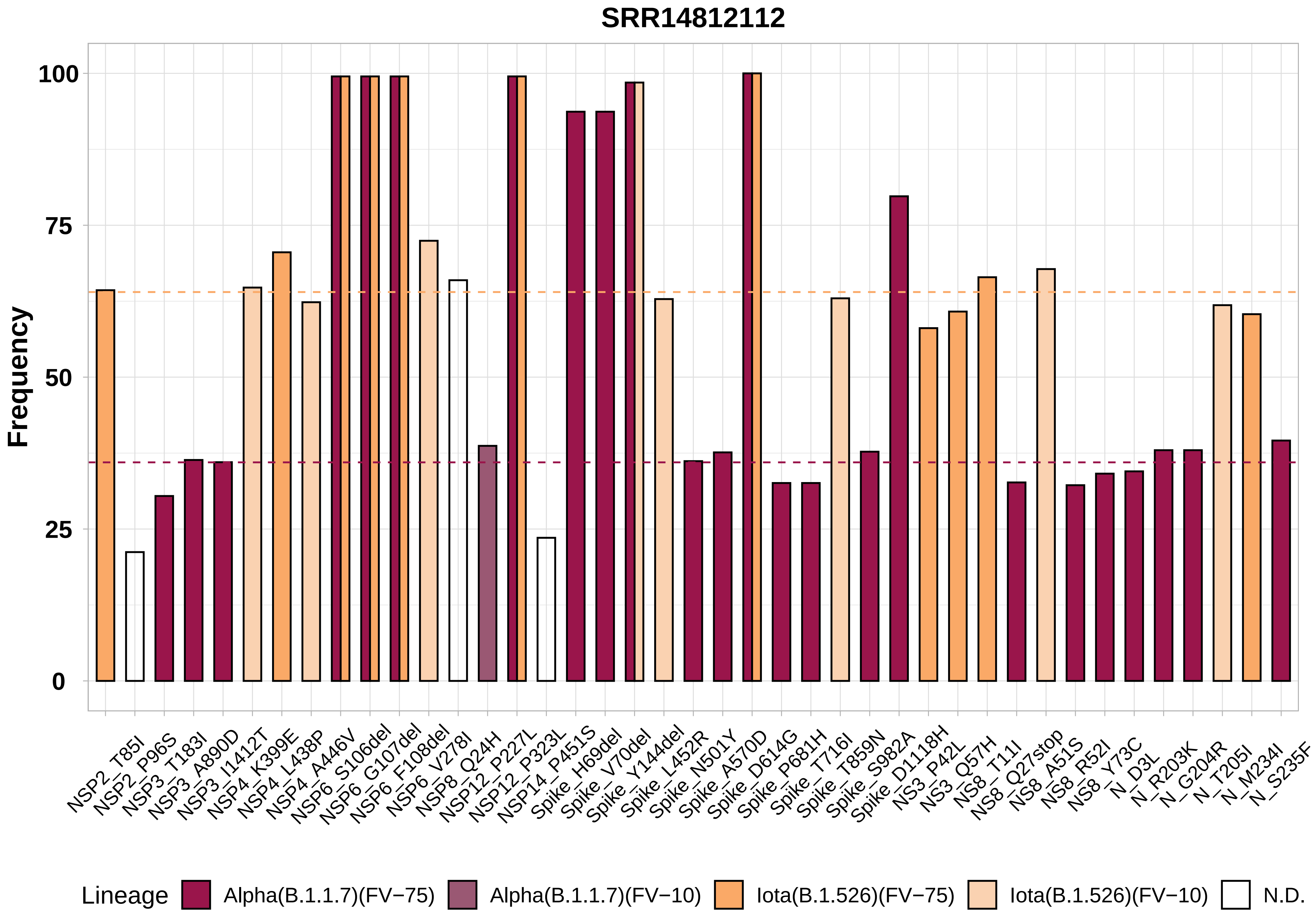
SRR14812112
Frequency
100
75
50
25
0
NSP2_T85I
NSP2_P96S
NSP3_T183I
NSP3_A890D
NSP3_I1412T
NSP4_K399E
NSP4_L438P
NSP4_A446V
NSP6_S106del
NSP6_G107del
NSP6_F108del
NSP6_V278I
NSP8_Q24H
NSP12_P227L
NSP12_P323L
NSP14_P451S
Spike_H69del
Spike_V70del
Spike_Y144del
Spike_L452R
Spike_N501Y
Spike_A570D
Spike_D614G
Spike_P681H
Spike_T716I
Spike_T859N
Spike_S982A
Spike_D1118H
NS3_P42L
NS3_Q57H
NS8_T11I
NS8_Q27stop
NS8_A51S
NS8_R52I
NS8_Y73C
N_D3L
N_R203K
N_G204R
N_T205I
N_M234I
N_S235F
Lineage
Alpha(B.1.1.7)(FV−75)
Alpha(B.1.1.7)(FV−10)
Iota(B.1.526)(FV−75)
Iota(B.1.526)(FV−10)
N.D.

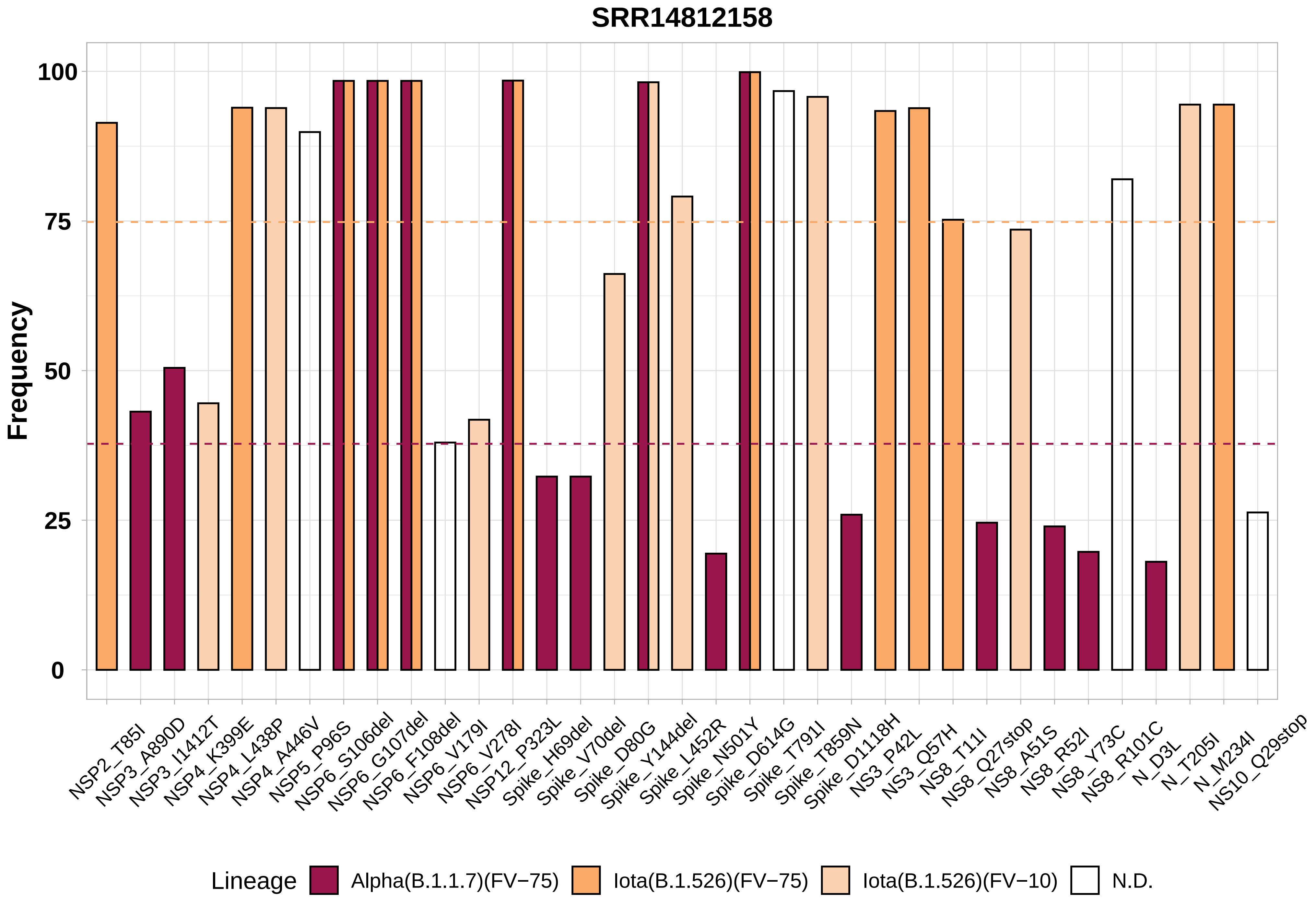

SRR14812158
Frequency
100
75
50
25
0
NSP2_T85I
NSP3_A890D
NSP3_I1412T
NSP4_K399E
NSP4_L438P
NSP4_A446V
NSP5_P96S
NSP6_S106del
NSP6_G107del
NSP6_F108del
NSP6_V179I
NSP6_V278I
NSP12_P323L
Spike_H69del
Spike_V70del
Spike_D80G
Spike_Y144del
Spike_L452R
Spike_N501Y
Spike_D614G
Spike_T791I
Spike_T859N
Spike_D1118H
NS3_P42L
NS3_Q57H
NS8_T11I
NS8_Q27stop
NS8_A51S
NS8_R52I
NS8_Y73C
NS8_R101C
N_D3L
N_T205I
N_M234I
NS10_Q29stop
Lineage
Alpha(B.1.1.7)(FV−75)
Iota(B.1.526)(FV−75)
Iota(B.1.526)(FV−10)
N.D.

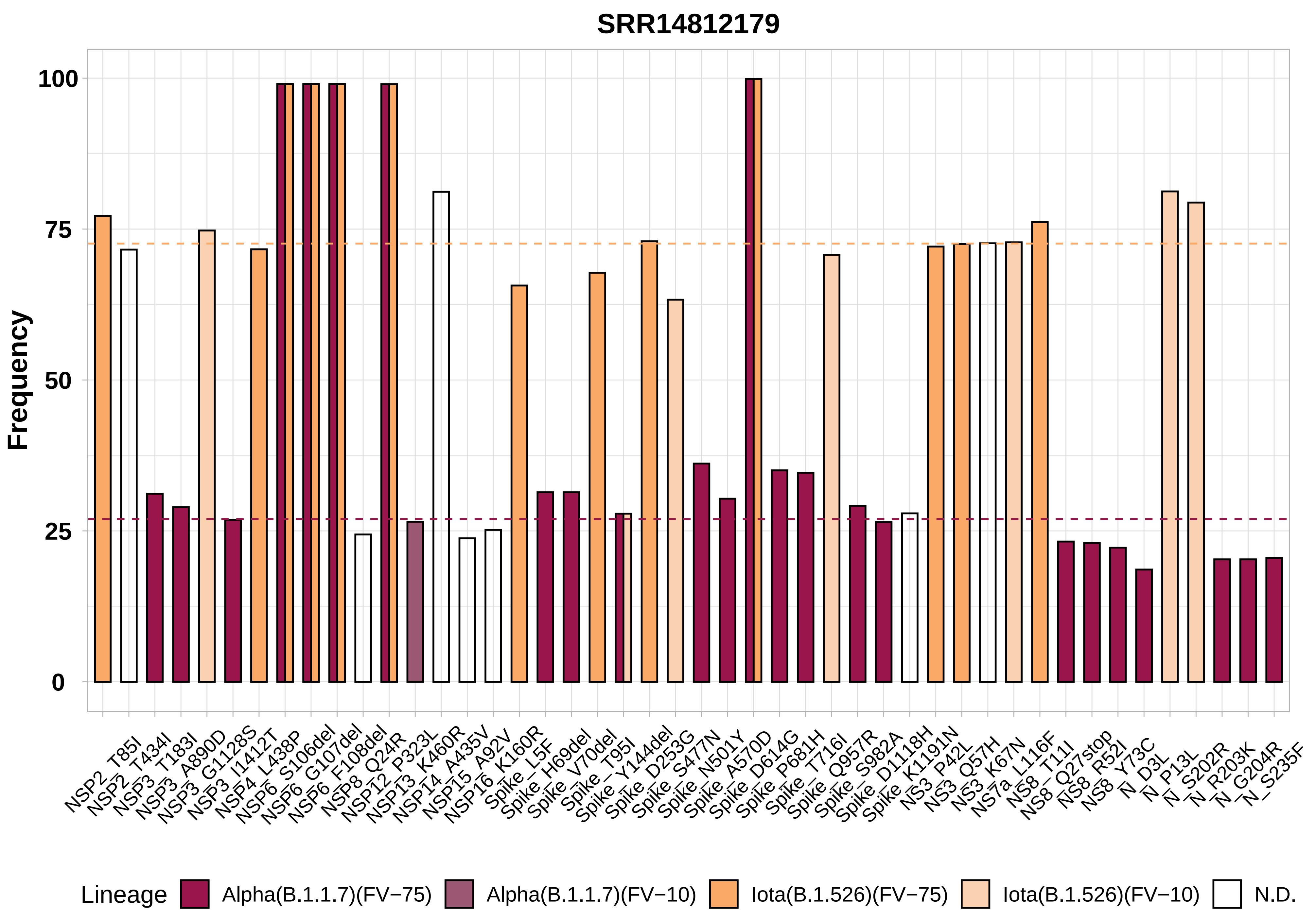

SRR14812179
Frequency
100
75
50
25
0
NSP2_T85I
NSP2_T434I
NSP3_T183I
NSP3_A890D
NSP3_G1128S
NSP3_I1412T
NSP4_L438P
NSP6_S106del
NSP6_G107del
NSP6_F108del
NSP8_Q24R
NSP12_P323L
NSP13_K460R
NSP14_A435V
NSP15_A92V
NSP16_K160R
Spike_L5F
Spike_H69del
Spike_V70del
Spike_T95I
Spike_Y144del
Spike_D253G
Spike_S477N
Spike_N501Y
Spike_A570D
Spike_D614G
Spike_P681H
Spike_T716I
Spike_Q957R
Spike_S982A
Spike_D1118H
Spike_K1191N
NS3_P42L
NS3_Q57H
NS3_K67N
NS7a_L116F
NS8_T11I
NS8_Q27stop
NS8_R52I
NS8_Y73C
N_D3L
N_P13L
N_S202R
N_R203K
N_G204R
N_S235F
Lineage
Alpha(B.1.1.7)(FV−75)
Alpha(B.1.1.7)(FV−10)
Iota(B.1.526)(FV−75)
Iota(B.1.526)(FV−10)
N.D.

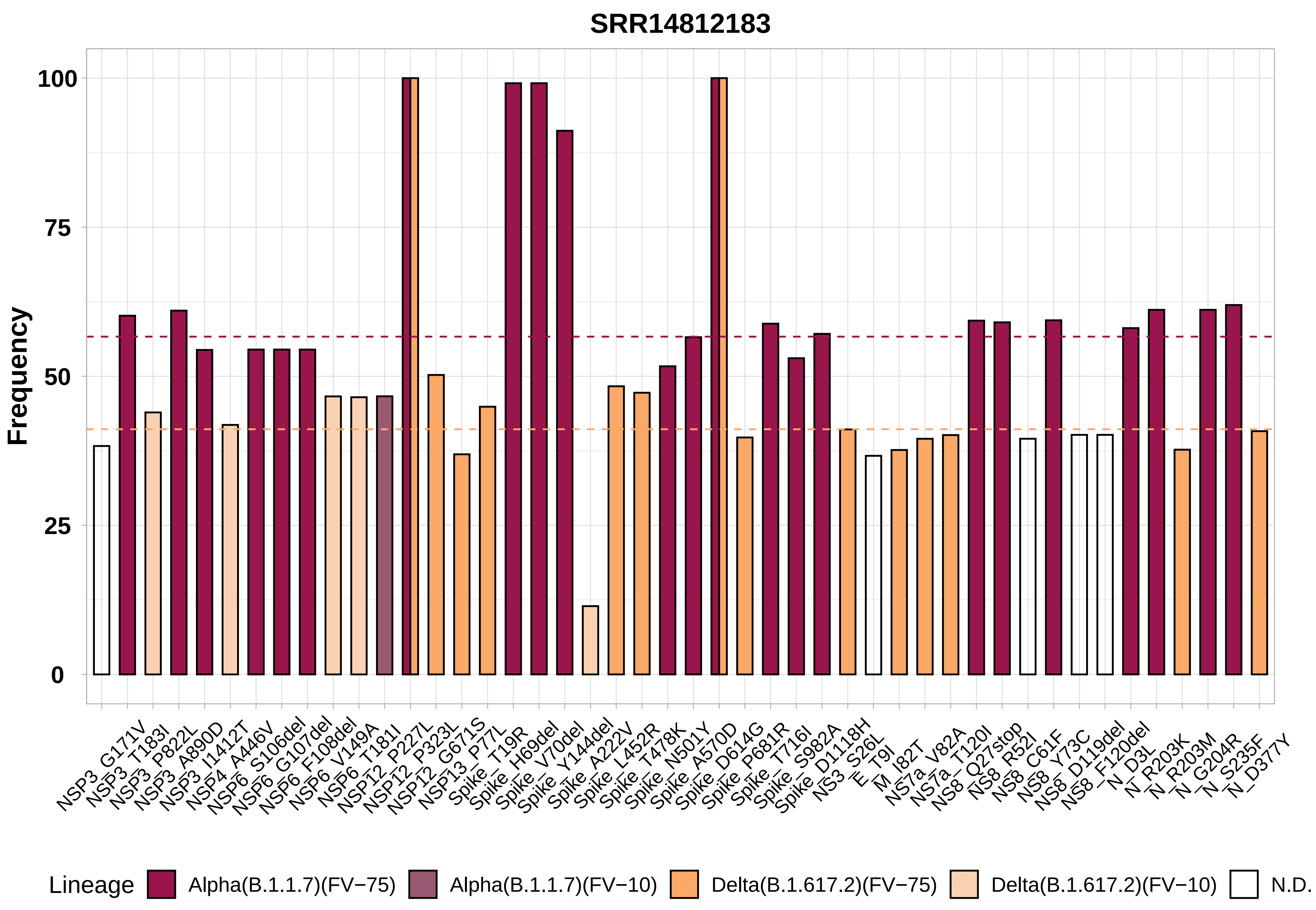
SRR14812183
Frequency
100
75
50
25
0
NSP3_G171V
NSP3_T183I
NSP3_P822L
NSP3_A890D
NSP3_I1412T
NSP4_A446V
NSP6_S106del
NSP6_G107del
NSP6_F108del
NSP6_V149A
NSP6_T181I
NSP12_P227L
NSP12_P323L
NSP12_G671S
NSP13_P77L
Spike_T19R
Spike_H69del
Spike_V70del
Spike_Y144del
Spike_A222V
Spike_L452R
Spike_T478K
Spike_N501Y
Spike_A570D
Spike_D614G
Spike_P681R
Spike_T716I
Spike_S982A
Spike_D1118H
NS3_S26L
E_T9I
M_I82T
NS7a_V82A
NS7a_T120I
NS8_Q27stop
NS8_R52I
NS8_C61F
NS8_Y73C
NS8_D119del
NS8_F120del
N_D3L
N_R203K
N_R203M
N_G204R
N_S235F
N_D377Y
Lineage
Alpha(B.1.1.7)(FV−75)
Alpha(B.1.1.7)(FV−10)
Delta(B.1.617.2)(FV−75)
Delta(B.1.617.2)(FV−10)
N.D.

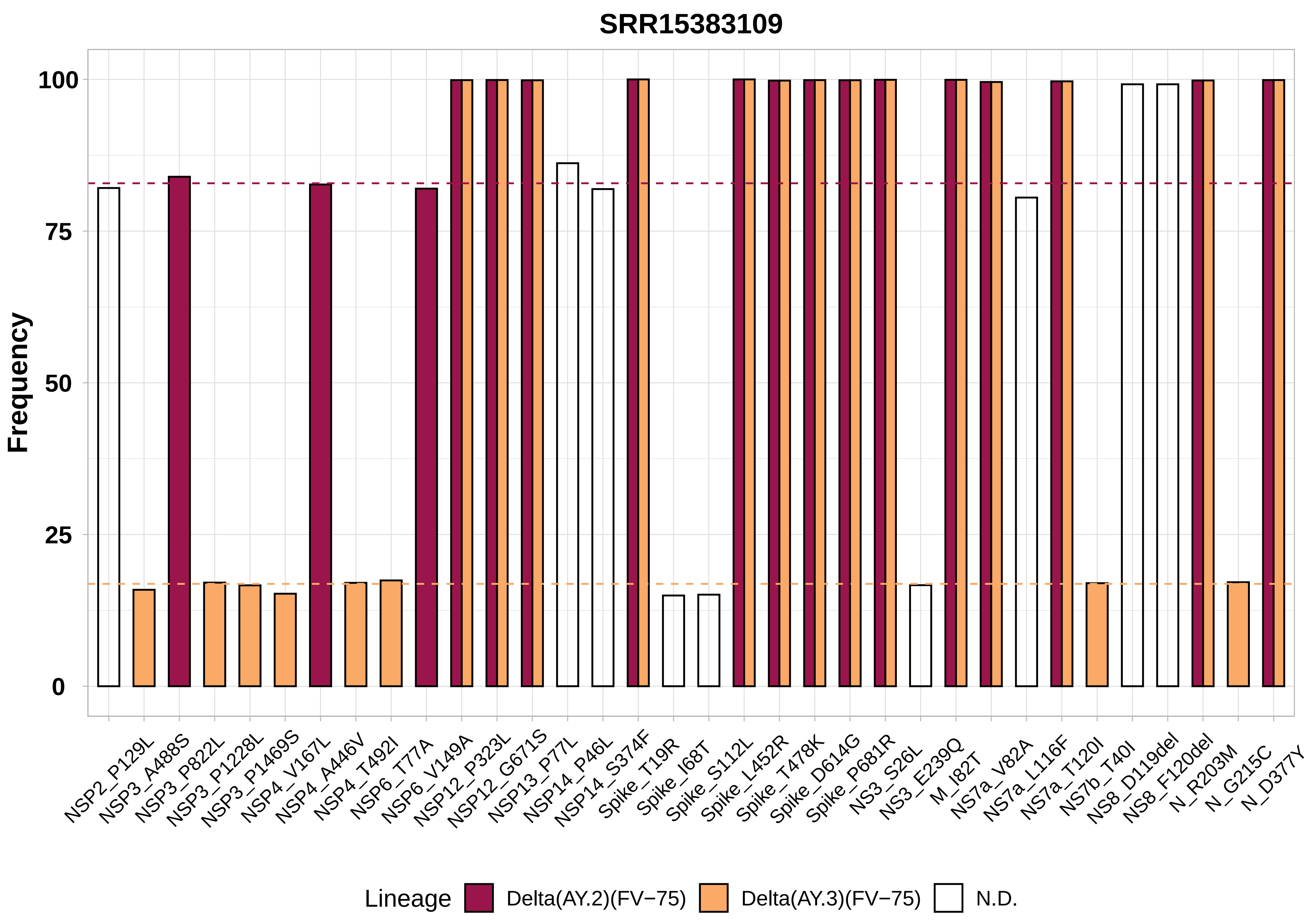
SRR15383109
Frequency
100
75
50
25
0
NSP2_P129L
NSP3_A488S
NSP3_P822L
NSP3_P1228L
NSP3_P1469S
NSP4_V167L
NSP4_A446V
NSP4_T492I
NSP6_T77A
NSP6_V149A
NSP12_P323L
NSP12_G671S
NSP13_P77L
NSP14_P46L
NSP14_S374F
Spike_T19R
Spike_I68T
Spike_S112L
Spike_L452R
Spike_T478K
Spike_D614G
Spike_P681R
NS3_S26L
NS3_E239Q
M_I82T
NS7a_V82A
NS7a_L116F
NS7a_T120I
NS7b_T40I
NS8_D119del
NS8_F120del
N_R203M
N_G215C
N_D377Y
Lineage
Delta(AY.2)(FV−75)
Delta(AY.3)(FV−75)
N.D.

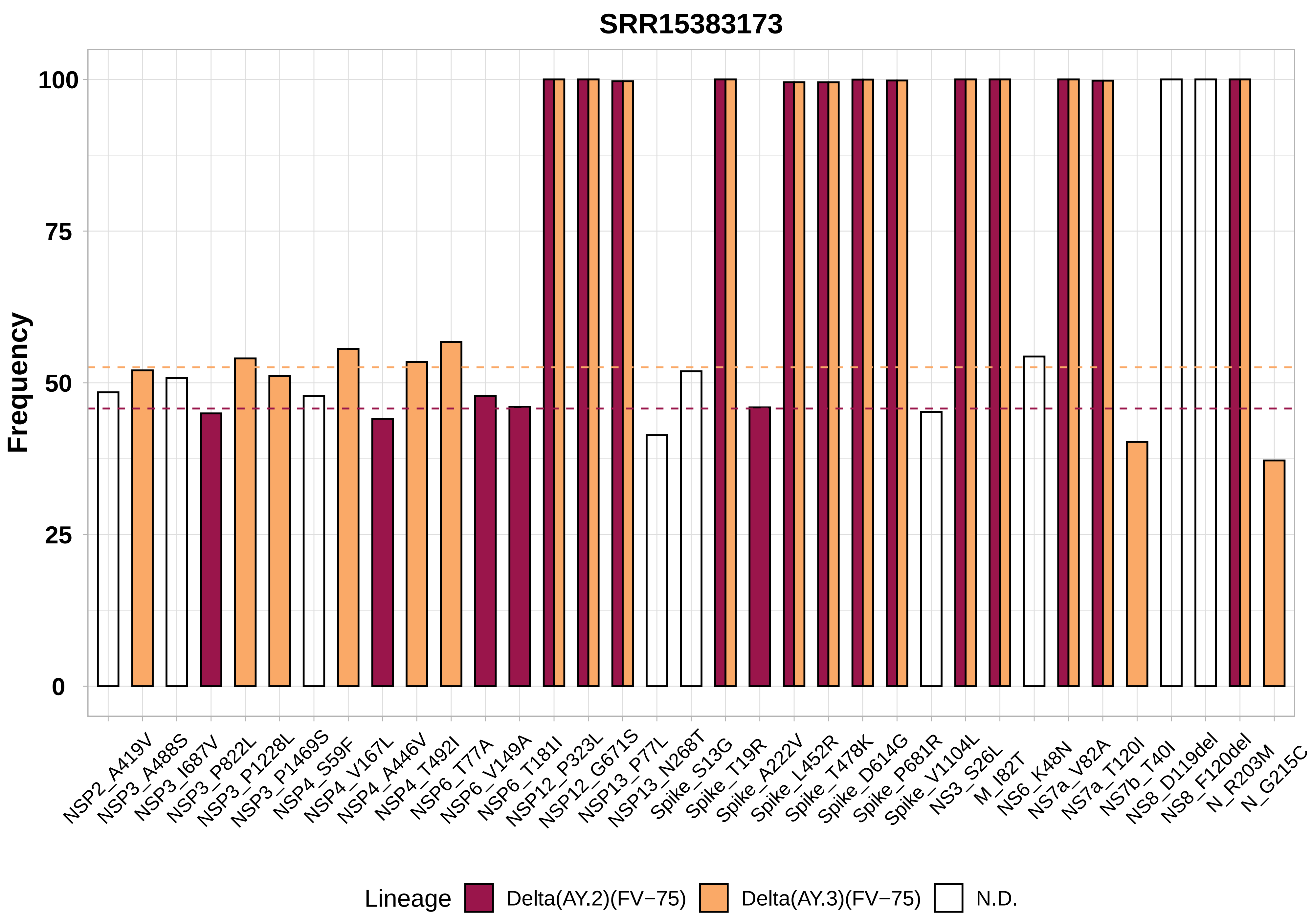

SRR15383173
Frequency
100
75
50
25
0
NSP2_A419V
NSP3_A488S
NSP3_I687V
NSP3_P822L
NSP3_P1228L
NSP3_P1469S
NSP4_S59F
NSP4_V167L
NSP4_A446V
NSP4_T492I
NSP6_T77A
NSP6_V149A
NSP6_T181I
NSP12_P323L
NSP12_G671S
NSP13_P77L
NSP13_N268T
Spike_S13G
Spike_T19R
Spike_A222V
Spike_L452R
Spike_T478K
Spike_D614G
Spike_P681R
Spike_V1104L
NS3_S26L
M_I82T
NS6_K48N
NS7a_V82A
NS7a_T120I
NS7b_T40I
NS8_D119del
NS8_F120del
N_R203M
N_G215C
Lineage
Delta(AY.2)(FV−75)
Delta(AY.3)(FV−75)
N.D.

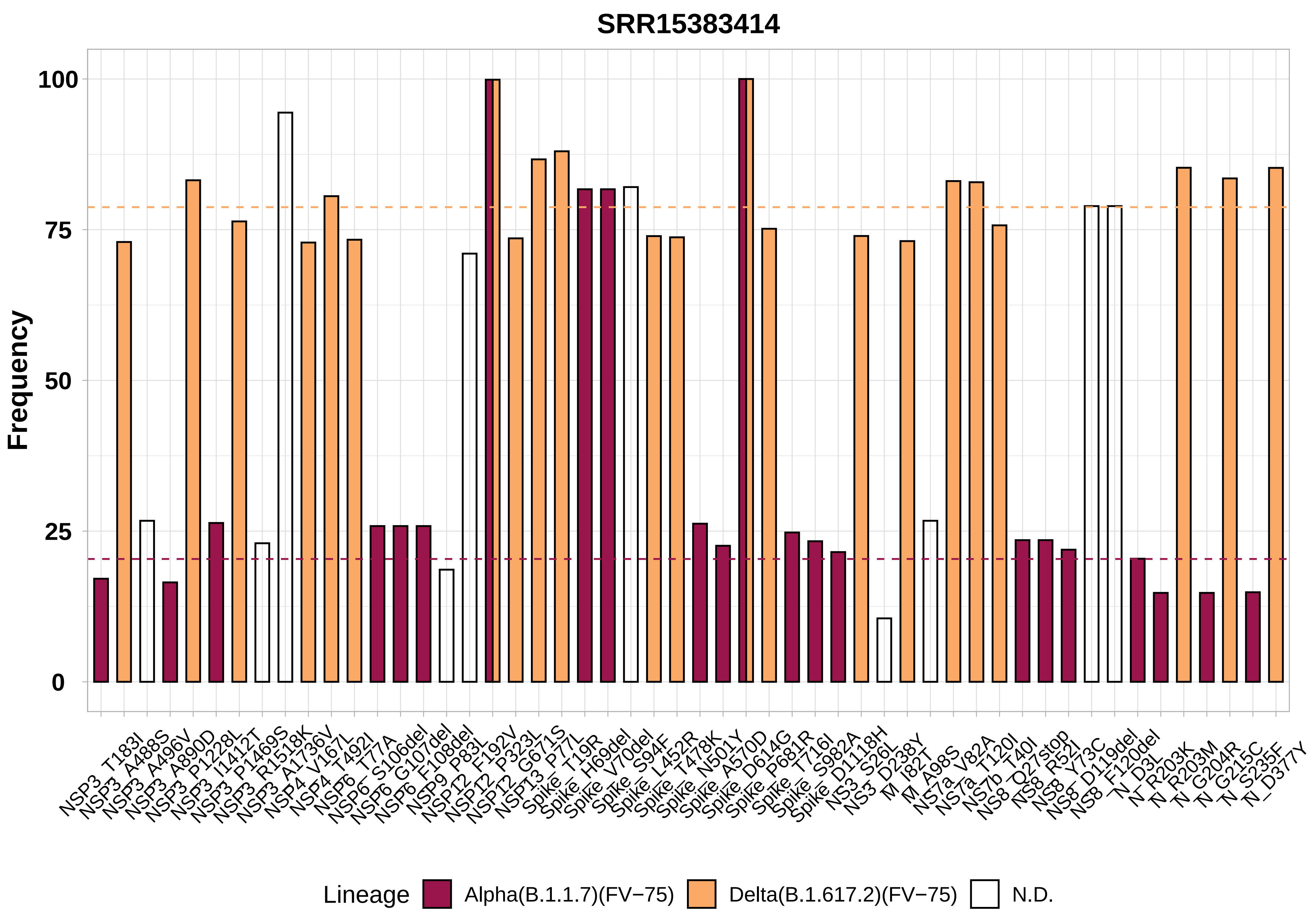
SRR15383414
Frequency
100
75
50
25
0
NSP3_T183I
NSP3_A488S
NSP3_A496V
NSP3_A890D
NSP3_P1228L
NSP3_I1412T
NSP3_P1469S
NSP3_R1518K
NSP3_A1736V
NSP4_V167L
NSP4_T492I
NSP6_T77A
NSP6_S106del
NSP6_G107del
NSP6_F108del
NSP9_P83L
NSP12_F192V
NSP12_P323L
NSP12_G671S
NSP13_P77L
Spike_T19R
Spike_H69del
Spike_V70del
Spike_S94F
Spike_L452R
Spike_T478K
Spike_N501Y
Spike_A570D
Spike_D614G
Spike_P681R
Spike_T716I
Spike_S982A
Spike_D1118H
NS3_S26L
NS3_D238Y
M_I82T
M_A98S
NS7a_V82A
NS7a_T120I
NS7b_T40I
NS8_Q27stop
NS8_R52I
NS8_Y73C
NS8_D119del
NS8_F120del
N_D3L
N_R203K
N_R203M
N_G204R
N_G215C
N_S235F
N_D377Y
Lineage
Alpha(B.1.1.7)(FV−75)
Delta(B.1.617.2)(FV−75)
N.D.

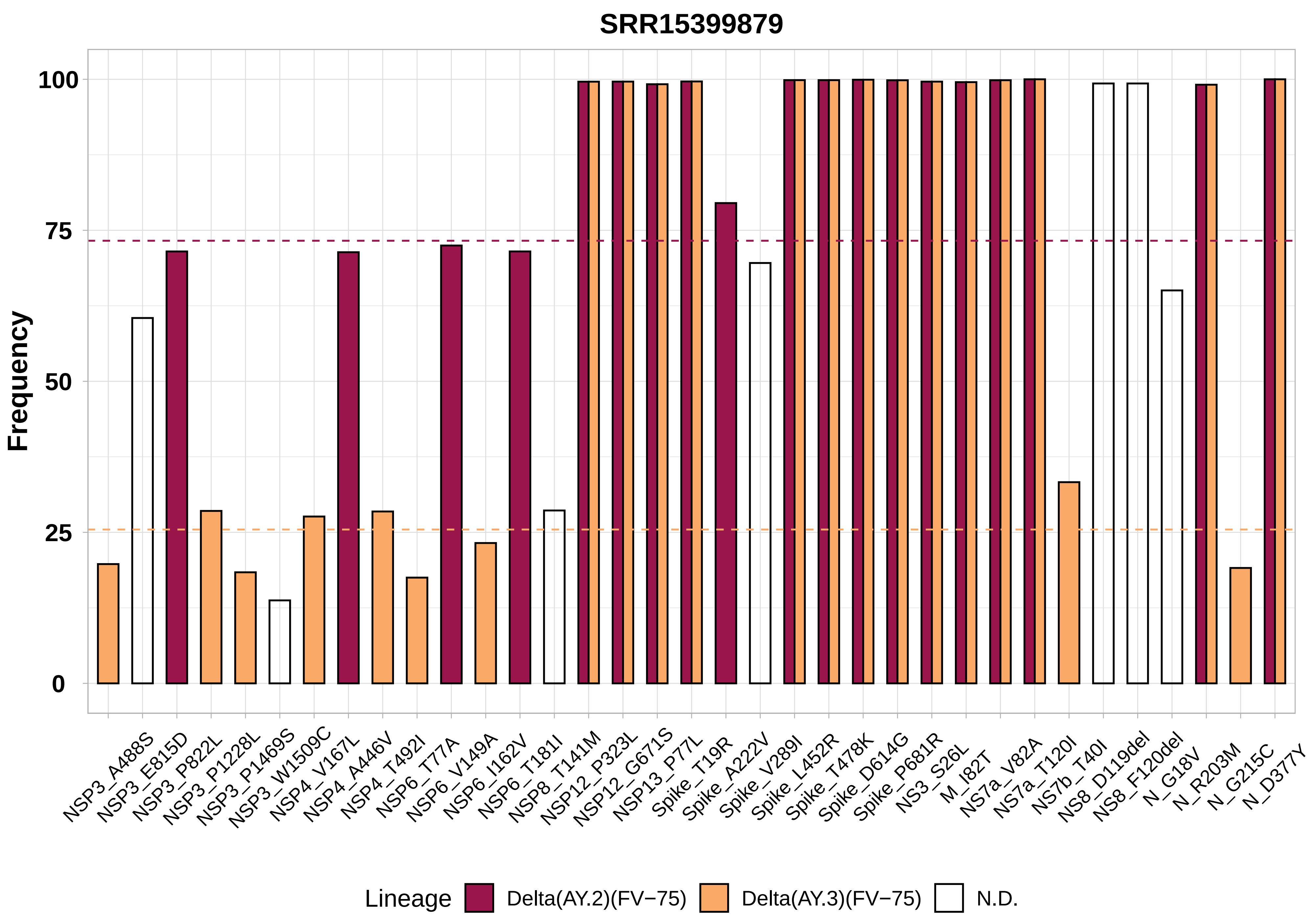

SRR15399879
Frequency
100
75
50
25
0
NSP3_A488S
NSP3_E815D
NSP3_P822L
NSP3_P1228L
NSP3_P1469S
NSP3_W1509C
NSP4_V167L
NSP4_A446V
NSP4_T492I
NSP6_T77A
NSP6_V149A
NSP6_I162V
NSP6_T181I
NSP8_T141M
NSP12_P323L
NSP12_G671S
NSP13_P77L
Spike_T19R
Spike_A222V
Spike_V289I
Spike_L452R
Spike_T478K
Spike_D614G
Spike_P681R
NS3_S26L
M_I82T
NS7a_V82A
NS7a_T120I
NS7b_T40I
NS8_D119del
NS8_F120del
N_G18V
N_R203M
N_G215C
N_D377Y
Lineage
Delta(AY.2)(FV−75)
Delta(AY.3)(FV−75)
N.D.

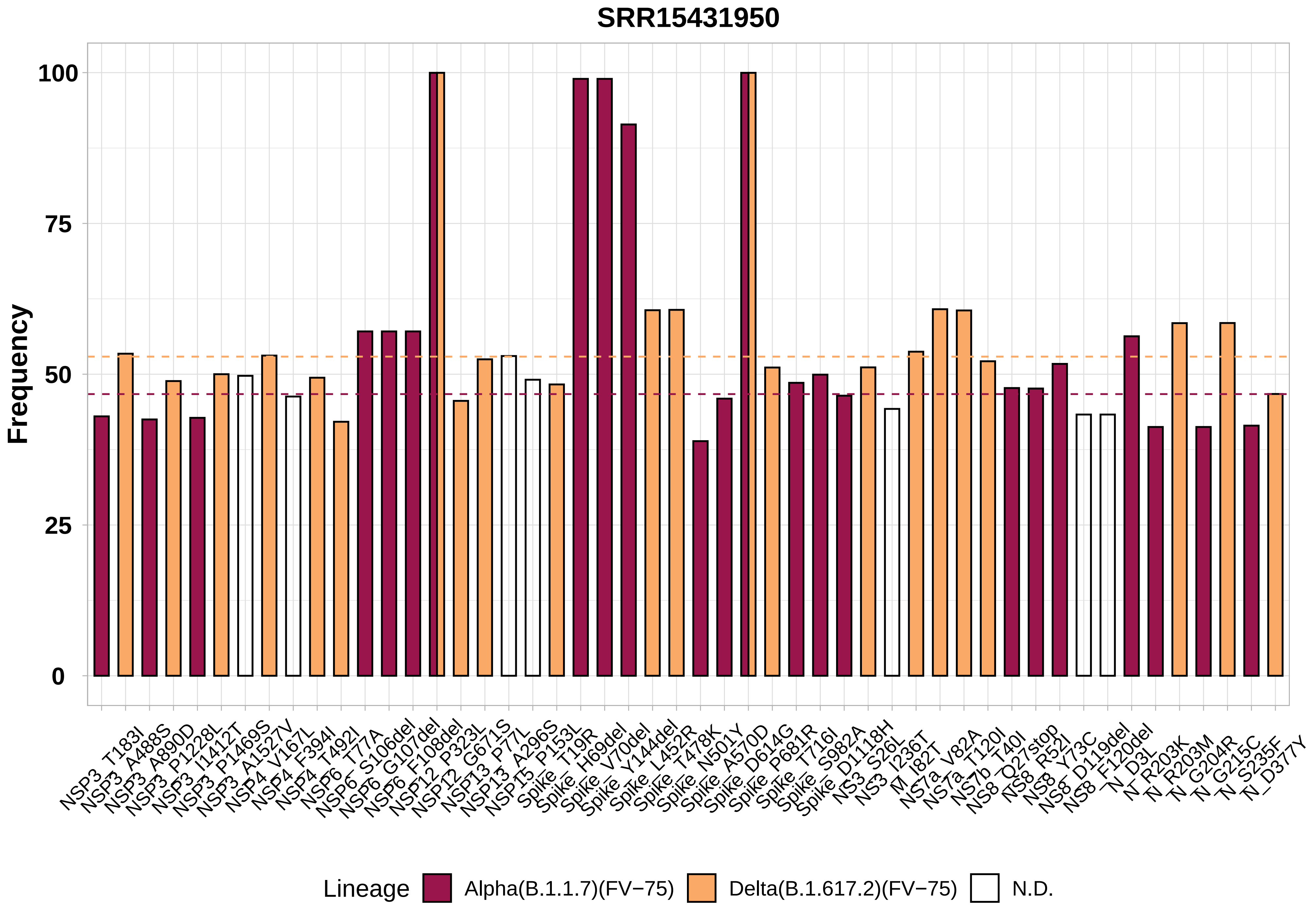

SRR15431950
Frequency
100
75
50
25
0
NSP3_T183I
NSP3_A488S
NSP3_A890D
NSP3_P1228L
NSP3_I1412T
NSP3_P1469S
NSP3_A1527V
NSP4_V167L
NSP4_F394I
NSP4_T492I
NSP6_T77A
NSP6_S106del
NSP6_G107del
NSP6_F108del
NSP12_P323L
NSP12_G671S
NSP13_P77L
NSP13_A296S
NSP15_P153L
Spike_T19R
Spike_H69del
Spike_V70del
Spike_Y144del
Spike_L452R
Spike_T478K
Spike_N501Y
Spike_A570D
Spike_D614G
Spike_P681R
Spike_T716I
Spike_S982A
Spike_D1118H
NS3_S26L
NS3_I236T
M_I82T
NS7a_V82A
NS7a_T120I
NS7b_T40I
NS8_Q27stop
NS8_R52I
NS8_Y73C
NS8_D119del
NS8_F120del
N_D3L
N_R203K
N_R203M
N_G204R
N_G215C
N_S235F
N_D377Y
Lineage
Alpha(B.1.1.7)(FV−75)
Delta(B.1.617.2)(FV−75)
N.D.

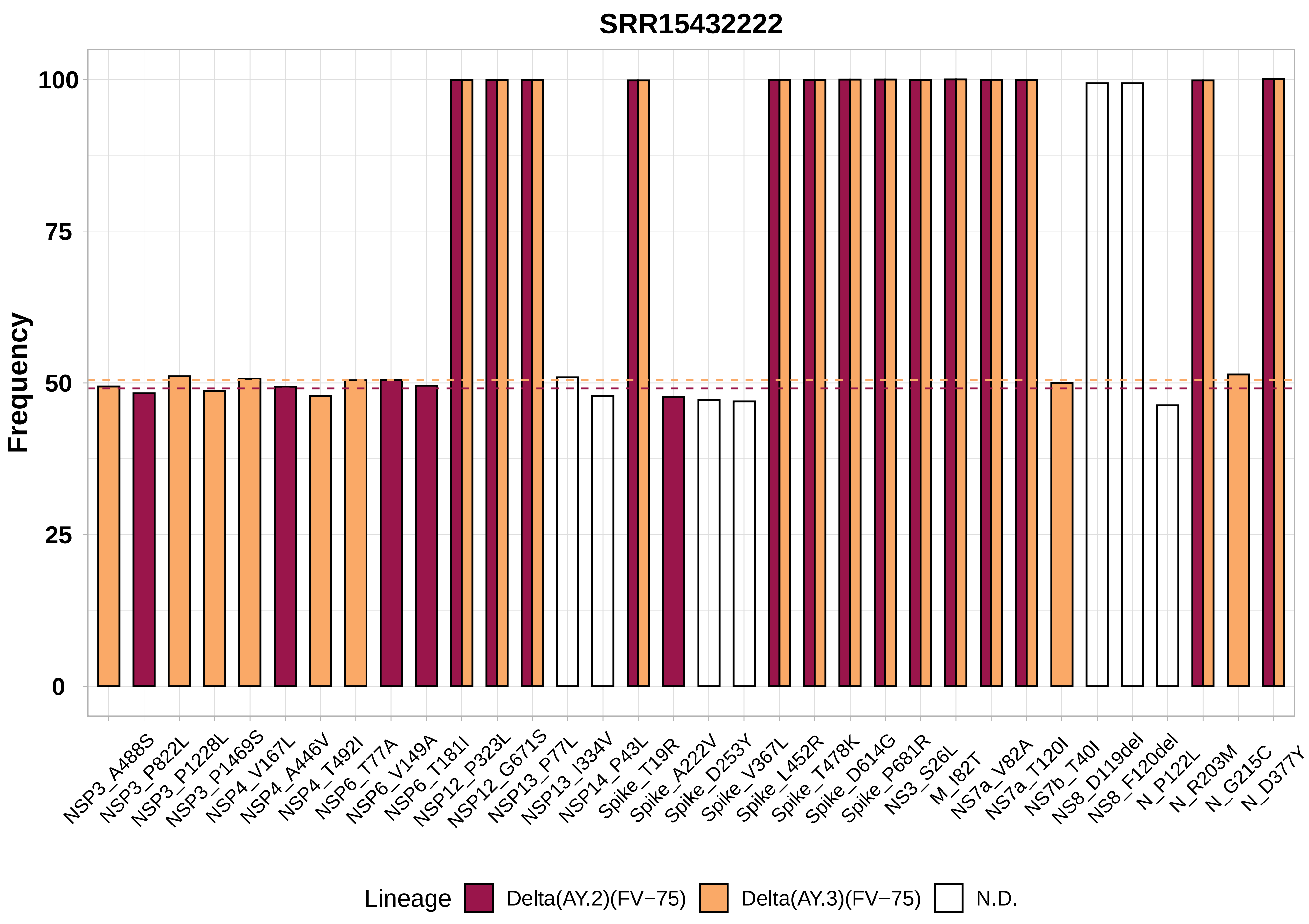

SRR15432222
Frequency
100
75
50
25
0
NSP3_A488S
NSP3_P822L
NSP3_P1228L
NSP3_P1469S
NSP4_V167L
NSP4_A446V
NSP4_T492I
NSP6_T77A
NSP6_V149A
NSP6_T181I
NSP12_P323L
NSP12_G671S
NSP13_P77L
NSP13_I334V
NSP14_P43L
Spike_T19R
Spike_A222V
Spike_D253Y
Spike_V367L
Spike_L452R
Spike_T478K
Spike_D614G
Spike_P681R
NS3_S26L
M_I82T
NS7a_V82A
NS7a_T120I
NS7b_T40I
NS8_D119del
NS8_F120del
N_P122L
N_R203M
N_G215C
N_D377Y
Lineage
Delta(AY.2)(FV−75)
Delta(AY.3)(FV−75)
N.D.

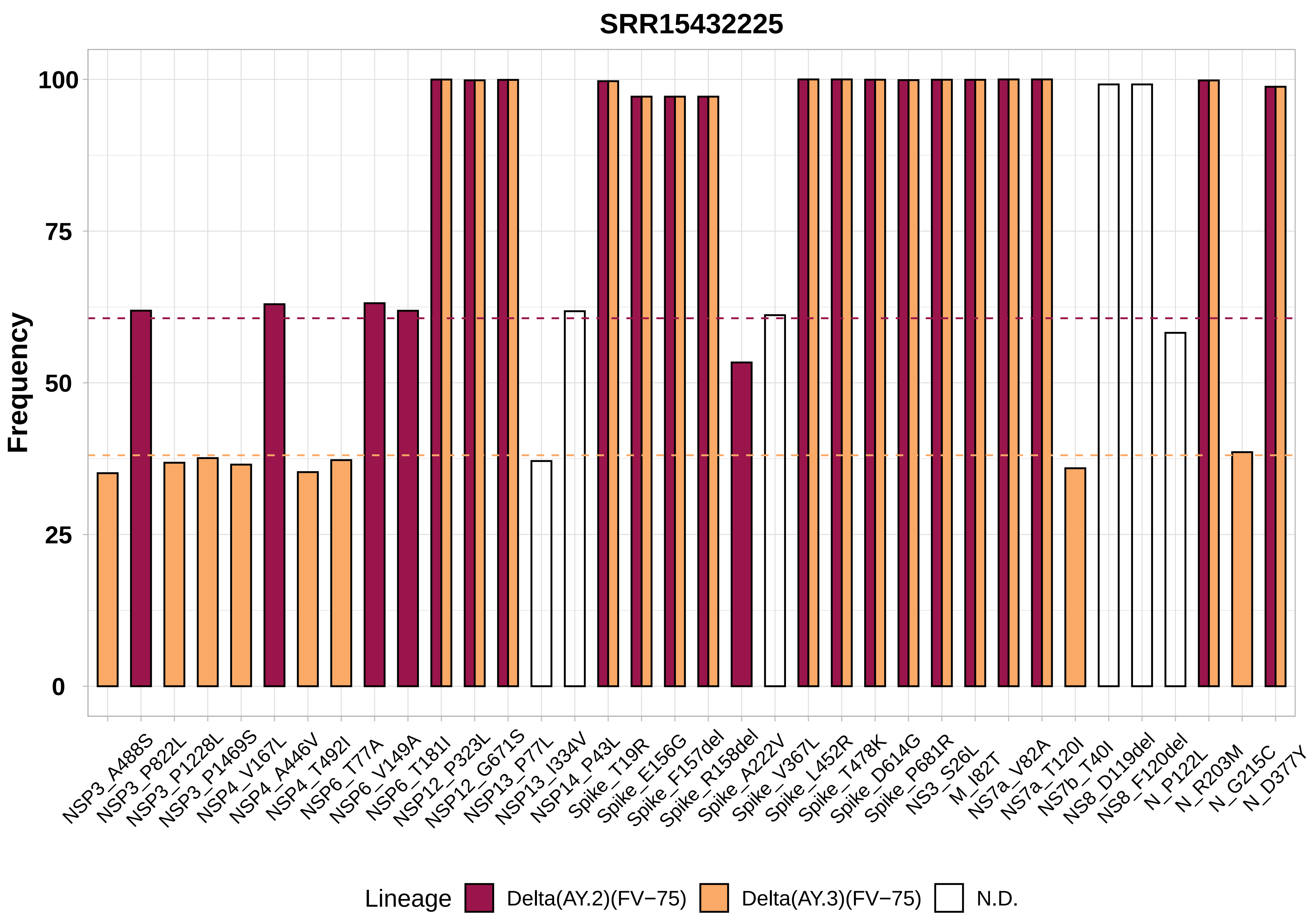

SRR15432225
Frequency
100
75
50
25
0
NSP3_A488S
NSP3_P822L
NSP3_P1228L
NSP3_P1469S
NSP4_V167L
NSP4_A446V
NSP4_T492I
NSP6_T77A
NSP6_V149A
NSP6_T181I
NSP12_P323L
NSP12_G671S
NSP13_P77L
NSP13_I334V
NSP14_P43L
Spike_T19R
Spike_E156G
Spike_F157del
Spike_R158del
Spike_A222V
Spike_V367L
Spike_L452R
Spike_T478K
Spike_D614G
Spike_P681R
NS3_S26L
M_I82T
NS7a_V82A
NS7a_T120I
NS7b_T40I
NS8_D119del
NS8_F120del
N_P122L
N_R203M
N_G215C
N_D377Y
Lineage
Delta(AY.2)(FV−75)
Delta(AY.3)(FV−75)
N.D.

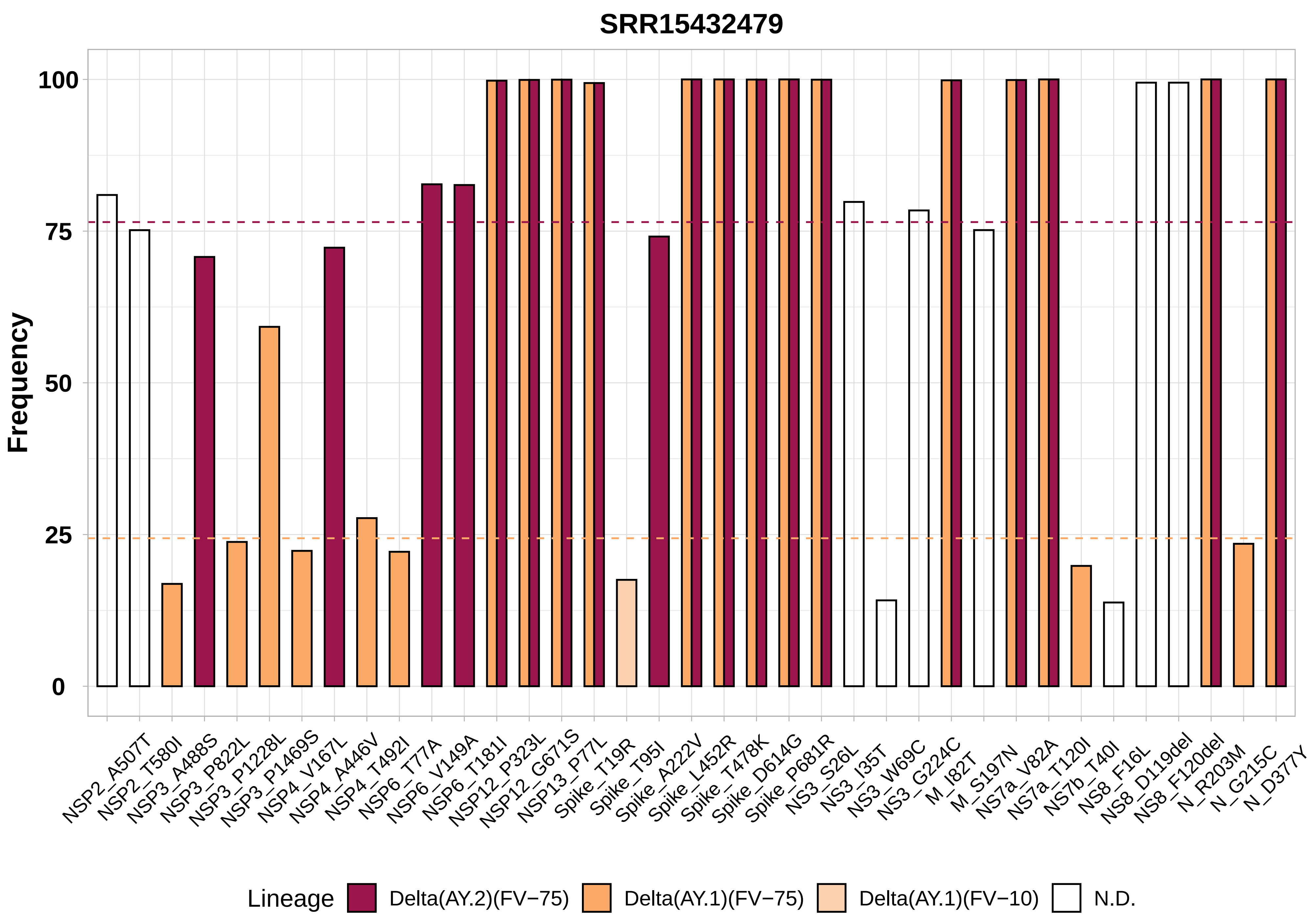
SRR15432479
Frequency
100
75
50
25
0
NSP2_A507T
NSP2_T580I
NSP3_A488S
NSP3_P822L
NSP3_P1228L
NSP3_P1469S
NSP4_V167L
NSP4_A446V
NSP4_T492I
NSP6_T77A
NSP6_V149A
NSP6_T181I
NSP12_P323L
NSP12_G671S
NSP13_P77L
Spike_T19R
Spike_T95I
Spike_A222V
Spike_L452R
Spike_T478K
Spike_D614G
Spike_P681R
NS3_S26L
NS3_I35T
NS3_W69C
NS3_G224C
M_I82T
M_S197N
NS7a_V82A
NS7a_T120I
NS7b_T40I
NS8_F16L
NS8_D119del
NS8_F120del
N_R203M
N_G215C
N_D377Y
Lineage
Delta(AY.2)(FV−75)
Delta(AY.1)(FV−75)
Delta(AY.1)(FV−10)
N.D.

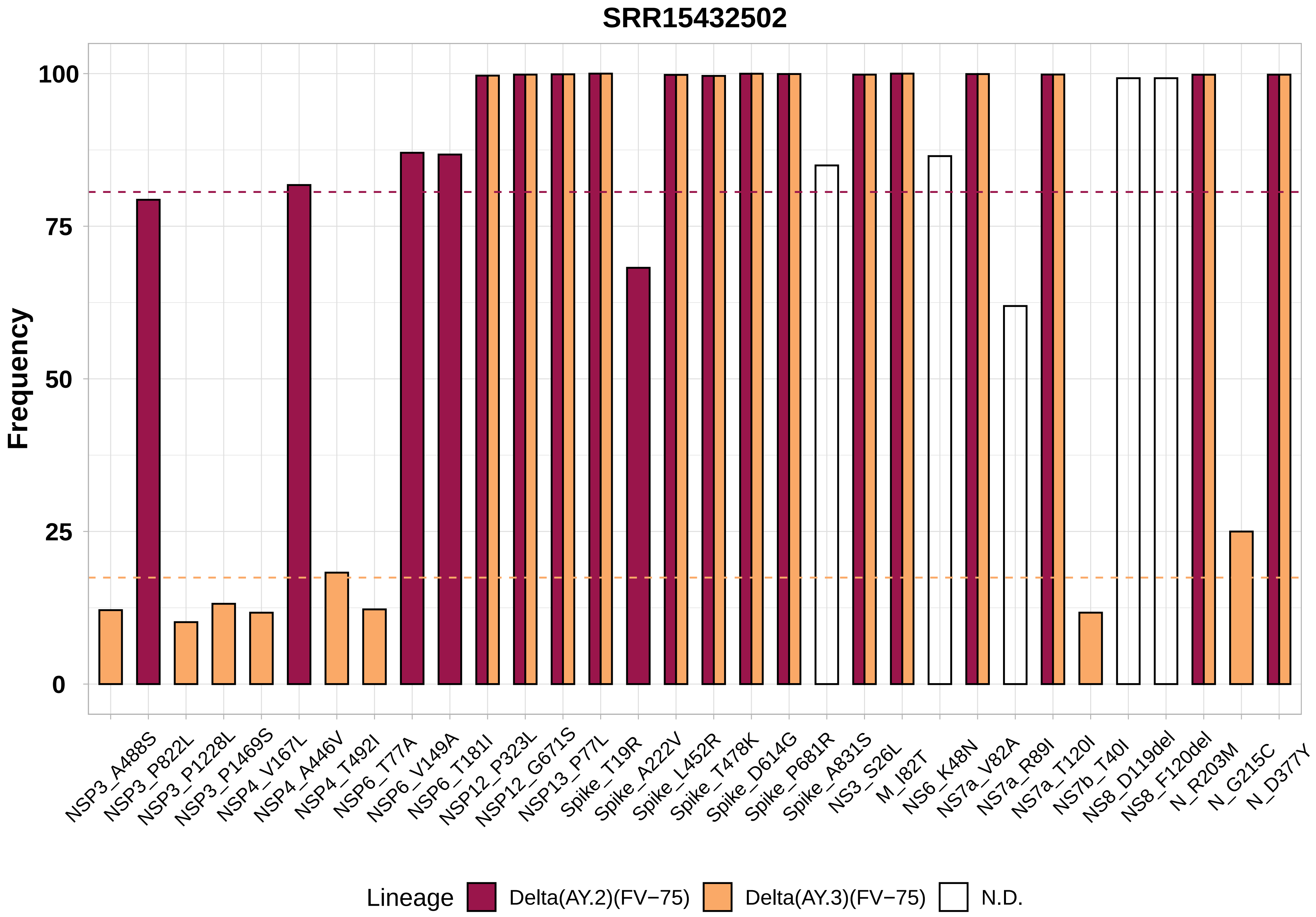
SRR15432502
Frequency
100
75
50
25
0
NSP3_A488S
NSP3_P822L
NSP3_P1228L
NSP3_P1469S
NSP4_V167L
NSP4_A446V
NSP4_T492I
NSP6_T77A
NSP6_V149A
NSP6_T181I
NSP12_P323L
NSP12_G671S
NSP13_P77L
Spike_T19R
Spike_A222V
Spike_L452R
Spike_T478K
Spike_D614G
Spike_P681R
Spike_A831S
NS3_S26L
M_I82T
NS6_K48N
NS7a_V82A
NS7a_R89I
NS7a_T120I
NS7b_T40I
NS8_D119del
NS8_F120del
N_R203M
N_G215C
N_D377Y
Lineage
Delta(AY.2)(FV−75)
Delta(AY.3)(FV−75)
N.D.

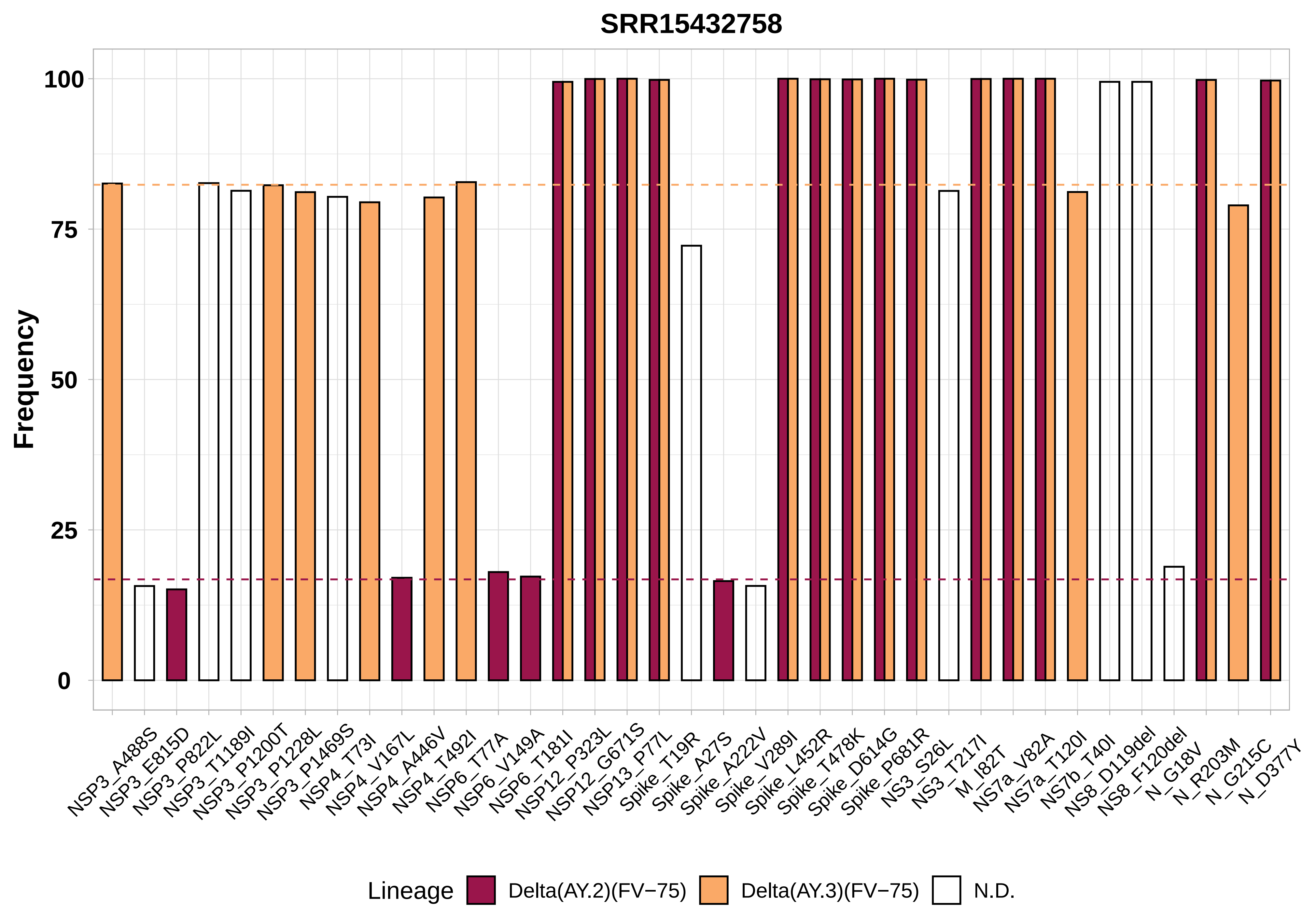
SRR15432758
Frequency
100
75
50
25
0
NSP3_A488S
NSP3_E815D
NSP3_P822L
NSP3_T1189I
NSP3_P1200T
NSP3_P1228L
NSP3_P1469S
NSP4_T73I
NSP4_V167L
NSP4_A446V
NSP4_T492I
NSP6_T77A
NSP6_V149A
NSP6_T181I
NSP12_P323L
NSP12_G671S
NSP13_P77L
Spike_T19R
Spike_A27S
Spike_A222V
Spike_V289I
Spike_L452R
Spike_T478K
Spike_D614G
Spike_P681R
NS3_S26L
NS3_T217I
M_I82T
NS7a_V82A
NS7a_T120I
NS7b_T40I
NS8_D119del
NS8_F120del
N_G18V
N_R203M
N_G215C
N_D377Y
Lineage
Delta(AY.2)(FV−75)
Delta(AY.3)(FV−75)
N.D.

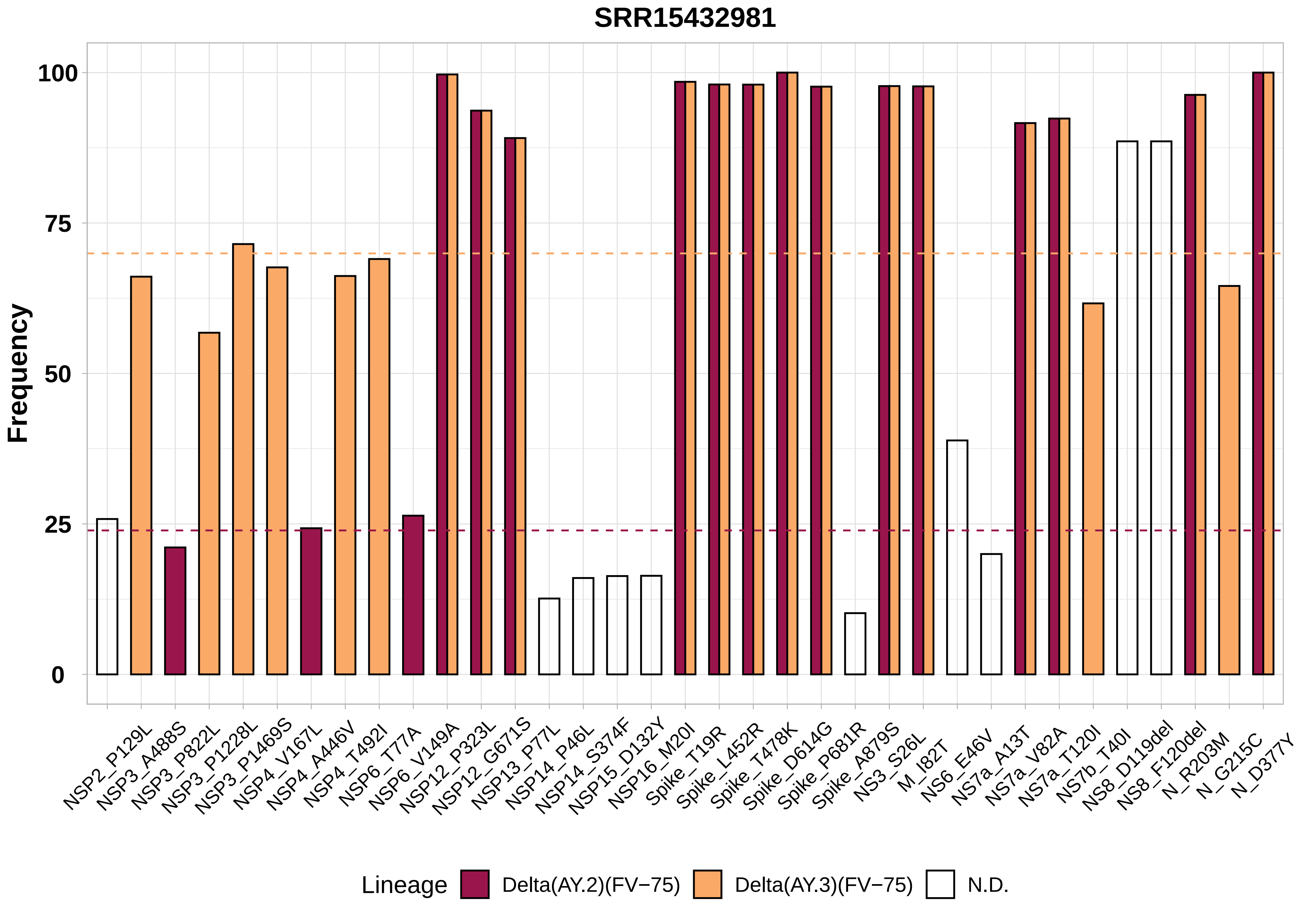

SRR15432981
Frequency
100
75
50
25
0
NSP2_P129L
NSP3_A488S
NSP3_P822L
NSP3_P1228L
NSP3_P1469S
NSP4_V167L
NSP4_A446V
NSP4_T492I
NSP6_T77A
NSP6_V149A
NSP12_P323L
NSP12_G671S
NSP13_P77L
NSP14_P46L
NSP14_S374F
NSP15_D132Y
NSP16_M20I
Spike_T19R
Spike_L452R
Spike_T478K
Spike_D614G
Spike_P681R
Spike_A879S
NS3_S26L
M_I82T
NS6_E46V
NS7a_A13T
NS7a_V82A
NS7a_T120I
NS7b_T40I
NS8_D119del
NS8_F120del
N_R203M
N_G215C
N_D377Y
Lineage
Delta(AY.2)(FV−75)
Delta(AY.3)(FV−75)
N.D.

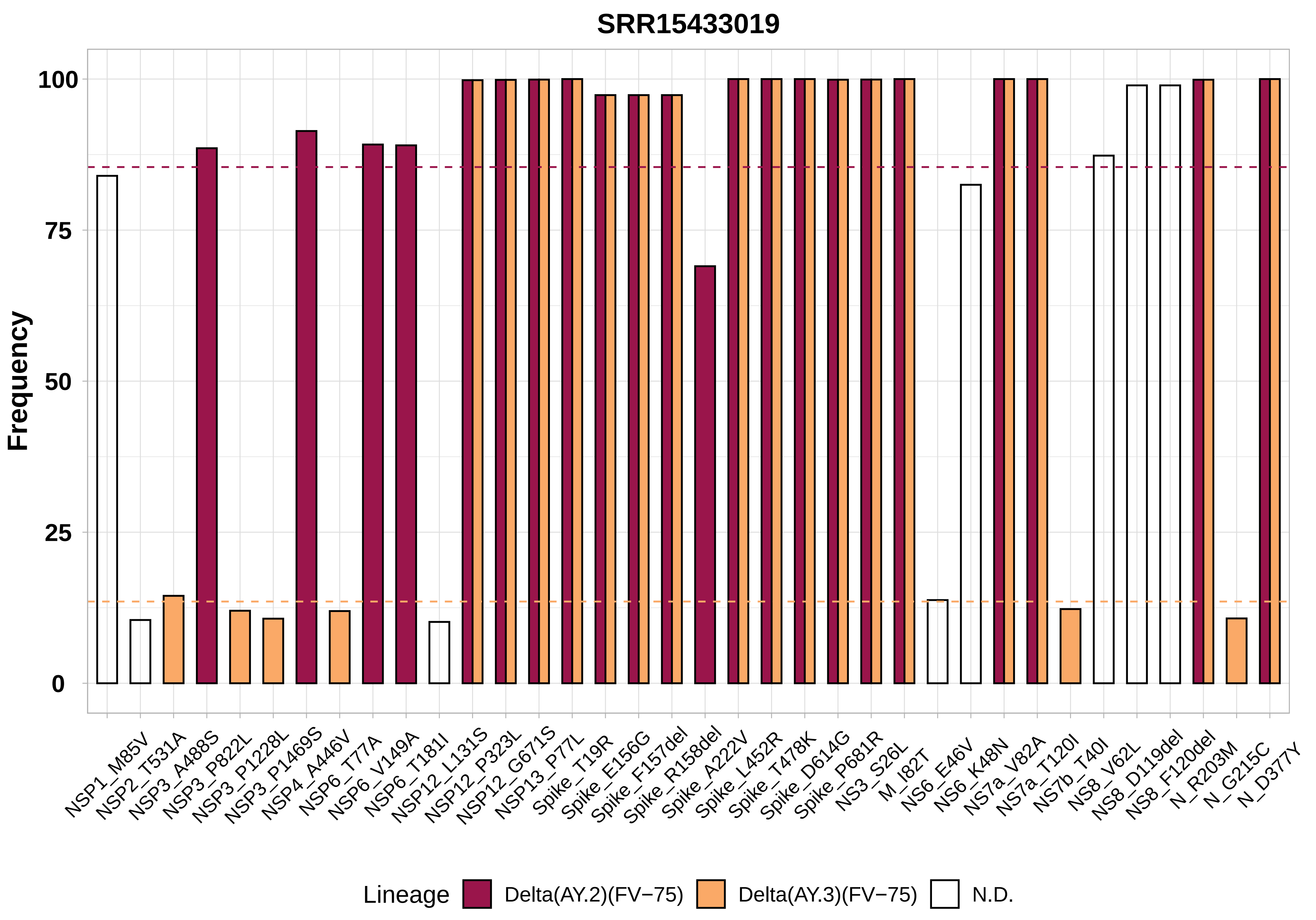

SRR15433019
Frequency
100
75
50
25
0
NSP1_M85V
NSP2_T531A
NSP3_A488S
NSP3_P822L
NSP3_P1228L
NSP3_P1469S
NSP4_A446V
NSP6_T77A
NSP6_V149A
NSP6_T181I
NSP12_L131S
NSP12_P323L
NSP12_G671S
NSP13_P77L
Spike_T19R
Spike_E156G
Spike_F157del
Spike_R158del
Spike_A222V
Spike_L452R
Spike_T478K
Spike_D614G
Spike_P681R
NS3_S26L
M_I82T
NS6_E46V
NS6_K48N
NS7a_V82A
NS7a_T120I
NS7b_T40I
NS8_V62L
NS8_D119del
NS8_F120del
N_R203M
N_G215C
N_D377Y
Lineage
Delta(AY.2)(FV−75)
Delta(AY.3)(FV−75)
N.D.

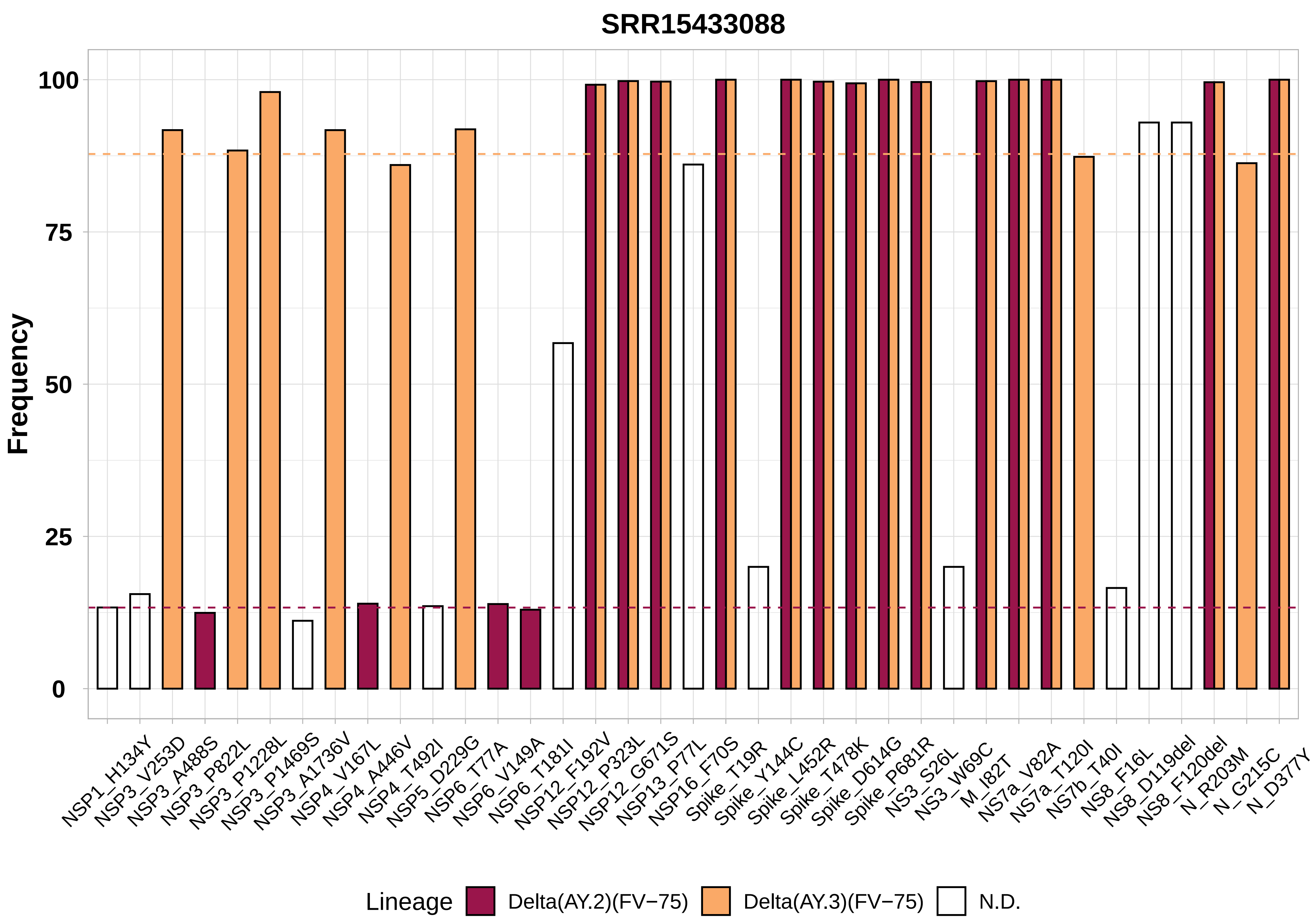

SRR15433088
Frequency
100
75
50
25
0
NSP1_H134Y
NSP3_V253D
NSP3_A488S
NSP3_P822L
NSP3_P1228L
NSP3_P1469S
NSP3_A1736V
NSP4_V167L
NSP4_A446V
NSP4_T492I
NSP5_D229G
NSP6_T77A
NSP6_V149A
NSP6_T181I
NSP12_F192V
NSP12_P323L
NSP12_G671S
NSP13_P77L
NSP16_F70S
Spike_T19R
Spike_Y144C
Spike_L452R
Spike_T478K
Spike_D614G
Spike_P681R
NS3_S26L
NS3_W69C
M_I82T
NS7a_V82A
NS7a_T120I
NS7b_T40I
NS8_F16L
NS8_D119del
NS8_F120del
N_R203M
N_G215C
N_D377Y
Lineage
Delta(AY.2)(FV−75)
Delta(AY.3)(FV−75)
N.D.

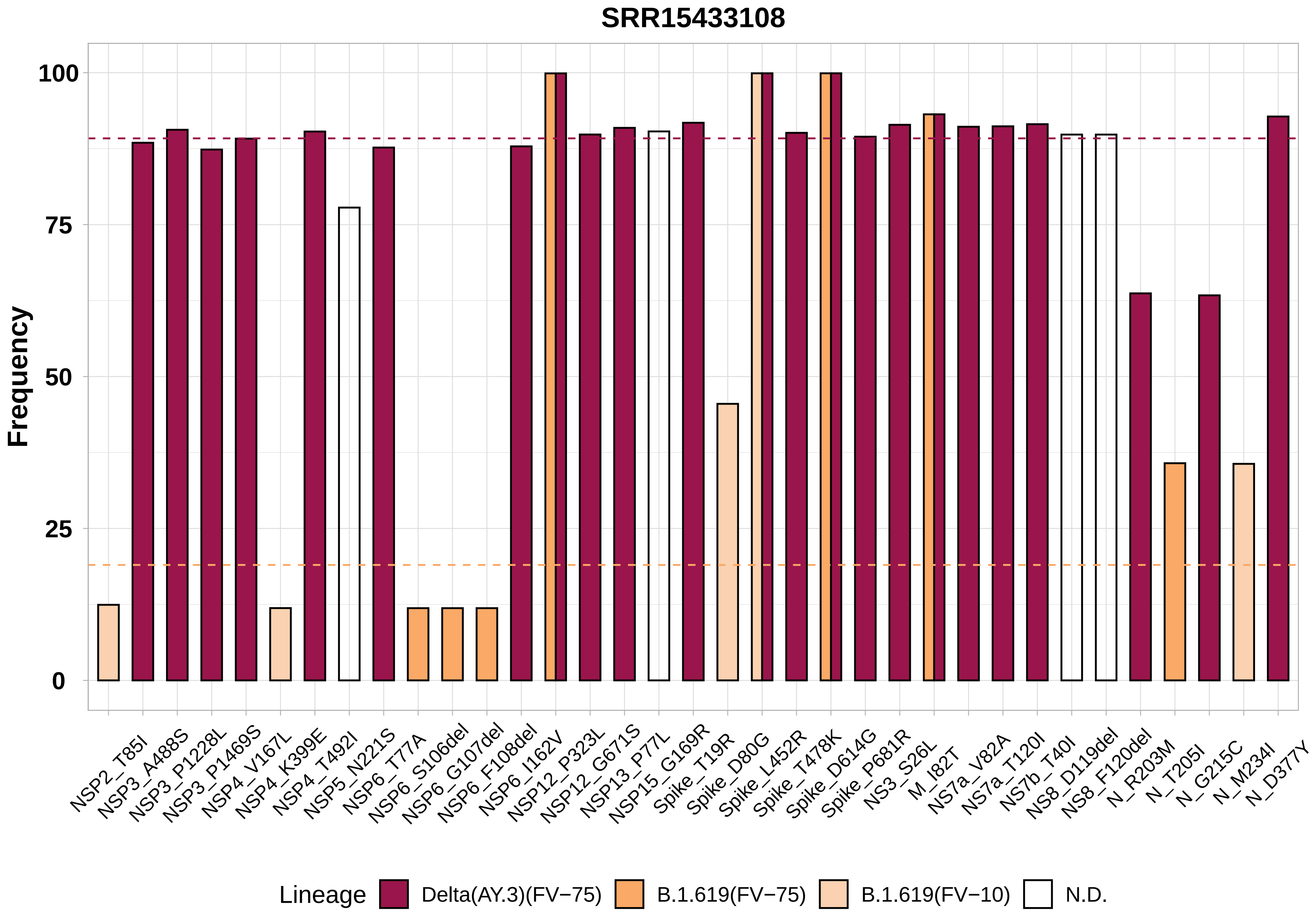
SRR15433108
Frequency
100
75
50
25
0
NSP2_T85I
NSP3_A488S
NSP3_P1228L
NSP3_P1469S
NSP4_V167L
NSP4_K399E
NSP4_T492I
NSP5_N221S
NSP6_T77A
NSP6_S106del
NSP6_G107del
NSP6_F108del
NSP6_I162V
NSP12_P323L
NSP12_G671S
NSP13_P77L
NSP15_G169R
Spike_T19R
Spike_D80G
Spike_L452R
Spike_T478K
Spike_D614G
Spike_P681R
NS3_S26L
M_I82T
NS7a_V82A
NS7a_T120I
NS7b_T40I
NS8_D119del
NS8_F120del
N_R203M
N_T205I
N_G215C
N_M234I
N_D377Y
Lineage
Delta(AY.3)(FV−75)
B.1.619(FV−75)
B.1.619(FV−10)
N.D.

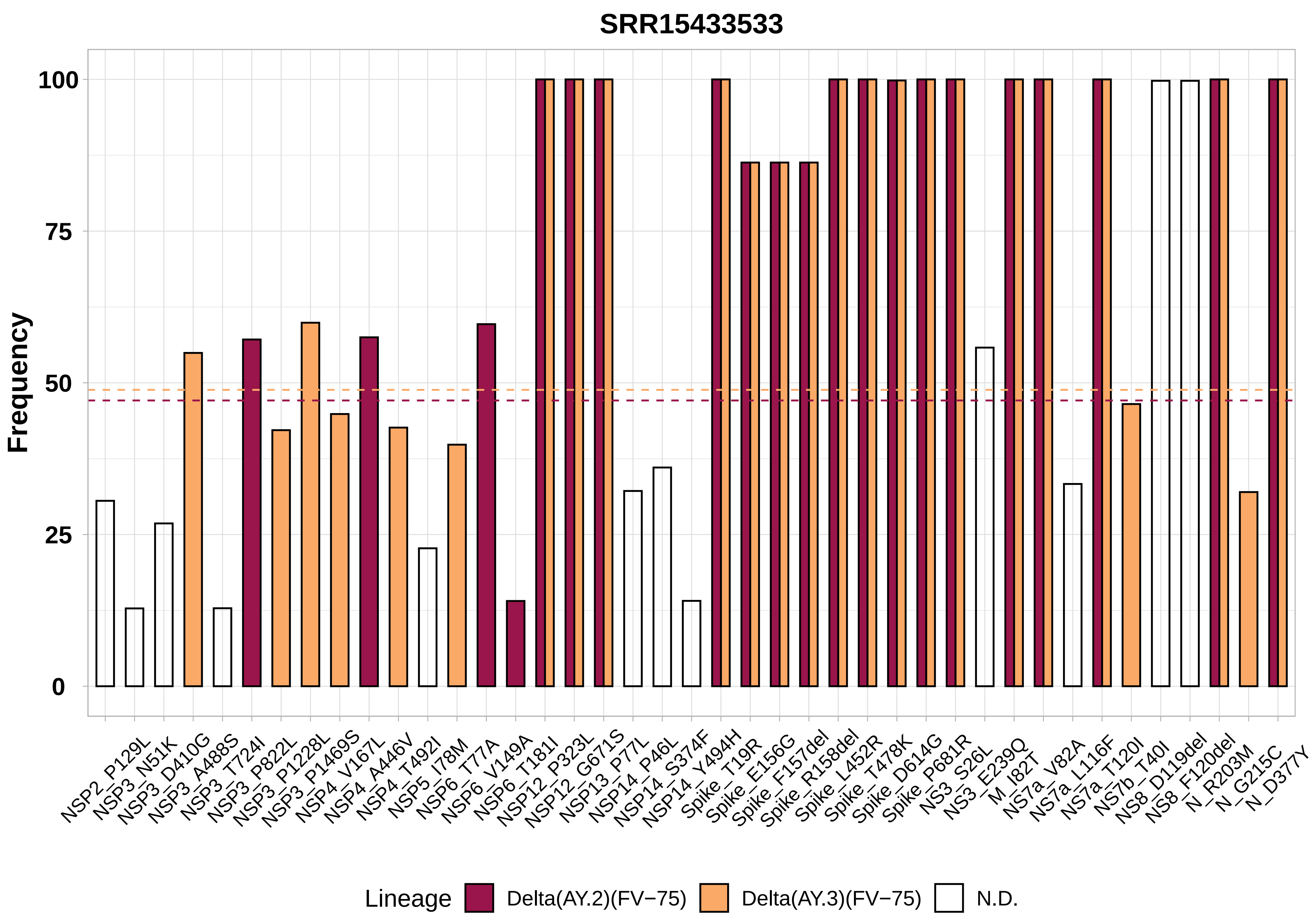

SRR15433533
Frequency
100
75
50
25
0
NSP2_P129L
NSP3_N51K
NSP3_D410G
NSP3_A488S
NSP3_T724I
NSP3_P822L
NSP3_P1228L
NSP3_P1469S
NSP4_V167L
NSP4_A446V
NSP4_T492I
NSP5_I78M
NSP6_T77A
NSP6_V149A
NSP6_T181I
NSP12_P323L
NSP12_G671S
NSP13_P77L
NSP14_P46L
NSP14_S374F
NSP14_Y494H
Spike_T19R
Spike_E156G
Spike_F157del
Spike_R158del
Spike_L452R
Spike_T478K
Spike_D614G
Spike_P681R
NS3_S26L
NS3_E239Q
M_I82T
NS7a_V82A
NS7a_L116F
NS7a_T120I
NS7b_T40I
NS8_D119del
NS8_F120del
N_R203M
N_G215C
N_D377Y
Lineage
Delta(AY.2)(FV−75)
Delta(AY.3)(FV−75)
N.D.

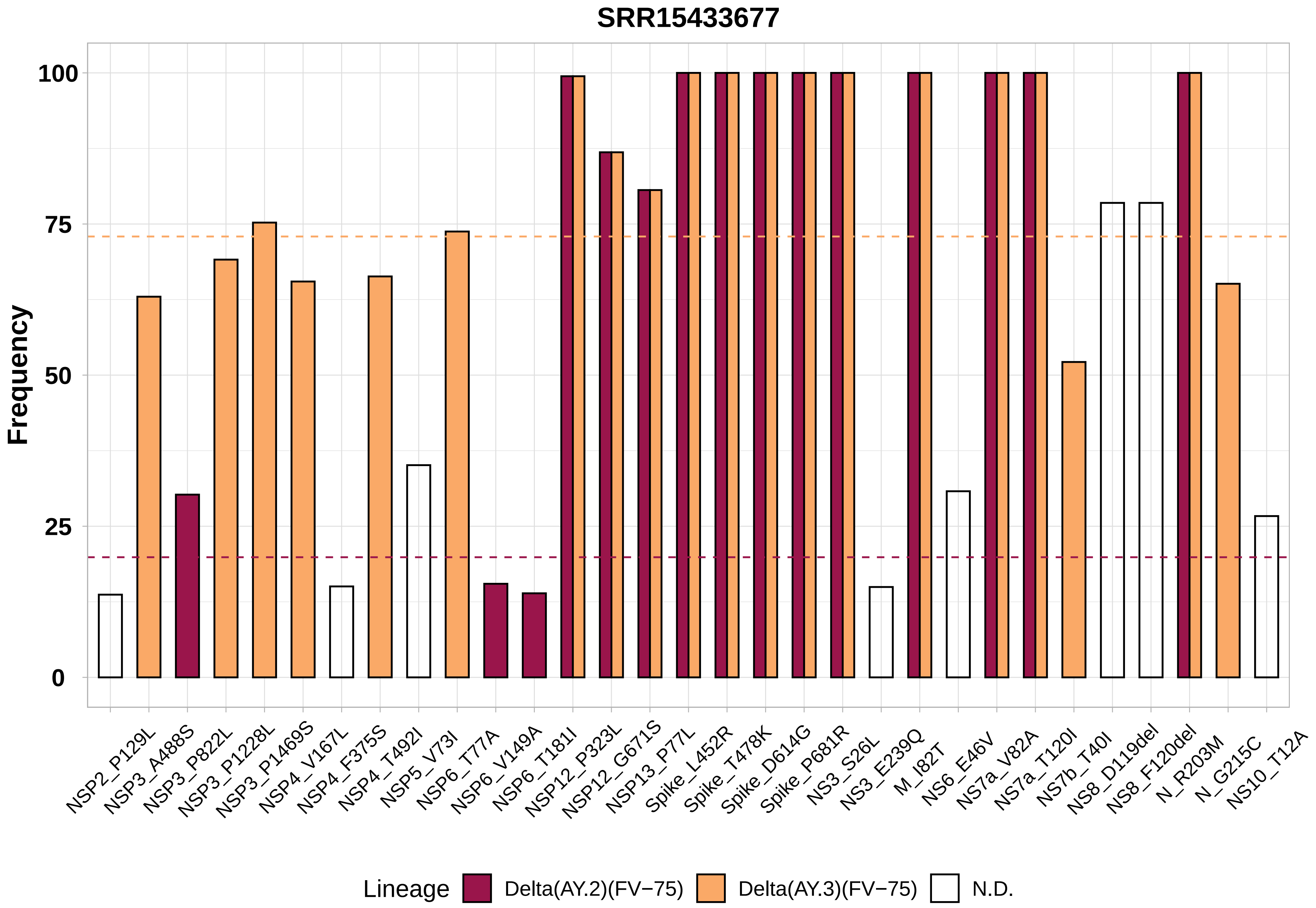

SRR15433677
Frequency
100
75
50
25
0
NSP2_P129L
NSP3_A488S
NSP3_P822L
NSP3_P1228L
NSP3_P1469S
NSP4_V167L
NSP4_F375S
NSP4_T492I
NSP5_V73I
NSP6_T77A
NSP6_V149A
NSP6_T181I
NSP12_P323L
NSP12_G671S
NSP13_P77L
Spike_L452R
Spike_T478K
Spike_D614G
Spike_P681R
NS3_S26L
NS3_E239Q
M_I82T
NS6_E46V
NS7a_V82A
NS7a_T120I
NS7b_T40I
NS8_D119del
NS8_F120del
N_R203M
N_G215C
NS10_T12A
Lineage
Delta(AY.2)(FV−75)
Delta(AY.3)(FV−75)
N.D.

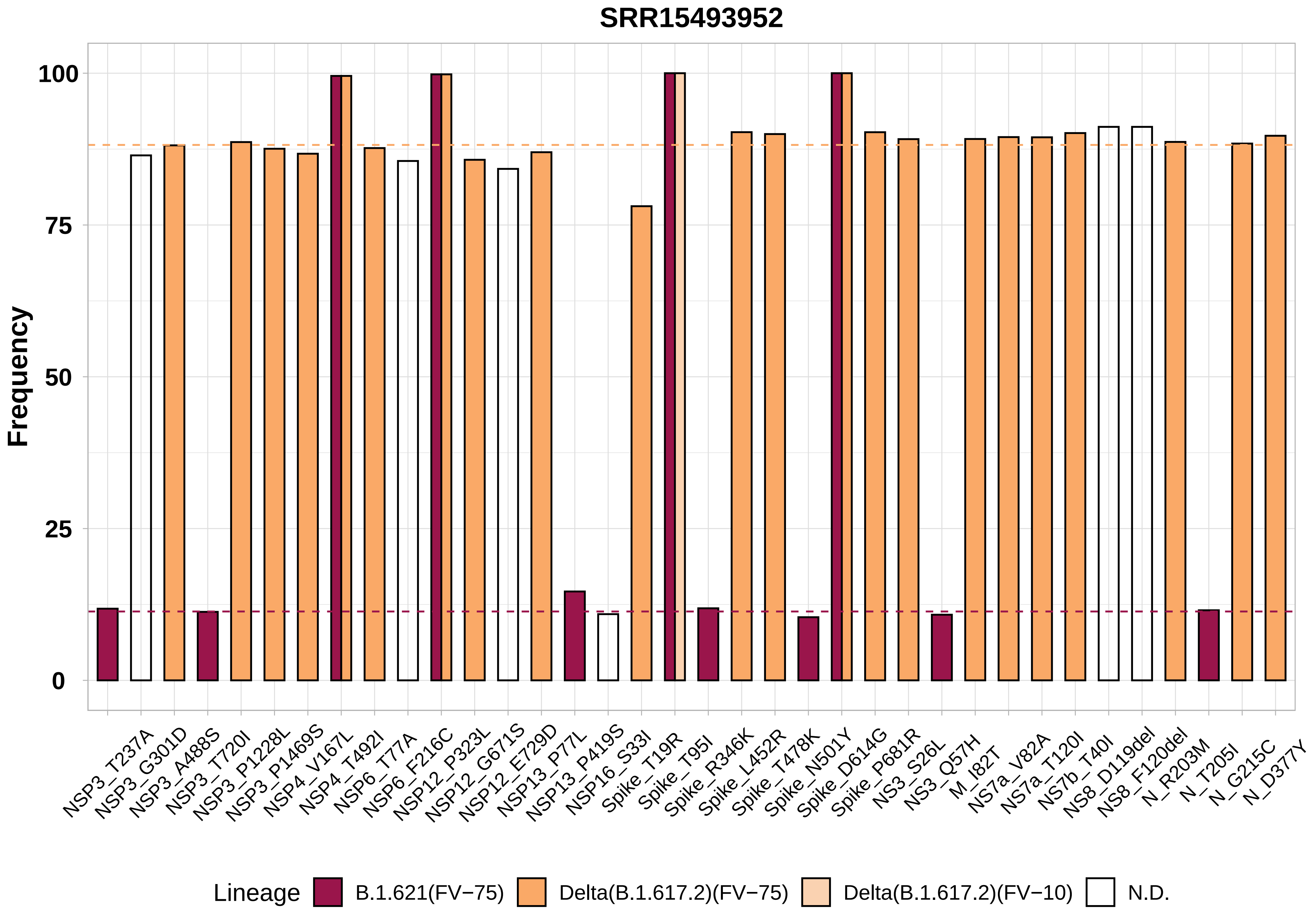
SRR15493952
Frequency
100
75
50
25
0
NSP3_T237A
NSP3_G301D
NSP3_A488S
NSP3_T720I
NSP3_P1228L
NSP3_P1469S
NSP4_V167L
NSP4_T492I
NSP6_T77A
NSP6_F216C
NSP12_P323L
NSP12_G671S
NSP12_E729D
NSP13_P77L
NSP13_P419S
NSP16_S33I
Spike_T19R
Spike_T95I
Spike_R346K
Spike_L452R
Spike_T478K
Spike_N501Y
Spike_D614G
Spike_P681R
NS3_S26L
NS3_Q57H
M_I82T
NS7a_V82A
NS7a_T120I
NS7b_T40I
NS8_D119del
NS8_F120del
N_R203M
N_T205I
N_G215C
N_D377Y
Lineage
B.1.621(FV−75)
Delta(B.1.617.2)(FV−75)
Delta(B.1.617.2)(FV−10)
N.D.

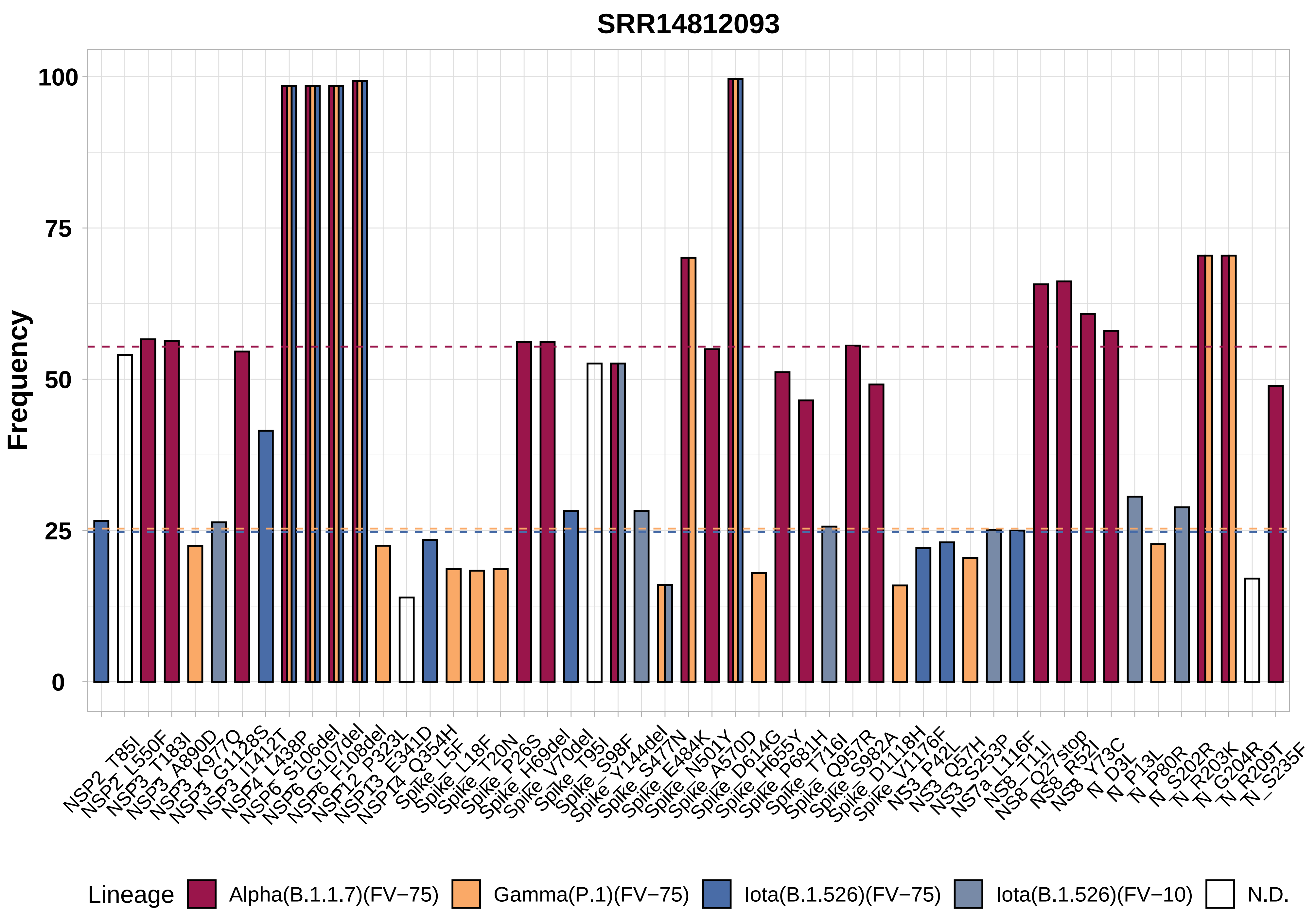

SRR14812093
Frequency
100
75
50
25
0
NSP2_T85I
NSP2_L550F
NSP3_T183I
NSP3_A890D
NSP3_K977Q
NSP3_G1128S
NSP3_I1412T
NSP4_L438P
NSP6_S106del
NSP6_G107del
NSP6_F108del
NSP12_P323L
NSP13_E341D
NSP14_Q354H
Spike_L5F
Spike_L18F
Spike_T20N
Spike_P26S
Spike_H69del
Spike_V70del
Spike_T95I
Spike_S98F
Spike_Y144del
Spike_S477N
Spike_E484K
Spike_N501Y
Spike_A570D
Spike_D614G
Spike_H655Y
Spike_P681H
Spike_T716I
Spike_Q957R
Spike_S982A
Spike_D1118H
Spike_V1176F
NS3_P42L
NS3_Q57H
NS3_S253P
NS7a_L116F
NS8_T11I
NS8_Q27stop
NS8_R52I
NS8_Y73C
N_D3L
N_P13L
N_P80R
N_S202R
N_R203K
N_G204R
N_R209T
N_S235F
Lineage
Alpha(B.1.1.7)(FV−75)
Gamma(P.1)(FV−75)
Iota(B.1.526)(FV−75)
Iota(B.1.526)(FV−10)
N.D.
