## Supplemental Figure 6 for "Genomic evidence for divergent co-infections of SARS-CoV-2 lineages"

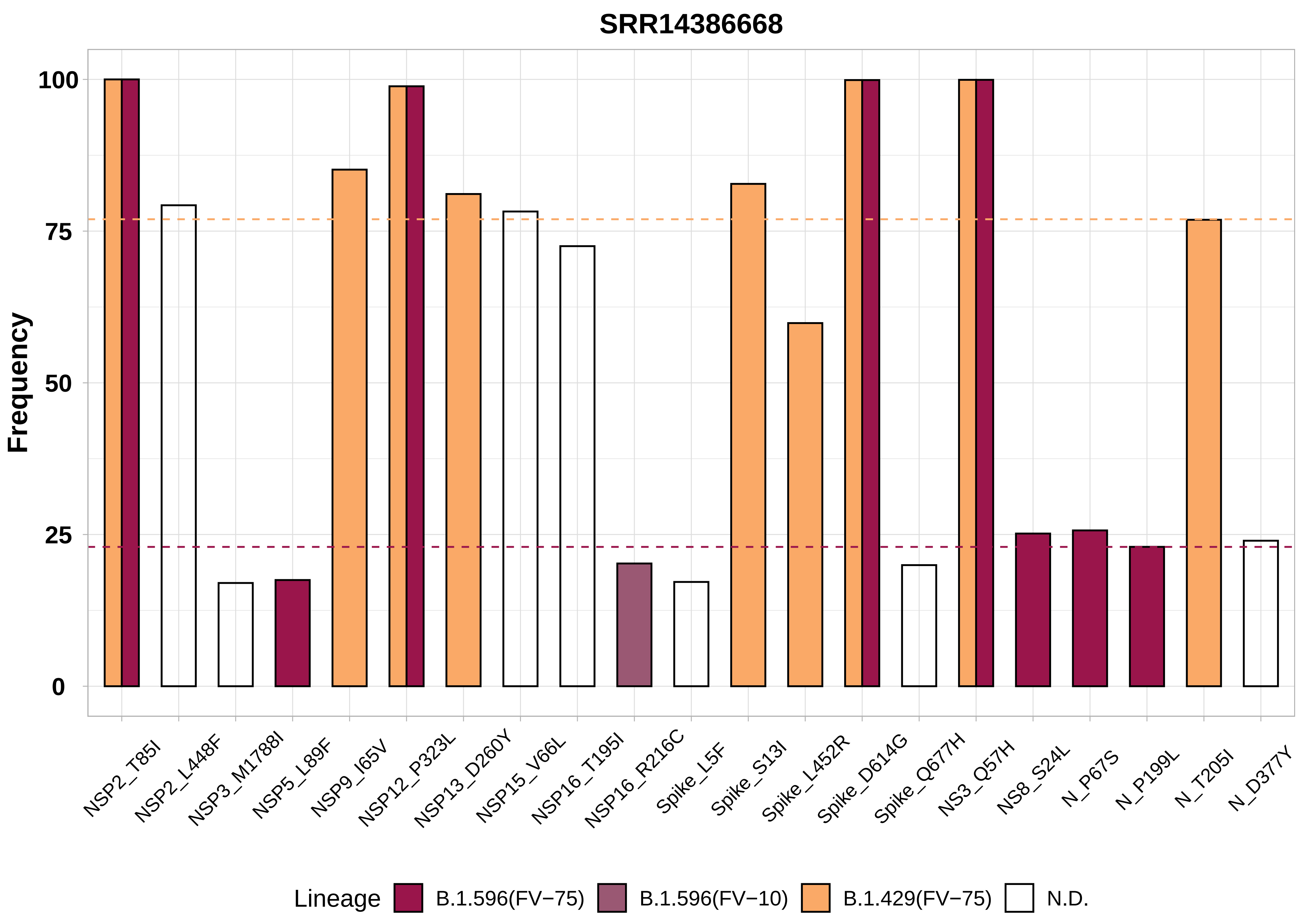

SRR14386668
Frequency
100
75
50
25
0
NSP2_T85I
NSP2_L448F
NSP3_M1788I
NSP5_L89F
NSP9_I65V
NSP12_P323L
NSP13_D260Y
NSP15_V66L
NSP16_T195I
NSP16_R216C
Spike_L5F
Spike_S13I
Spike_L452R
Spike_D614G
Spike_Q677H
NS3_Q57H
NS8_S24L
N_P67S
N_P199L
N_T205I
N_D377Y
Lineage
B.1.596(FV−75)
B.1.596(FV−10)
B.1.429(FV−75)
N.D.

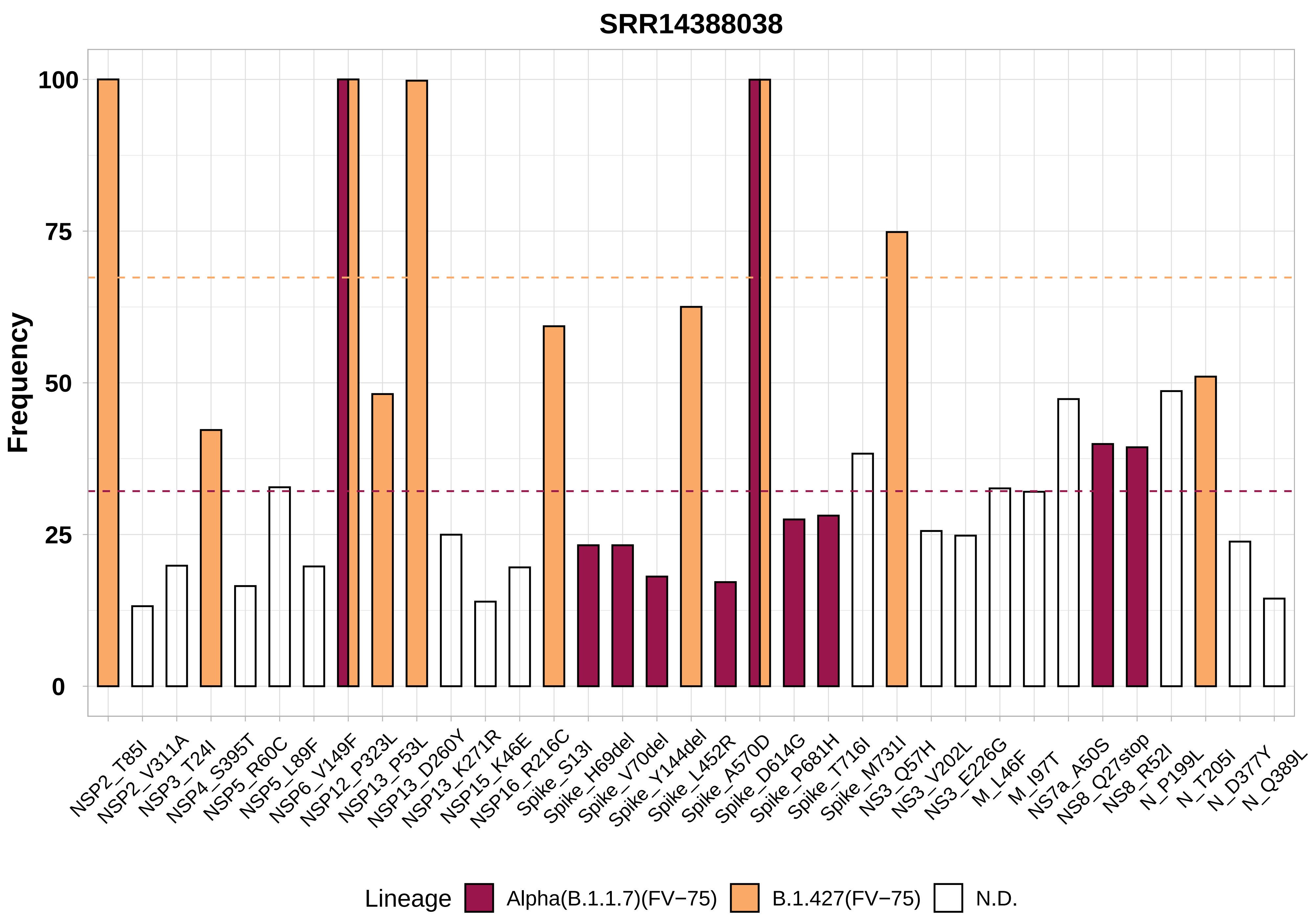

SRR14388038
Frequency
100
75
50
25
0
NSP2_T85I
NSP2_V311A
NSP3_T24I
NSP4_S395T
NSP5_R60C
NSP5_L89F
NSP6_V149F
NSP12_P323L
NSP13_P53L
NSP13_D260Y
NSP13_K271R
NSP15_K46E
NSP16_R216C
Spike_S13I
Spike_H69del
Spike_V70del
Spike_Y144del
Spike_L452R
Spike_A570D
Spike_D614G
Spike_P681H
Spike_T716I
Spike_M731I
NS3_Q57H
NS3_V202L
NS3_E226G
M_L46F
M_I97T
NS7a_A50S
NS8_Q27stop
NS8_R52I
N_P199L
N_T205I
N_D377Y
N_Q389L
Lineage
Alpha(B.1.1.7)(FV−75)
B.1.427(FV−75)
N.D.

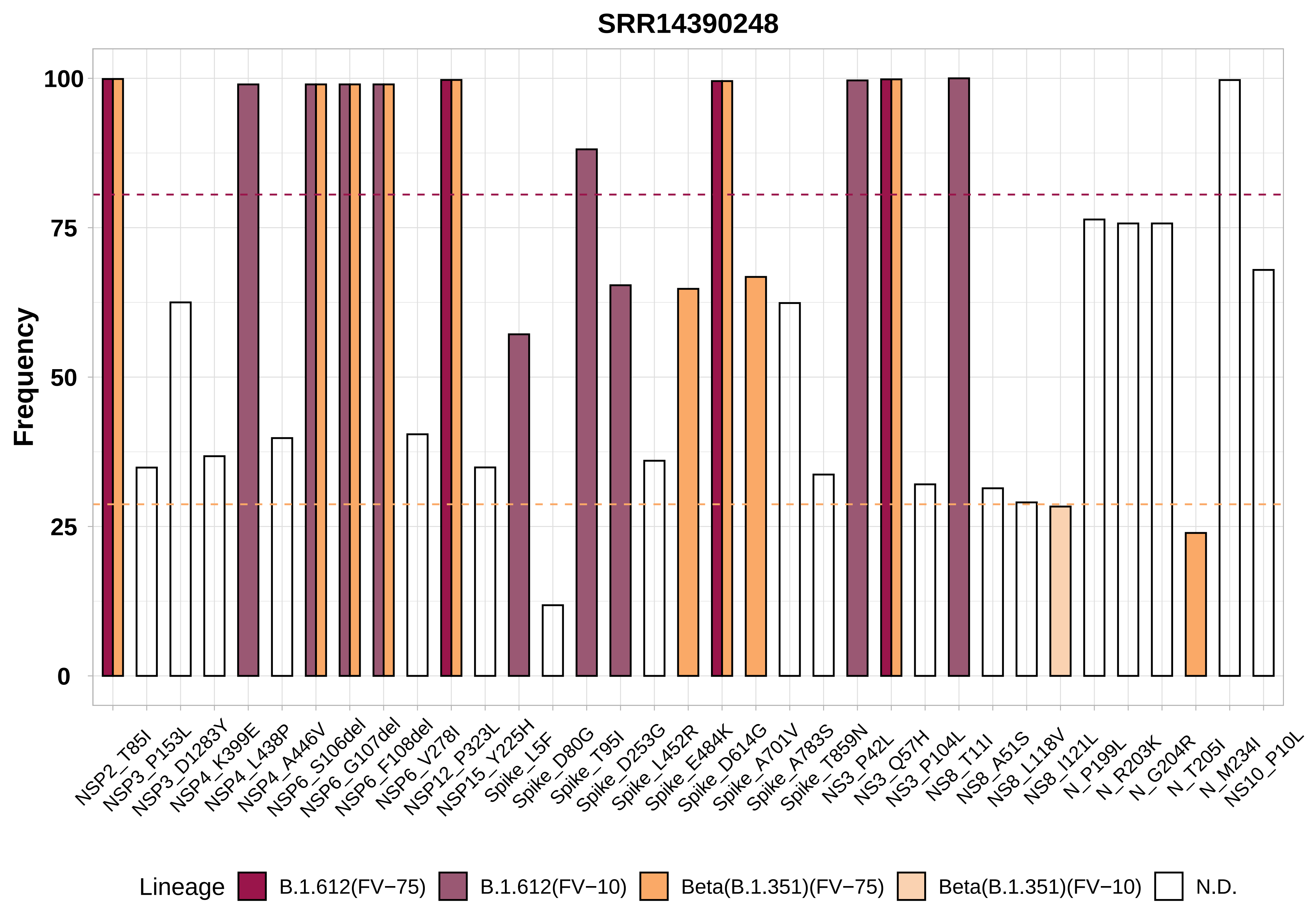

SRR14390248
Frequency
100
75
50
25
0
NSP2_T85I
NSP3_P153L
NSP3_D1283Y
NSP4_K399E
NSP4_L438P
NSP4_A446V
NSP6_S106del
NSP6_G107del
NSP6_F108del
NSP6_V278I
NSP12_P323L
NSP15_Y225H
Spike_L5F
Spike_D80G
Spike_T95I
Spike_D253G
Spike_L452R
Spike_E484K
Spike_D614G
Spike_A701V
Spike_A783S
Spike_T859N
NS3_P42L
NS3_Q57H
NS3_P104L
NS8_T11I
NS8_A51S
NS8_L118V
NS8_I121L
N_P199L
N_R203K
N_G204R
N_T205I
N_M234I
NS10_P10L
Lineage
B.1.612(FV−75)
B.1.612(FV−10)
Beta(B.1.351)(FV−75)
Beta(B.1.351)(FV−10)
N.D.

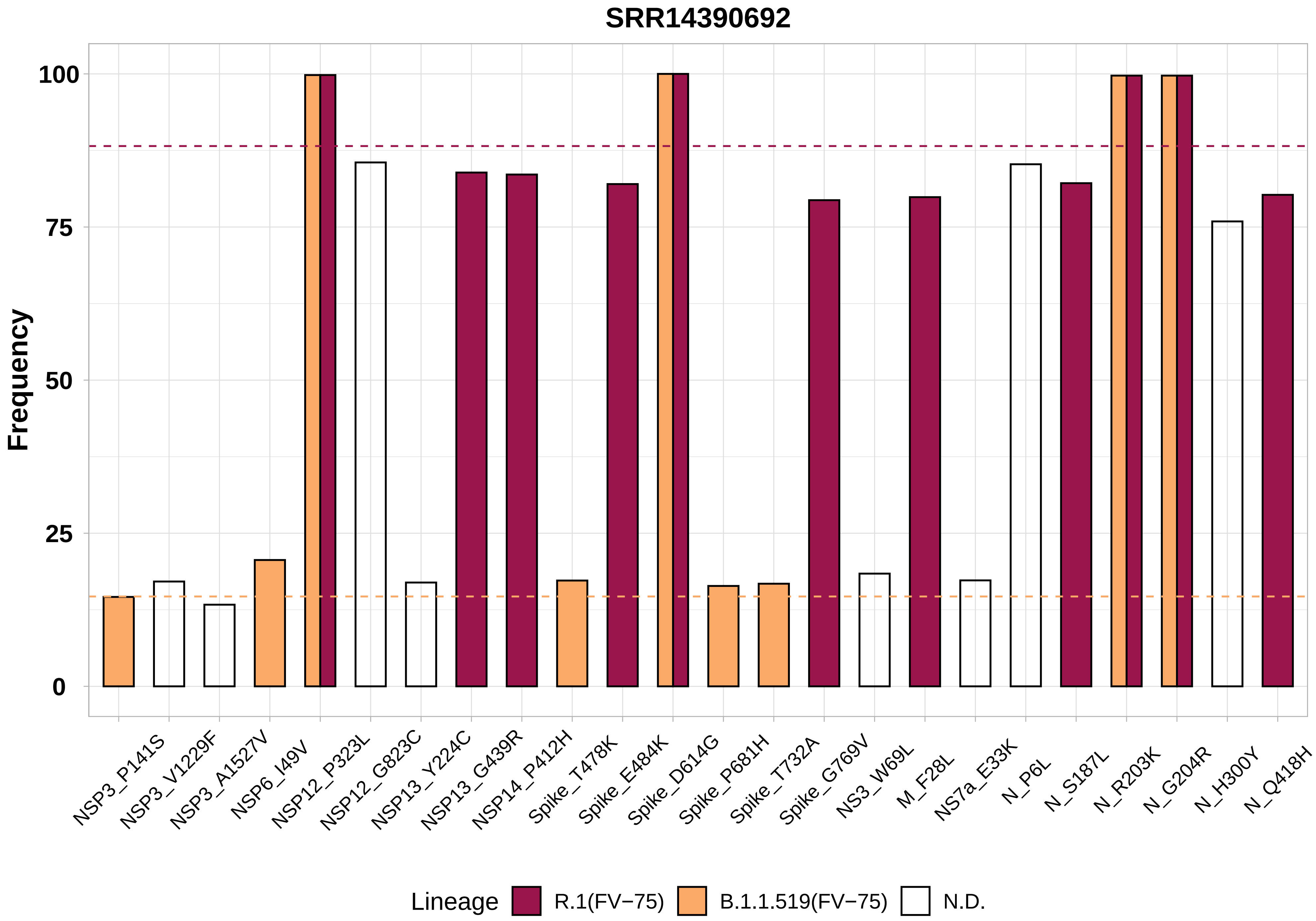

SRR14390692
Frequency
100
75
50
25
0
NSP3_P141S
NSP3_V1229F
NSP3_A1527V
NSP6_I49V
NSP12_P323L
NSP12_G823C
NSP13_Y224C
NSP13_G439R
NSP14_P412H
Spike_T478K
Spike_E484K
Spike_D614G
Spike_P681H
Spike_T732A
Spike_G769V
NS3_W69L
M_F28L
NS7a_E33K
N_P6L
N_S187L
N_R203K
N_G204R
N_H300Y
N_Q418H
Lineage
R.1(FV−75)
B.1.1.519(FV−75)
N.D.

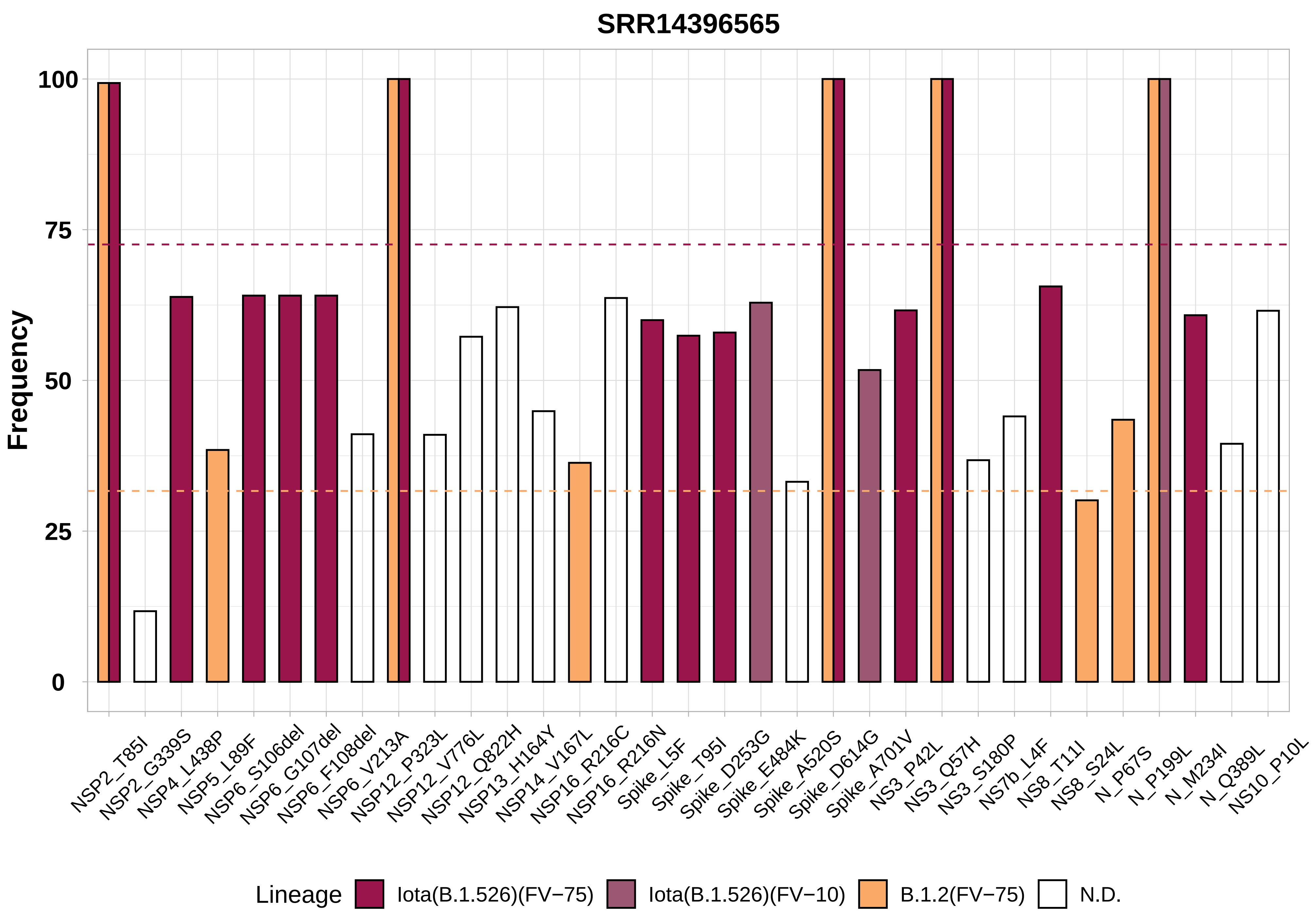

SRR14396565
Frequency
100
75
50
25
0
NSP2_T85I
NSP2_G339S
NSP4_L438P
NSP5_L89F
NSP6_S106del
NSP6_G107del
NSP6_F108del
NSP6_V213A
NSP12_P323L
NSP12_V776L
NSP12_Q822H
NSP13_H164Y
NSP14_V167L
NSP16_R216C
NSP16_R216N
Spike_L5F
Spike_T95I
Spike_D253G
Spike_E484K
Spike_A520S
Spike_D614G
Spike_A701V
NS3_P42L
NS3_Q57H
NS3_S180P
NS7b_L4F
NS8_T11I
NS8_S24L
N_P67S
N_P199L
N_M234I
N_Q389L
NS10_P10L
Lineage
Iota(B.1.526)(FV−75)
Iota(B.1.526)(FV−10)
B.1.2(FV−75)
N.D.

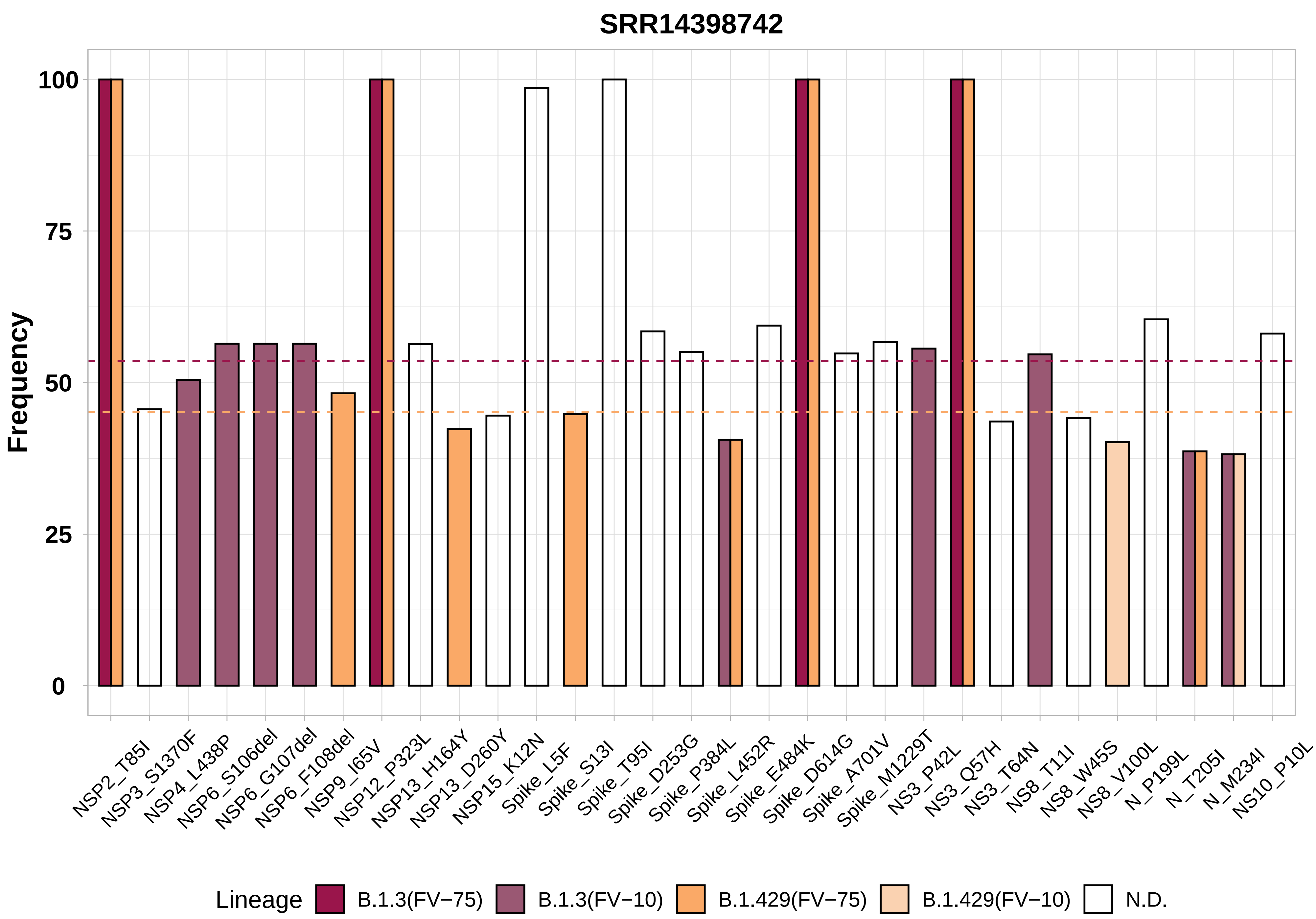

SRR14398742
Frequency
100
75
50
25
0
NSP2_T85I
NSP3_S1370F
NSP4_L438P
NSP6_S106del
NSP6_G107del
NSP6_F108del
NSP9_I65V
NSP12_P323L
NSP13_H164Y
NSP13_D260Y
NSP15_K12N
Spike_L5F
Spike_S13I
Spike_T95I
Spike_D253G
Spike_P384L
Spike_L452R
Spike_E484K
Spike_D614G
Spike_A701V
Spike_M1229T
NS3_P42L
NS3_Q57H
NS3_T64N
NS8_T11I
NS8_W45S
NS8_V100L
N_P199L
N_T205I
N_M234I
NS10_P10L
Lineage
B.1.3(FV−75)
B.1.3(FV−10)
B.1.429(FV−75)
B.1.429(FV−10)
N.D.

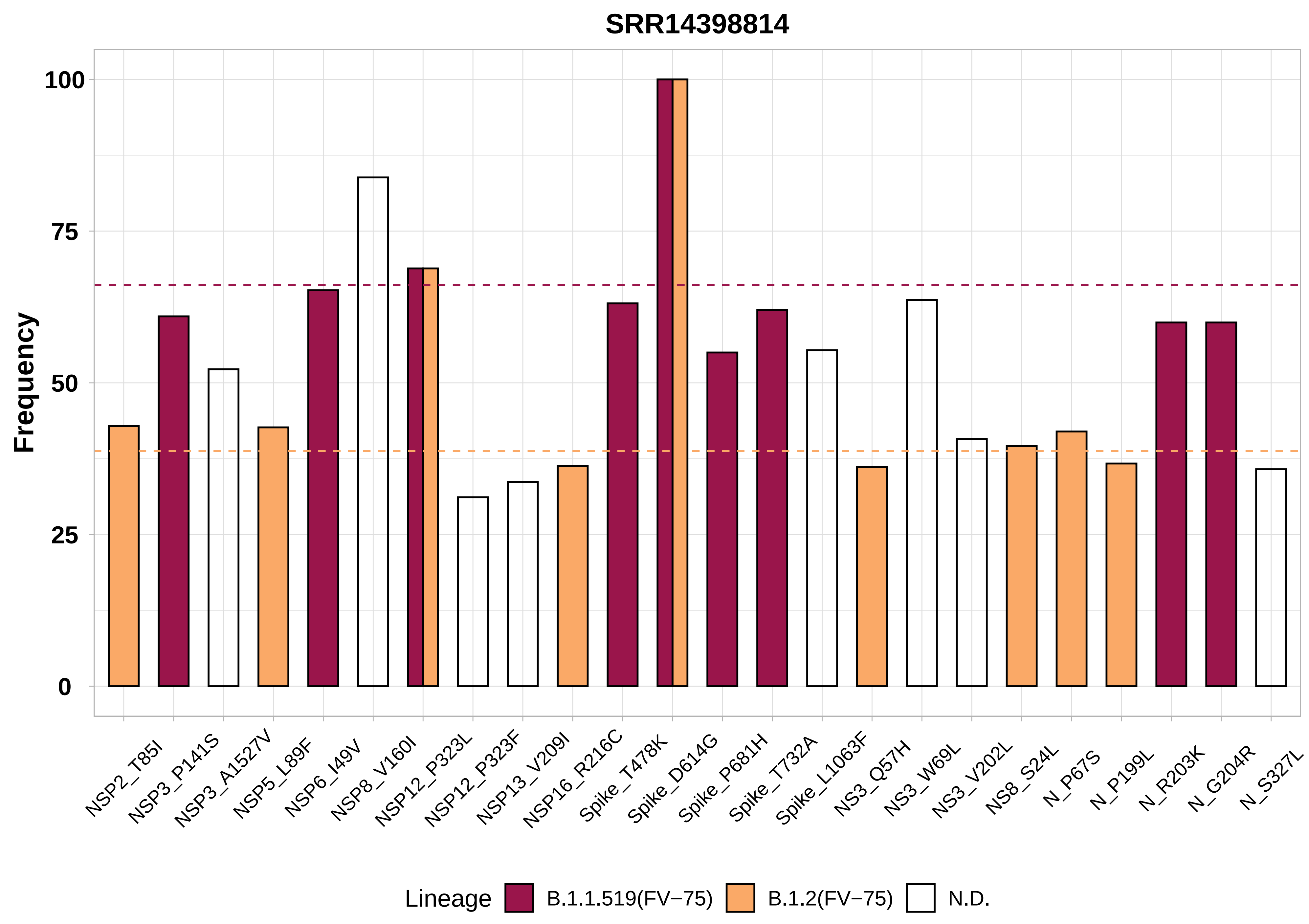

SRR14398814
Frequency
100
75
50
25
0
NSP2_T85I
NSP3_P141S
NSP3_A1527V
NSP5_L89F
NSP6_I49V
NSP8_V160I
NSP12_P323L
NSP12_P323F
NSP13_V209I
NSP16_R216C
Spike_T478K
Spike_D614G
Spike_P681H
Spike_T732A
Spike_L1063F
NS3_Q57H
NS3_W69L
NS3_V202L
NS8_S24L
N_P67S
N_P199L
N_R203K
N_G204R
N_S327L
Lineage
B.1.1.519(FV−75)
B.1.2(FV−75)
N.D.

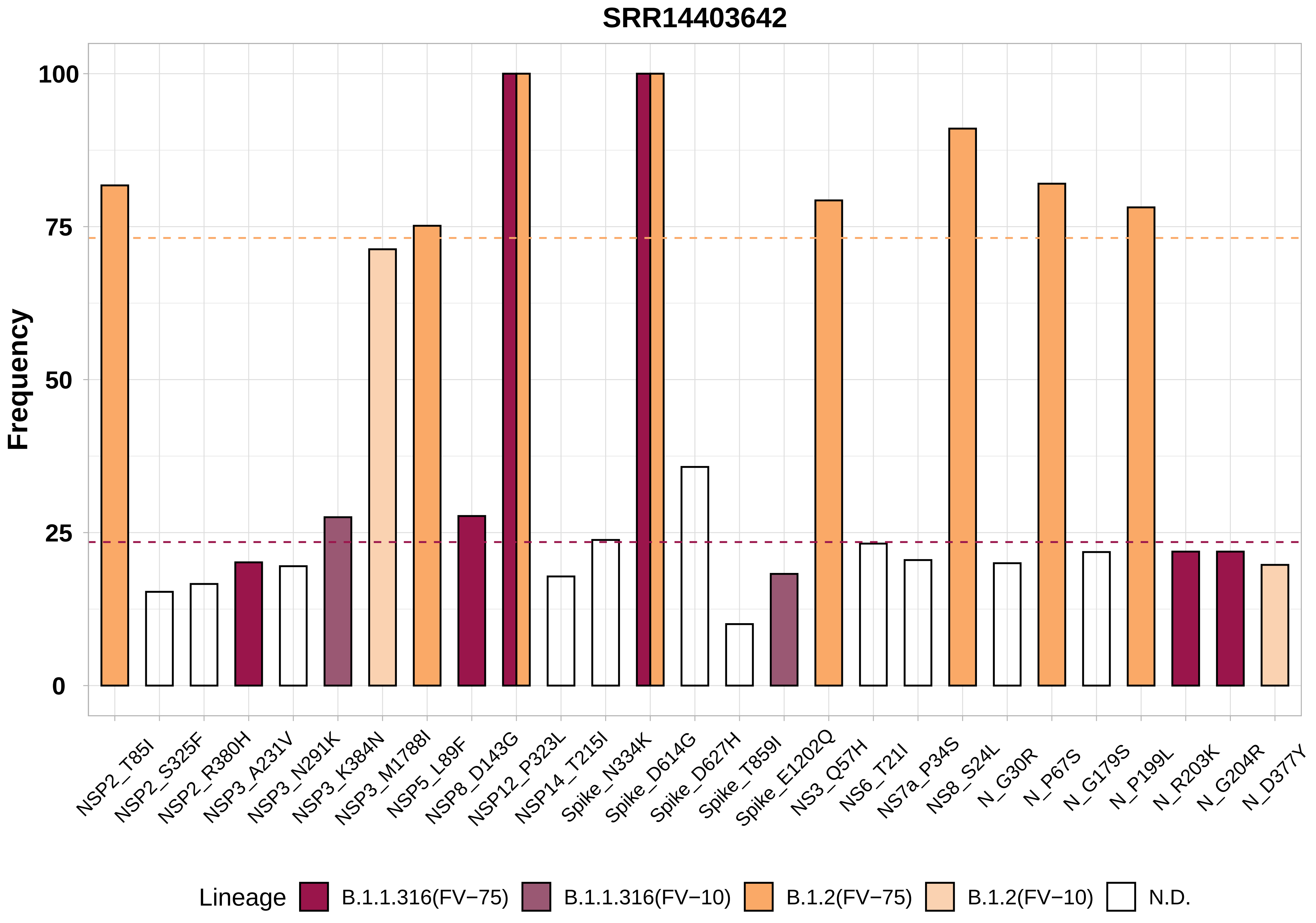

SRR14403642
Frequency
100
75
50
25
0
NSP2_T85I
NSP2_S325F
NSP2_R380H
NSP3_A231V
NSP3_N291K
NSP3_K384N
NSP3_M1788I
NSP5_L89F
NSP8_D143G
NSP12_P323L
NSP14_T215I
Spike_N334K
Spike_D614G
Spike_D627H
Spike_T859I
Spike_E1202Q
NS3_Q57H
NS6_T21I
NS7a_P34S
NS8_S24L
N_G30R
N_P67S
N_G179S
N_P199L
N_R203K
N_G204R
N_D377Y
Lineage
B.1.1.316(FV−75)
B.1.1.316(FV−10)
B.1.2(FV−75)
B.1.2(FV−10)
N.D.

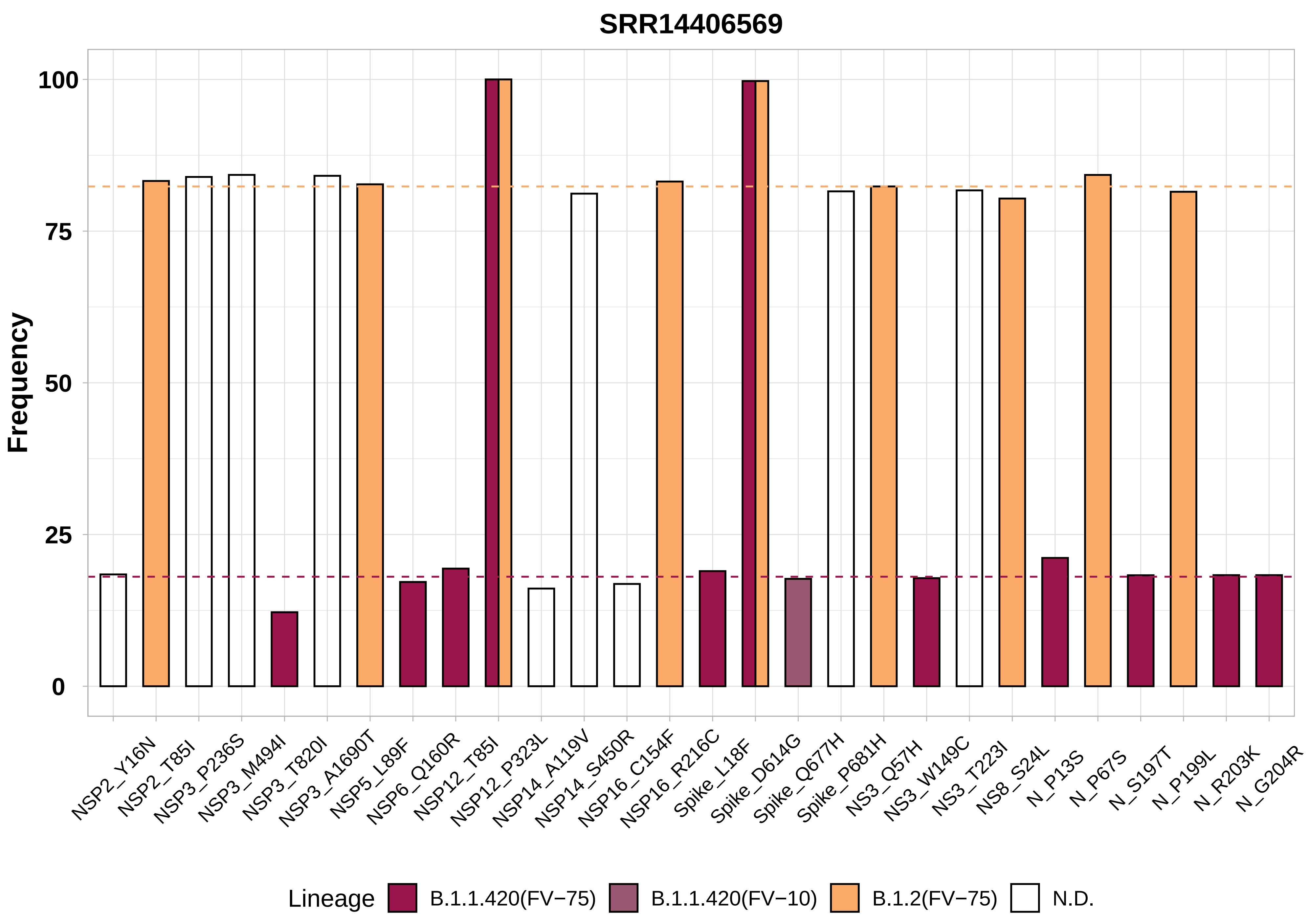

SRR14406569
Frequency
100
75
50
25
0
NSP2_Y16N
NSP2_T85I
NSP3_P236S
NSP3_M494I
NSP3_T820I
NSP3_A1690T
NSP5_L89F
NSP6_Q160R
NSP12_T85I
NSP12_P323L
NSP14_A119V
NSP14_S450R
NSP16_C154F
NSP16_R216C
Spike_L18F
Spike_D614G
Spike_Q677H
Spike_P681H
NS3_Q57H
NS3_W149C
NS3_T223I
NS8_S24L
N_P13S
N_P67S
N_S197T
N_P199L
N_R203K
N_G204R
Lineage
B.1.1.420(FV−75)
B.1.1.420(FV−10)
B.1.2(FV−75)
N.D.

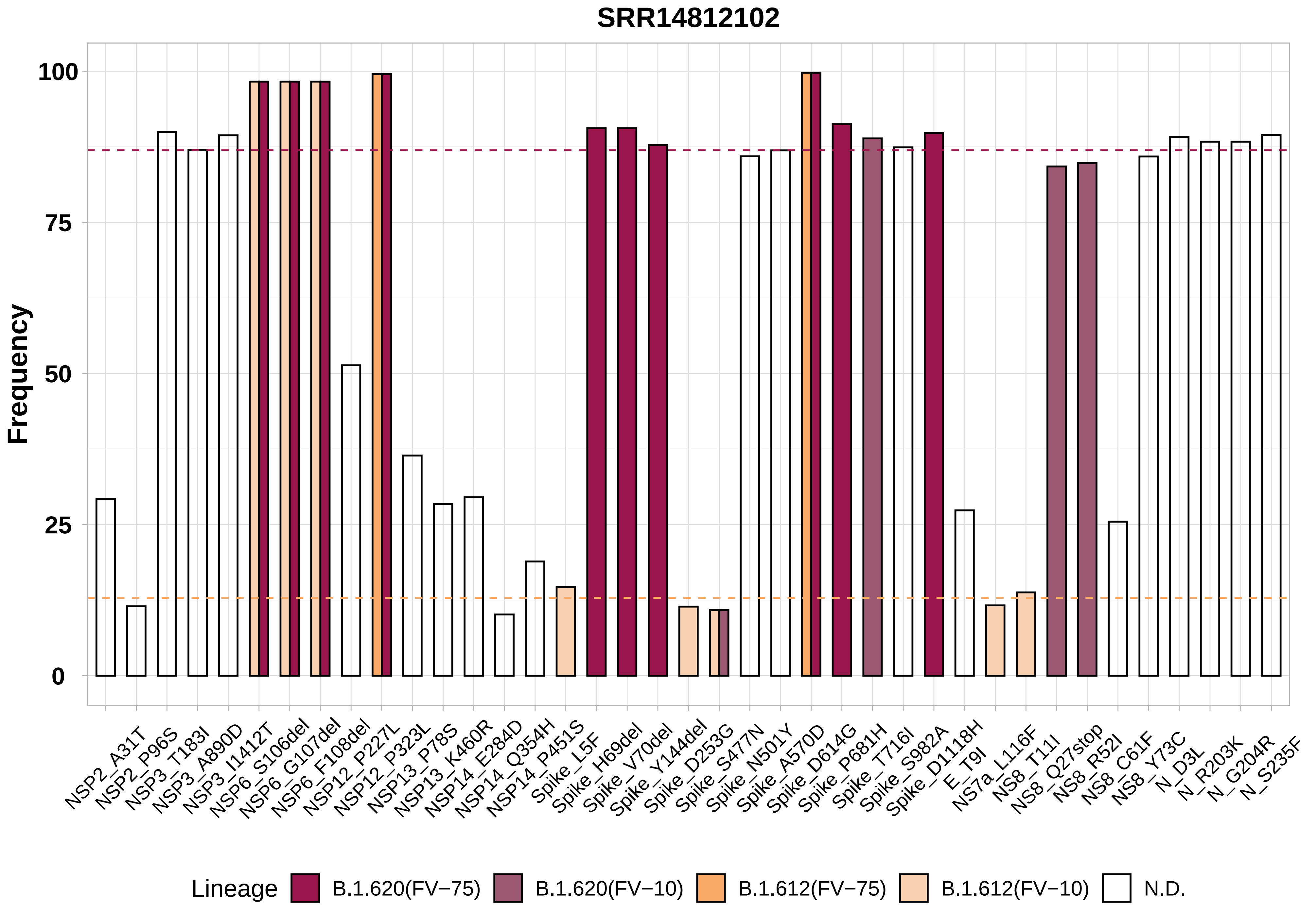
SRR14812102
Frequency
100
75
50
25
0
NSP2_A31T
NSP2_P96S
NSP3_T183I
NSP3_A890D
NSP3_I1412T
NSP6_S106del
NSP6_G107del
NSP6_F108del
NSP12_P227L
NSP12_P323L
NSP13_P78S
NSP13_K460R
NSP14_E284D
NSP14_Q354H
NSP14_P451S
Spike_L5F
Spike_H69del
Spike_V70del
Spike_Y144del
Spike_D253G
Spike_S477N
Spike_N501Y
Spike_A570D
Spike_D614G
Spike_P681H
Spike_T716I
Spike_S982A
Spike_D1118H
E_T9I
NS7a_L116F
NS8_T11I
NS8_Q27stop
NS8_R52I
NS8_C61F
NS8_Y73C
N_D3L
N_R203K
N_G204R
N_S235F
Lineage
B.1.620(FV−75)
B.1.620(FV−10)
B.1.612(FV−75)
B.1.612(FV−10)
N.D.

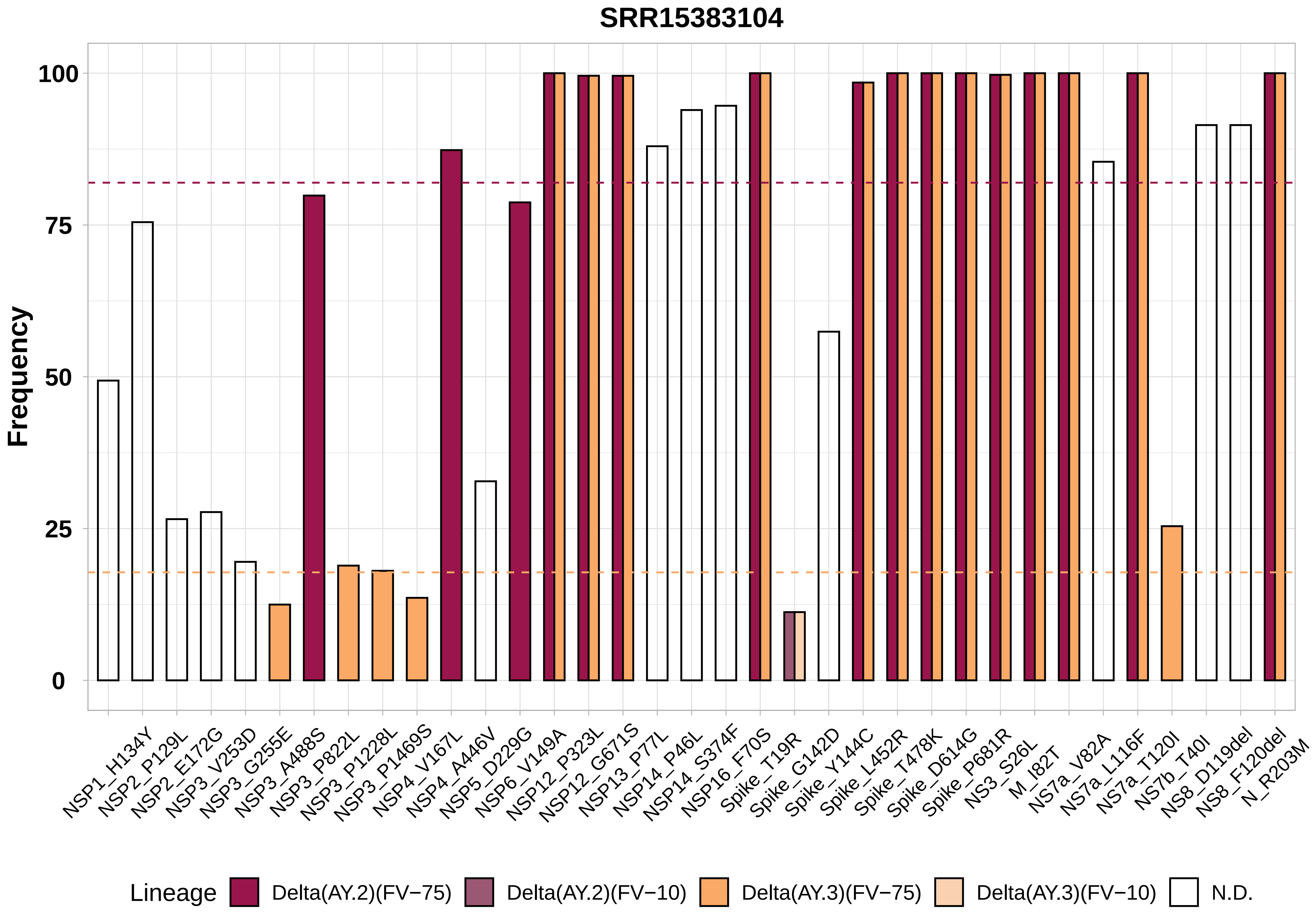

SRR15383104
Frequency
100
75
50
25
0
NSP1_H134Y
NSP2_P129L
NSP2_E172G
NSP3_V253D
NSP3_G255E
NSP3_A488S
NSP3_P822L
NSP3_P1228L
NSP3_P1469S
NSP4_V167L
NSP4_A446V
NSP5_D229G
NSP6_V149A
NSP12_P323L
NSP12_G671S
NSP13_P77L
NSP14_P46L
NSP14_S374F
NSP16_F70S
Spike_T19R
Spike_G142D
Spike_Y144C
Spike_L452R
Spike_T478K
Spike_D614G
Spike_P681R
NS3_S26L
M_I82T
NS7a_V82A
NS7a_L116F
NS7a_T120I
NS7b_T40I
NS8_D119del
NS8_F120del
N_R203M
Lineage
Delta(AY.2)(FV−75)
Delta(AY.2)(FV−10)
Delta(AY.3)(FV−75)
Delta(AY.3)(FV−10)
N.D.

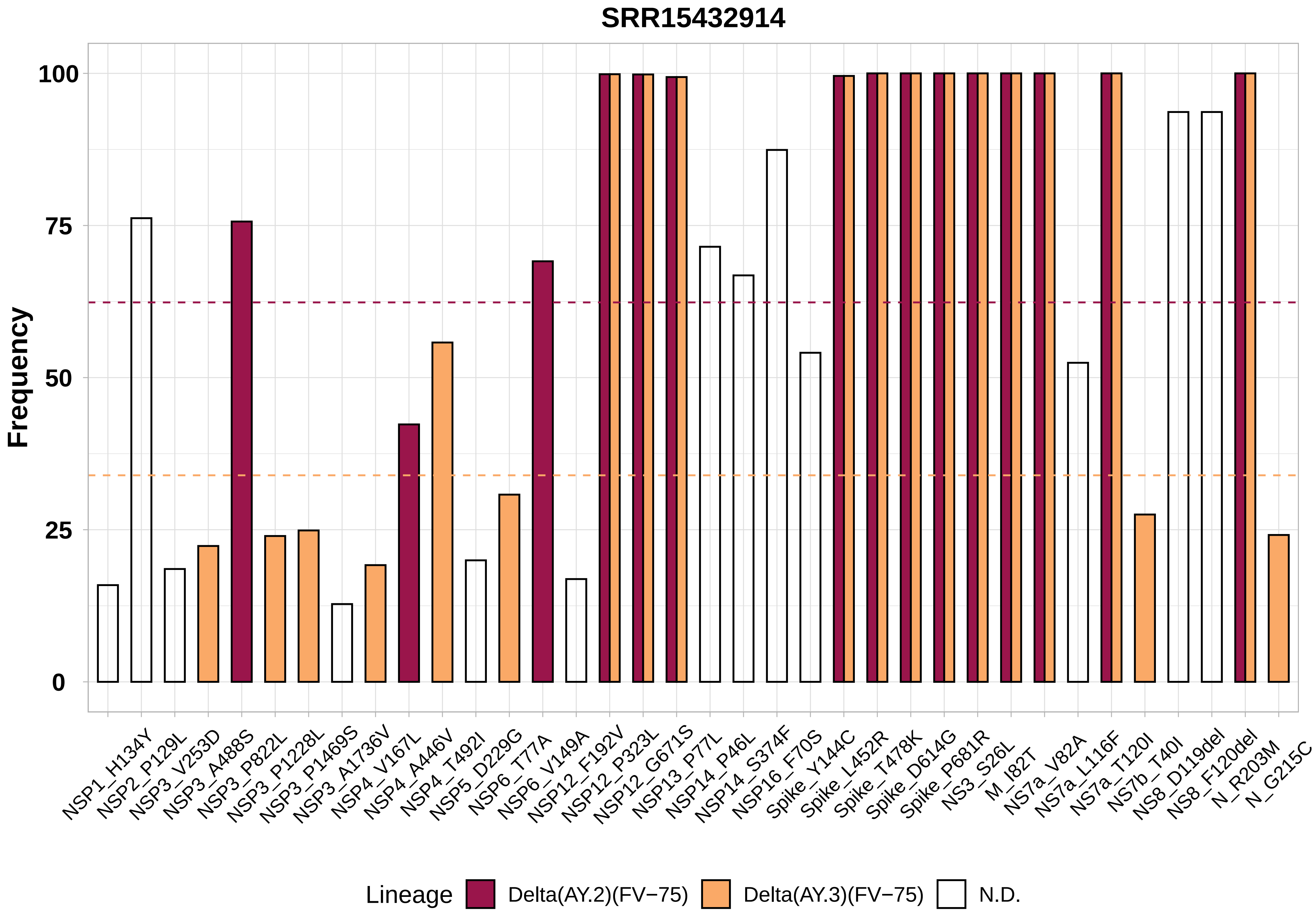
SRR15432914
Frequency
100
75
50
25
0
NSP1_H134Y
NSP2_P129L
NSP3_V253D
NSP3_A488S
NSP3_P822L
NSP3_P1228L
NSP3_P1469S
NSP3_A1736V
NSP4_V167L
NSP4_A446V
NSP4_T492I
NSP5_D229G
NSP6_T77A
NSP6_V149A
NSP12_F192V
NSP12_P323L
NSP12_G671S
NSP13_P77L
NSP14_P46L
NSP14_S374F
NSP16_F70S
Spike_Y144C
Spike_L452R
Spike_T478K
Spike_D614G
Spike_P681R
NS3_S26L
M_I82T
NS7a_V82A
NS7a_L116F
NS7a_T120I
NS7b_T40I
NS8_D119del
NS8_F120del
N_R203M
N_G215C
Lineage
Delta(AY.2)(FV−75)
Delta(AY.3)(FV−75)
N.D.

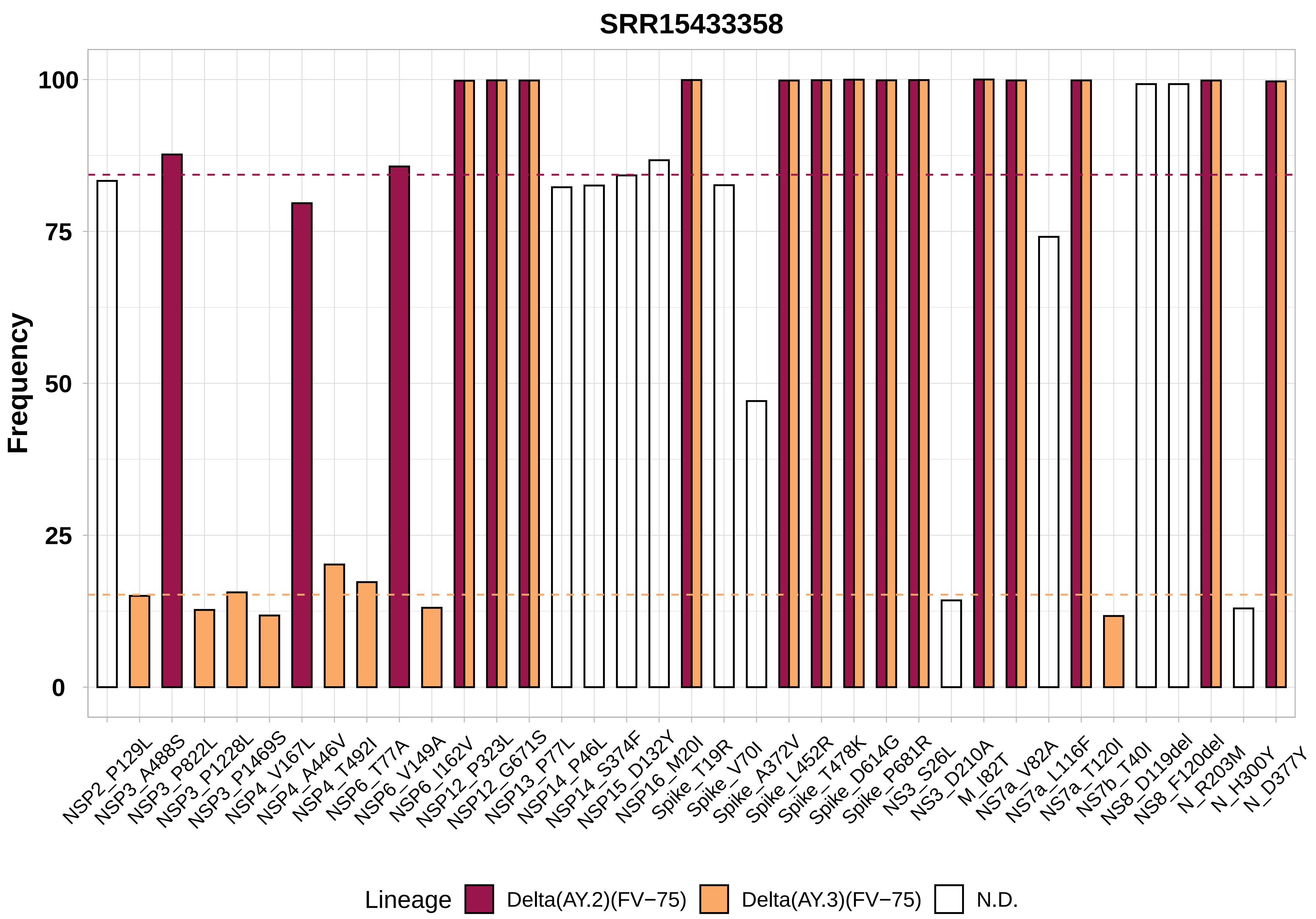

SRR15433358
Frequency
100
75
50
25
0
NSP2_P129L
NSP3_A488S
NSP3_P822L
NSP3_P1228L
NSP3_P1469S
NSP4_V167L
NSP4_A446V
NSP4_T492I
NSP6_T77A
NSP6_V149A
NSP6_I162V
NSP12_P323L
NSP12_G671S
NSP13_P77L
NSP14_P46L
NSP14_S374F
NSP15_D132Y
NSP16_M20I
Spike_T19R
Spike_V70I
Spike_A372V
Spike_L452R
Spike_T478K
Spike_D614G
Spike_P681R
NS3_S26L
NS3_D210A
M_I82T
NS7a_V82A
NS7a_L116F
NS7a_T120I
NS7b_T40I
NS8_D119del
NS8_F120del
N_R203M
N_H300Y
N_D377Y
Lineage
Delta(AY.2)(FV−75)
Delta(AY.3)(FV−75)
N.D.

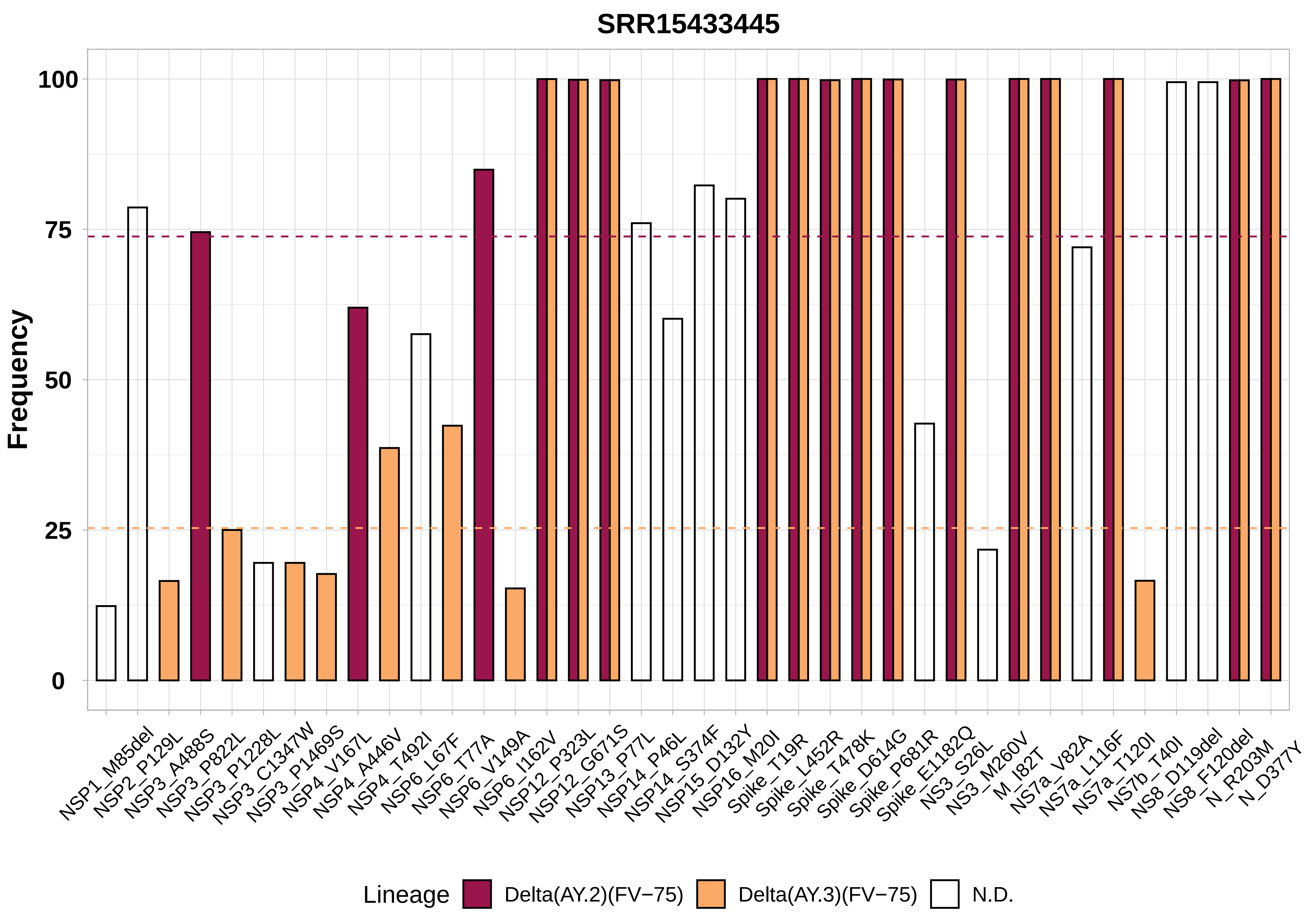

SRR15433445
Frequency
100
75
50
25
0
NSP1_M85del
NSP2_P129L
NSP3_A488S
NSP3_P822L
NSP3_P1228L
NSP3_C1347W
NSP3_P1469S
NSP4_V167L
NSP4_A446V
NSP4_T492I
NSP6_L67F
NSP6_T77A
NSP6_V149A
NSP6_I162V
NSP12_P323L
NSP12_G671S
NSP13_P77L
NSP14_P46L
NSP14_S374F
NSP15_D132Y
NSP16_M20I
Spike_T19R
Spike_L452R
Spike_T478K
Spike_D614G
Spike_P681R
Spike_E1182Q
NS3_S26L
NS3_M260V
M_I82T
NS7a_V82A
NS7a_L116F
NS7a_T120I
NS7b_T40I
NS8_D119del
NS8_F120del
N_R203M
N_D377Y
Lineage
Delta(AY.2)(FV−75)
Delta(AY.3)(FV−75)
N.D.

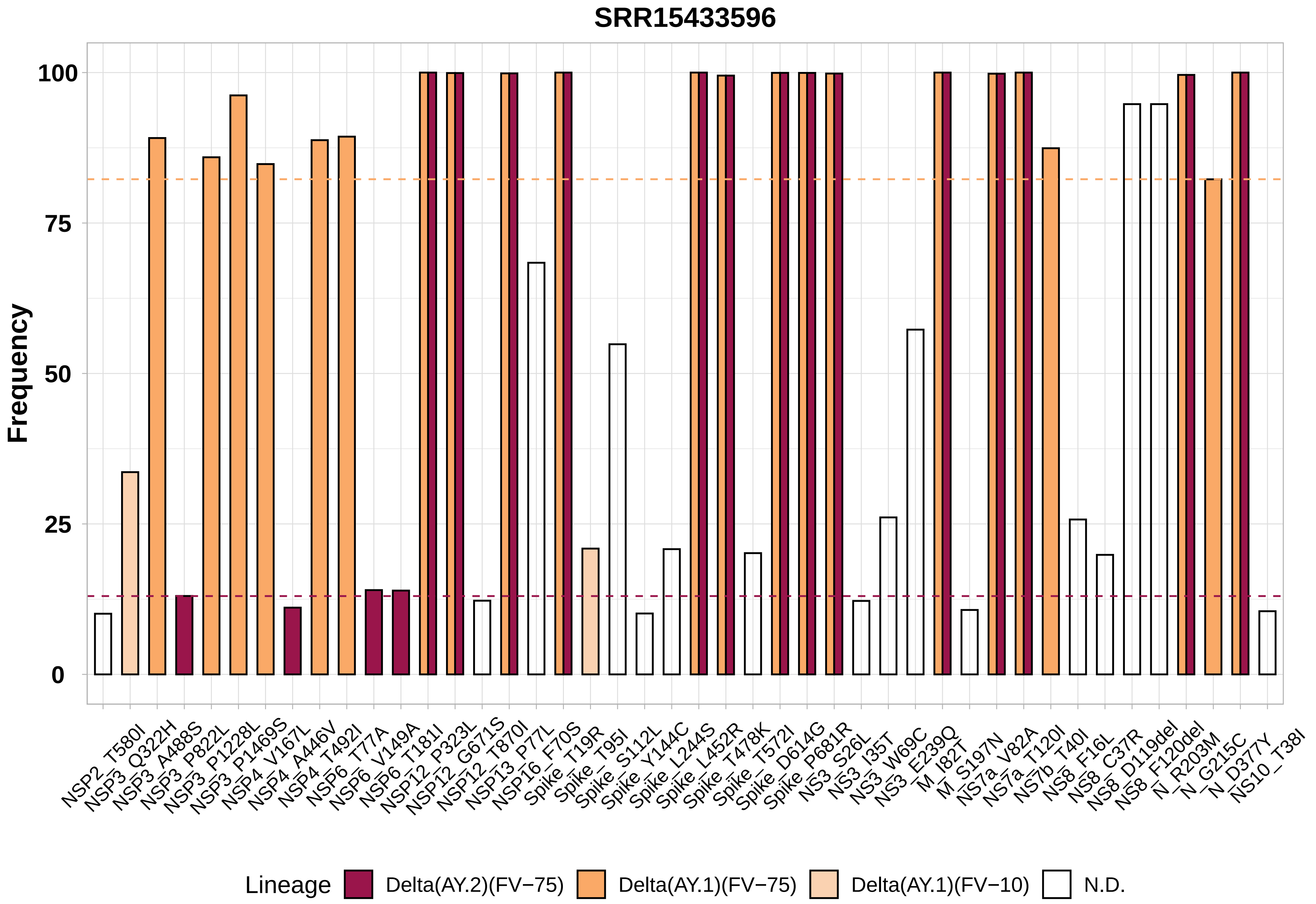

SRR15433596
Frequency
100
75
50
25
0
NSP2_T580I
NSP3_Q322H
NSP3_A488S
NSP3_P822L
NSP3_P1228L
NSP3_P1469S
NSP4_V167L
NSP4_A446V
NSP4_T492I
NSP6_T77A
NSP6_V149A
NSP6_T181I
NSP12_P323L
NSP12_G671S
NSP12_T870I
NSP13_P77L
NSP16_F70S
Spike_T19R
Spike_T95I
Spike_S112L
Spike_Y144C
Spike_L244S
Spike_L452R
Spike_T478K
Spike_T572I
Spike_D614G
Spike_P681R
NS3_S26L
NS3_I35T
NS3_W69C
NS3_E239Q
M_I82T
M_S197N
NS7a_V82A
NS7a_T120I
NS7b_T40I
NS8_F16L
NS8_C37R
NS8_D119del
NS8_F120del
N_R203M
N_G215C
N_D377Y
NS10_T38I
Lineage
Delta(AY.2)(FV−75)
Delta(AY.1)(FV−75)
Delta(AY.1)(FV−10)
N.D.

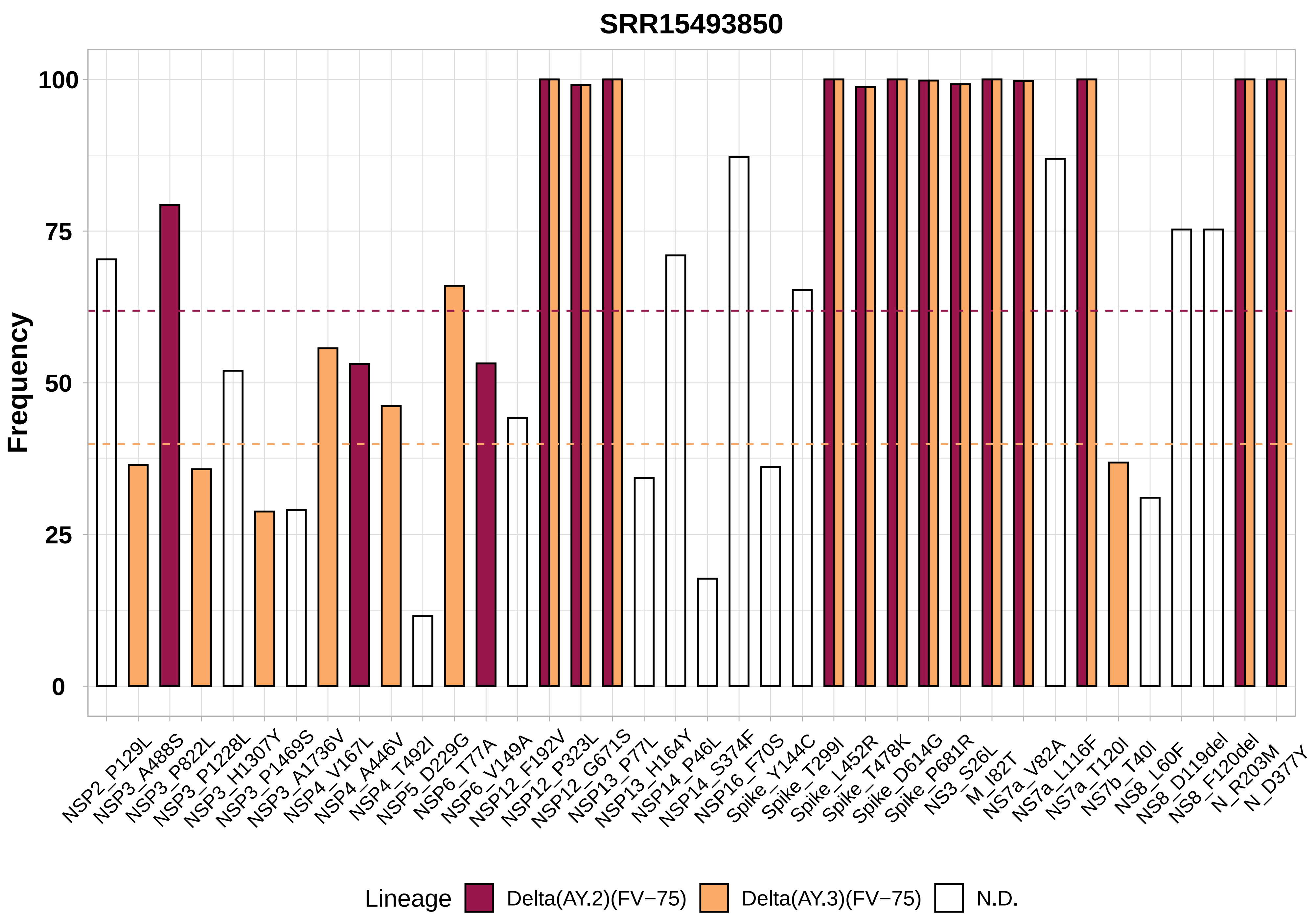

SRR15493850
Frequency
100
75
50
25
0
NSP2_P129L
NSP3_A488S
NSP3_P822L
NSP3_P1228L
NSP3_H1307Y
NSP3_P1469S
NSP3_A1736V
NSP4_V167L
NSP4_A446V
NSP4_T492I
NSP5_D229G
NSP6_T77A
NSP6_V149A
NSP12_F192V
NSP12_P323L
NSP12_G671S
NSP13_P77L
NSP13_H164Y
NSP14_P46L
NSP14_S374F
NSP16_F70S
Spike_Y144C
Spike_T299I
Spike_L452R
Spike_T478K
Spike_D614G
Spike_P681R
NS3_S26L
M_I82T
NS7a_V82A
NS7a_L116F
NS7a_T120I
NS7b_T40I
NS8_L60F
NS8_D119del
NS8_F120del
N_R203M
N_D377Y
Lineage
Delta(AY.2)(FV−75)
Delta(AY.3)(FV−75)
N.D.
